# Establishment of the grass lignin metabolic network during monocot evolution

**DOI:** 10.64898/2026.09.14.751253

**Authors:** Bethany Moore, Yuri Takeda-Kimura, Steven D. Karlen, Jorge El-Azaz, Michael R. McKain, James H. Leebens-Mack, John Ralph, Hiroshi A. Maeda

## Abstract

Plants produce a vast diversity of chemical compounds, yet the evolutionary history of the underlying complex metabolic networks remains poorly understood. Grasses (Poaceae family) possess a unique lignin metabolic network that can utilize both phenylalanine and tyrosine as precursors to synthesize canonical lignin as well as non-canonical acylated and tricin-conjugated lignin subunits. This study traces the evolutionary remodeling and establishment of the grass lignin metabolic network by combining phylogenomics with biochemical and chemical analyses across Poaceae, Poales, and other monocot species. These multidisciplinary comparative analyses reveal that the acylated lignin evolved in commelinids, followed by tyrosine-derived lignin biosynthesis in non-grass graminids through fine-tuning of pathway enzymes and transcriptional factors. Later, biosynthesis and incorporation of the flavonoid tricin into the cell wall took place within the core grasses via the evolution of chrysoeriol 5′-hydroxylase (C5′H) activity within the CYP75B enzyme family. These findings illustrate the emergence of different network modules at distinct times during monocot evolution that became integrated into the complex yet coherent metabolic network underlying the unique lignin chemical diversity of grasses.

## Introduction

Plants produce an extensive array of primary and specialized metabolites that are essential for growth, development, and adaptation. Many of these metabolites are crucial for human society as food, medicine, fuel, and materials. Genes and enzymes involved in diverse plant natural product biosynthesis have been identified through co-expression and synteny analyses aided by machine learning (*1–6*). Investigation of enzyme biochemical evolution revealed that gene duplication and neofunctionalization play a crucial role in the evolution of new enzymes and biosynthetic pathways (*7–13*). Metabolic pathways, although typically depicted as linear, are often nonlinear, highly interconnected, modular, and are captured from specific cells or tissues (*14–17*). Changes in a certain enzyme or pathway may be constrained by or contingent upon other components and modules of a larger metabolic network (*18–20*). Little is known when and how different pathway modules have been recruited during the modification and establishment of metabolic networks. Consequently, the evolutionary history of complex metabolic networks that underlie tremendous chemical and biochemical diversity of plants remains poorly understood.

Lignin biosynthetic pathways provide an intriguing example of a highly complex metabolic network that diversified across various plant groups. Early land plants lacked mechanical reinforcement, necessitating the evolution of lignin for facilitating water transport, supporting upright structures, and establishing a physical barrier, enabling plant expansion across the terrestrial environment (*21*). Lignin, an aromatic polymer, is generally derived from three monolignols, *p*-coumaryl, coniferyl, and sinapyl alcohols, that are integrated into the secondary cell wall after polymerization as *p*-hydroxyphenyl (H), guaiacyl (G), and syringyl (S) units. In most plants, lignin is synthesized from the aromatic amino acid L-phenylalanine, which is derived from the shikimate pathway and synthesized by arogenate dehydratase (ADT) (*22–24*). Phenylalanine ammonia-lyase (PAL) catalyzes the conversion of phenylalanine into cinnamic acid, the committed step of lignin/phenylpropanoid biosynthesis (**Fig. 1A**). Cinnamate 4- hydroxylase (C4H) subsequently converts cinnamic acid to *p*-coumaric acid, which is converted by 4-coumarate:CoA ligase (4CL) to *p-*coumaroyl-CoA (*p*CA-CoA), a common precursor of all three monolignols as well as other phenylpropanoid derivatives (*25–27*). To synthesize G and S monolignols, *p*CA-CoA is converted to caffeoyl-CoA by *p*-hydroxycinnamoyl-CoA:shikimate *p*- hydroxycinnamoyl transferase (HCT), *p*-coumaroyl ester 3-hydroxylase (C3′H), and caffeoyl shikimate esterase (CSE, **Fig. 1A**). The subsequent step, converting the G to S-type precursor, is facilitated by caffeoyl-CoA 3-*O*-methyltransferase (CCoAOMT), ferulate 5-hydroxylase (F5H), and caffeic acid *O*-methyltransferase (COMT). Cinnamoyl-CoA reductase (CCR) and cinnamyl alcohol dehydrogenase (CAD) catalyze the reduction reactions to produce the H, G, and S monolignols. In addition to the production of the “canonical” lignin, some plants have “non-canonical” lignin that incorporate different aromatic monomers into their cell wall (*28–30*).

**Fig. 1.**
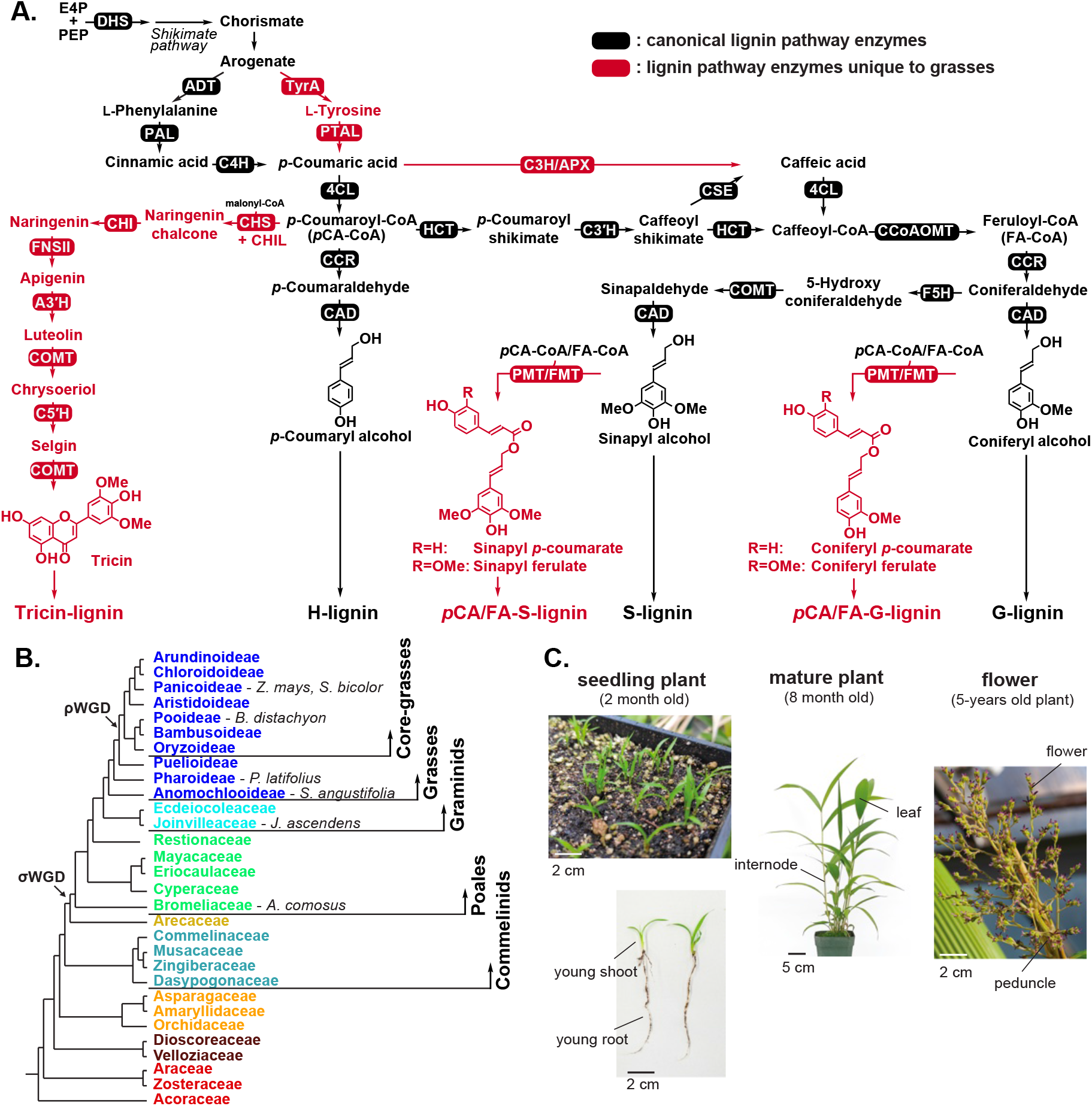
Network analysis of lignin biosynthesis across grasses and non-grass monocot to trace the evolutionary history of grass lignin biosynthesis. **A.** Proposed lignin biosynthetic pathway in grasses. Enzyme abbreviations are shown in **Table S2**. **B.** The family tree of monocot plants showing three ancient whole genome duplication (WGD) events. The colors on the phylogeny denote different taxonomic clades of monocots, corresponding to the molecular gene trees presented in this manuscript. **C.** Tissues used for RNA-seq analysis in *Joinvillea ascendens*

Grasses (Poaceae, Poales) are an agriculturally important and ecologically diverse and dominant plant family (*31–34*). Among many traits that distinguish grasses, such as the spikelet inflorescence structure and starch-rich endosperms (*35–37*), grasses exhibit unique modifications in their lignin metabolic network, as compared to other angiosperms and monocots. Grasses can synthesize lignin through both phenylalanine and L-tyrosine due to the presence of bifunctional phenylalanine/tyrosine ammonia-lyase (PTAL), which catalyzes the direct conversion of tyrosine into *p*-coumaric acid in addition to its PAL activity (*38–40*). This unique dual lignin pathway is supported by altered regulation of the upstream shikimate and tyrosine biosynthetic pathways, because of deregulated 3-deoxy-D-*arabino*heptulosonate-7-phosphate synthase (DHS) and arogenate dehydrogenase (TyrA) enzymes, that efficiently provide both tyrosine and phenylalanine precursors (*41*, *42*) (**Fig. 1A**). Additionally, some grasses, such as *Brachypodium distachyon*, can directly convert *p*-coumaric acid into caffeic acid via the bifunctional enzyme coumarate 3-hydroxylase/ascorbate peroxidase (C3H/APX) (*43*).

Grasses also possess distinct lignin structures. For example, grass lignin is predominantly acylated with *p*-coumarate, forming *p*-coumaroylated lignin (*p*CA lignin) through the enzymatic action of *p*-coumaroylation of the G- and S-monolignols via the enzyme *p*CA-CoA:monolignol transferase (PMT). The monolignol *p*-coumarate conjugates are incorporated into the growing lignin polymer by free-radical coupling processes analogous to those for the monolignols themselves (*29*, *44*, *45*) (**Fig. 1A**). The *p*CA lignin has been identified in other commelinid monocots including the Zingiberales, Arecales, and Poales orders (*46*), as well as in eudicots including kenaf (*Hibiscus cannabinus*) (*28*) and Rosales families Cannabaceae, Urticaceae, and Moraceae (*47*, *48*). Although the prevalence of *p*CA lignin is significantly higher, commelinids also produce feruloylated lignin (FA lignin), catalyzed by feruloyl-CoA (FA-CoA):monolignol transferase (FMT) (*49*, *50*) (**Fig. 1A**).

Tricin, a widely distributed flavone, is present in soluble forms, either in its free form or as *O*- glycosides. Tricin is a bioactive polyphenolic compound with remarkable pharmacological activity (*51–54*). Tricin is also incorporated into the lignin structure of grasses (*53–55*) as well as a few other monocots and the eudicot *Medicago sativa* (*55*, *56*). Tricin is synthesized from *p*CA-CoA through the general flavonoid pathway, catalyzed by chalcone synthase (CHS), chalcone isomerase (CHI), and chalcone isomerase-like (CHIL) enzymes (*57*). Within the tricin-specific pathway, the flavone backbone is formed by flavone synthase II (FNSII) (*58*), followed by aromatic ring hydroxylation by apigenin 3′-hydroxylase/chrysoeriol 5ʹ-hydroxylase (A3′H-C5′H) (*59*). Methylation is carried out by COMT, which is also involved in the biosynthesis of the S-type monolignol biosynthesis (*60*, *61*) (**Fig. 1A**). Although the presence of these unique features of grass “specialized” lignin pathways, including PTAL, PMT and FMT, and the tricin biosynthetic enzymes, has been extensively documented (*43*, *49*, *56*), the mechanisms underlying the diversification and reconfiguring of these metabolic networks during the evolution of grasses and other closely related monocots remain unknown (**Fig. 1B**).

Numerous genomes of grasses and monocots have been sequenced due to the agricultural importance of these plant species, including cereals such as wheat, rice, and maize (*62–65*). Additionally genomes of banana, pineapple, and palm have been sequenced (*66–68*) (**Fig. 1B**). Genome sequences for various species sister to core grasses, such as *Streptochaeta angustifolia* (Family Streptochaeteae) (*36*) and *Pharus latifolius* (*69*), have recently become available. Non-grass graminids, *Joinvillea ascendens* and *Ecdeiocolea monostachya* have also had their genomes sequenced (*40*). These abundant genomic resources facilitate comparative phylogenomic analyses, enabling the dissection of complex traits uniquely evolved within this plant lineage. They also allow for comparisons of the impacts of the *rho* (ρ) whole genome duplication (WGD) event, which occurred just before the emergence of grasses, on these traits (*70*).

In this study, we trace the evolutionary history of the grass lignin network using a combination of co-expression, copy number, phylogenetic, synteny, chemical, and biochemical analyses across monocots, including grasses and closely related species, particularly *Joinvillea ascendens,* which represents the sister lineage to all grasses (**Fig. 1C**). The obtained data collectively reveal the evolution of different enzymes and network modules at distinct stages of monocot evolution and their integration into the complex yet coherent metabolic network of grass lignin biosynthesis. The study underscores the potential to leverage a growing number of plant genomes, in conjunction with network, chemical, and biochemical analyses, to unravel the metabolic network evolution underlying plant chemical diversity.

## Results

### Key lignin enzyme families expand or contract in grasses and monocots

Gene copy number contributes to genetic and phenotypic variation across species, where duplicated genes provide a primary genetic material for the evolution of new functions (*71*). High turnover of gene content within a gene family may indicate functional divergence across species (*71*, *72*). In this study, we determined the copy numbers of enzymes and transcription factors potentially involved in lignin biosynthesis across 44 green plants (including 15 monocots) using orthogroups for gene family membership. The analysis was subsequently expanded using 33 monocot species (see **Methods**, **tables S1** and **S2** for green plant and monocot analyses, respectively). Consistent with a prior study (*27*), many enzymes in the lignin pathway emerged or expanded in land plants after divergence from green algae, including orthogroups of C4H, 4CL, CCR, CAD, COMT, and flavonoid tricin-related cytochrome P450 (CYP450) enzymes A3′H- C5′H and FNSII. The PAL orthogroup expanded in both green algae and land plants (**fig. S1**). Other enzymes involved in later steps of G and S lignin synthesis expanded with gymnosperms and angiosperms (*i.e.*, CCoAOMT) or emerged with angiosperms (*i.e.*, F5H, **fig. S1**).

Within the monocot clade, certain enzymes exhibited alterations in their copy number relative to the outgroup *Amborella trichopoda* (**Fig. 2**). The BAHD acyltransferases involved in *p*CA and FA lignin synthesis (*i.e.*, PMT, FMT) (*44*, *49*) and hemicellulose modification (*73*) expanded in the Asparagales, Zingiberales (Musaceae), Arecales (Arecaceae), and Poales orders, which coincide with the observed cell wall acylation (*46*, *49*). Additionally, PALs underwent a second and significant expansion within grasses relative to other monocots, particularly within the core grass clade (**Fig. 2**). The CYP93G family containing FNSII involved in tricin biosynthesis is also expanded within Poales and some Arecaceae (*e.g.*, palms) relative to other monocots or *Amborella trichopoda* (**Fig. 2**). Although most core lignin enzymes have been largely maintained across land plants (**Fig. 2**; **fig. S1**), the copy number of the CSE enzyme family, involved in the biosynthesis of caffeic acid via the caffeoyl-shikimate shunt (*74*), has decreased or been lost in many grasses (**Fig. 2**) (*75*). This is likely because grasses can “shortcut” caffeic acid synthesis directly from *p*- coumaric acid via a cytosolic peroxidase (*43*). In contrast, CSE is rarely lost outside of the grass family and is present in all other monocots we sampled (including the non-core grasses) and in most eudicots except for *Mimulus guttatus* and *Amaranthus hypochondriacus* (**fig. S1**). The copy number of the lignin pathway genes therefore expanded and contracted within monocots, which may reflect functionally diverged components of the lignin metabolic network.

**Fig. 2.**
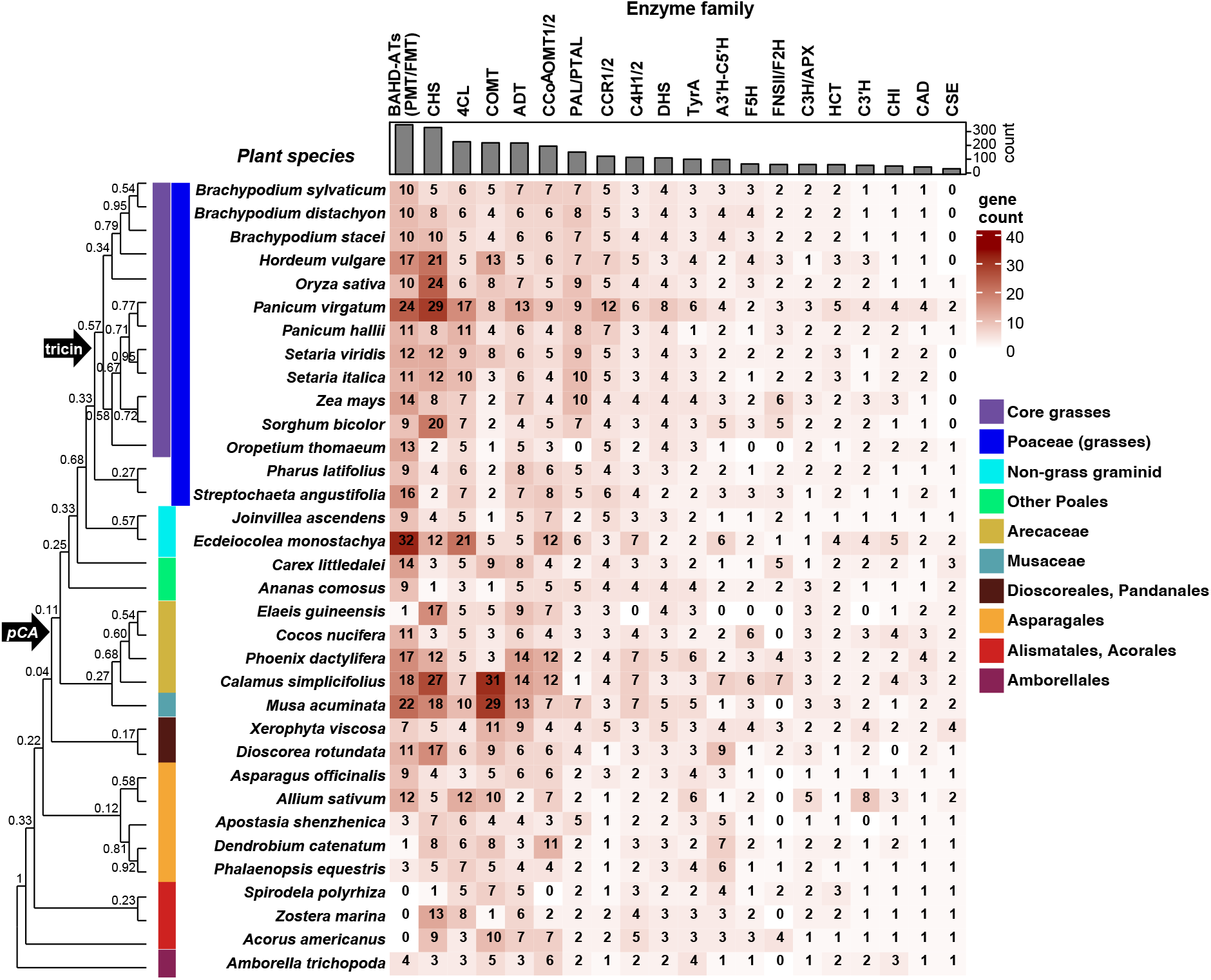
Gene-copy analysis reveals the expansion of PMT, PAL, and FNSII/F2H enzyme families and the loss of CSE in monocots. The heatmap represents the number of gene isoforms of an enzyme family belonging to the lignin pathway (column) in a given species (row). The darker the red color, the more isoforms are present. The bar graph at the top indicates the total number of isoforms for an enzyme family in all species. The color bars on the phylogeny denote different taxonomic clades of monocots as indicated. **Abbreviations:** PMT: *p*-Coumaroyl-CoA monolignol transferase; FMT: feruloyl-CoA monolignol transferase; BAHD: benzyl alcohol *O*-acetyltransferase (BEAT), anthocyanin *O*- hydroxycinnamoyltransferase (AHCT), anthranilate *N*-hydroxycinnamoyl/benzoyltransferase (HCBT), and deacetylvindoline 4-*O*-acetyltransferase (DAT); CHS: chalcone synthase; 4CL: 4- coumarate:CoA ligase; COMT: caffeic acid *O*-methyltransferase; ADT: arogenate dehydratase; CCoAOMT: caffeoyl-CoA 3-*O*-methyltransferase; PAL/PTAL: phenylalanine ammonia lyase/ phenylalanine/tyrosine ammonia-lyase; CCR: cinnamoyl-CoA reductase; C4H: cinnamate 4- hydroxylase; DHS: 3-deoxy-D-arabinoheptulosonate-7-phosphate synthase; TyrA: arogenate dehydrogenase; A3′H-C5′H: apigenin 3′-hydroxylase/chrysoeriol 5′-hydroxylase; F5H: ferulate 5- hydroxylase; FNSII/F2H: flavone synthase II/ flavanone 2-hydroxylase; C3H/APX: coumarate 3- hydroxylase/ascorbate peroxidase; HCT: Hydroxycinnamoyl-CoA shikimate/quinate hydroxycinnamoyl transferase; C3′H: *p*-coumaroyl ester 3-hydroxylase; CHI: chalcone isomerase; CAD: cinnamyl alcohol dehydrogenase; CSE: caffeoyl shikimate esterase.

### Conserved and distinct sets of genes are found in the lignin co-expression network of *Brachypodium* and maize

To understand the recruitment of novel genes/enzymes into the grass lignin metabolic network, we constructed and analyzed the lignin co-expression network in three monocot species, two grasses and one non-grass graminid. For the model grass *B. distachyon*, we used expression data derived from a large sampling of different organs and growth stages (total 43) (*76*). Although a co-expression network was previously constructed for *B. distachyon* using the highest reciprocal rank (HRR) based on Pearson’s correlation (*76*, *77*), we employed Spearman’s rank correlation (*78*, *79*) to capture broader networks that encompass more weakly expressed genes (*e.g.*, some TyrA isoforms, see **Methods**).

To identify genes that co-express with the grass-specific tyrosine-derived lignin network, we initially clustered genes based around the *PTAL* gene (bradi3g49250). *PTAL* frequently co-occurred within the same module as *TyrA1* (bradi1g34790), encoding one of three TyrA isoforms of *B. distachyon* that synthesize tyrosine (*41*), or their modules strongly overlapped (see **Methods, table S3**). We therefore combined them into one *PTAL*-*TyrA1* module. Enrichment analysis revealed that the *PTAL-TyrA1* module had significant overrepresentation in the phenylpropanoid pathway genes (**table S4**) and contained most core lignin pathway genes, including *PAL*, *4CL*, *HCT*, *C3′H*, *CCR*, *CAD*, *COMT*, and *F5H* (**Fig. 3A**, **fig. S2A**, **table S3**). Additionally, *PTAL* and *TyrA1* were co-expressed with grass-specific genes encoding PMT and tricin biosynthetic enzymes (*i.e.*, CHS, CHI, and FNSII), as well as the transcription factors MYB58/63, and secondary cell wall-associated MYB1 (SWAM1), that are known to activate lignin gene expression in angiosperms (*80*, *81*) (**fig. S3**). The *PTAL-TyrA1* module also contained numerous enzymes involved in secondary cell wall polysaccharide biosynthesis (**table S3**) (*82*).

**Fig. 3.**
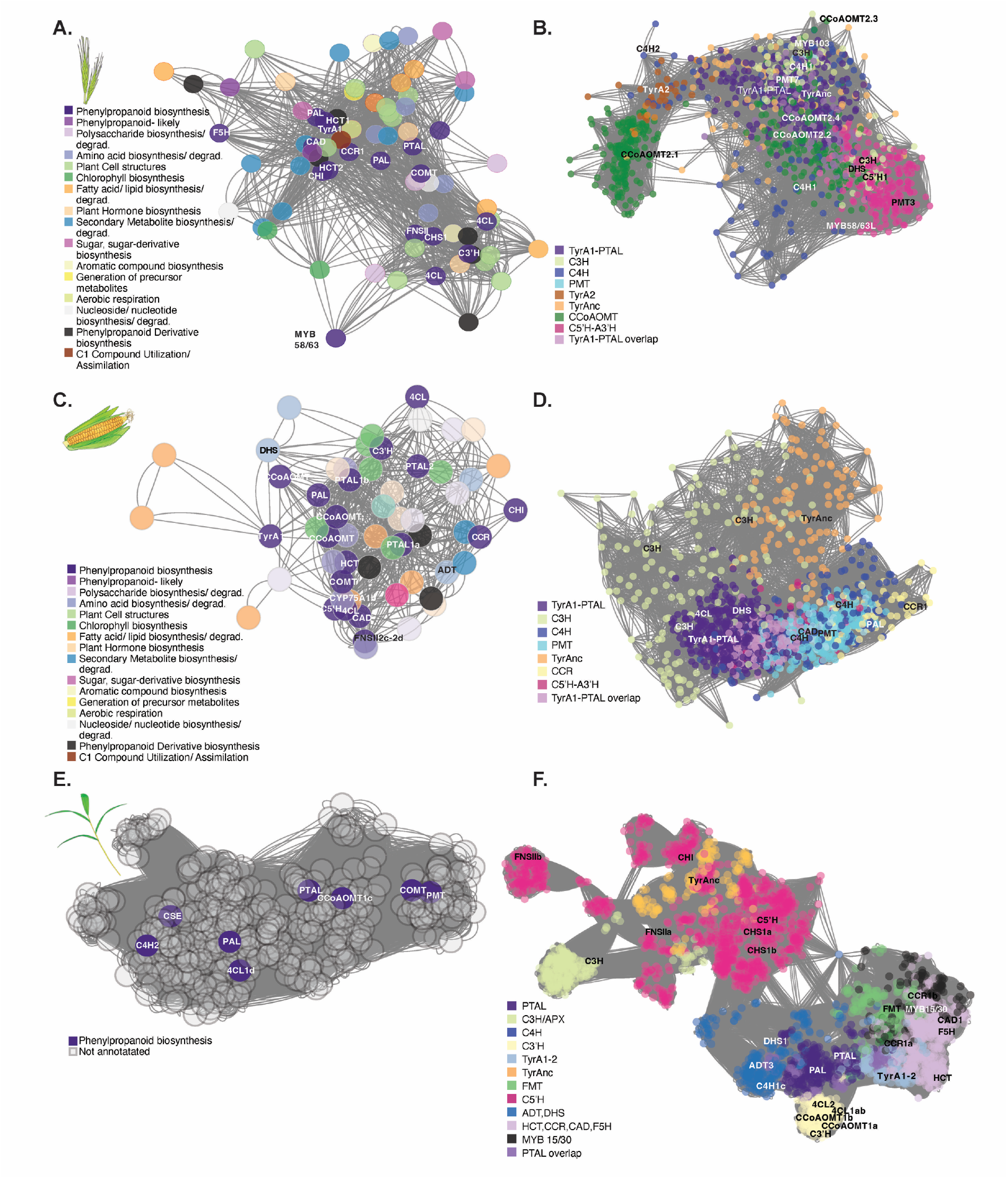
Co-expression of lignin networks change between grasses *Brachypodium distachyon* and *Zea mays,* and the non-grass graminid *Joinvillea ascendens*. **A.** Co-expression of the lignin network in *B. distachyon* based on PTAL and TyrA1 enzymes, where each node is a gene and connections are representative of the distances between genes based on Spearman’s mutual rank. Known lignin enzymes are in dark purple and labeled with their enzyme abbreviation. Other colors correspond to pathways in different categories. Only genes that have an enzyme or pathway annotation are shown. **B.** Overlapping gene-network modules in *B. distachyon*. Genes in modules of the lignin network based around PTAL/TyrA1 are in dark purple and light purple, whereas those in light purple also overlap with other modules. Genes in modules based on other enzymes are in the following colors: C3H/APX: light green, CCoAOMT: dark green, C4H: dark blue, PMT: light blue, A3′H-C5′H: dark pink, TyrA2: orange-brown, TyrAnc: light orange. **C.** Co-expression of the lignin network in *Z. mays* based on PTAL and TyrA1 enzymes. Known lignin enzymes are in dark purple and labeled with their enzyme abbreviation. Other colors correspond to pathways in different categories. Only genes that have an enzyme or pathway annotation are shown. **D.** Overlapping gene network modules in *Z. mays*. The same color scheme from figure 3B is applied here with the exception that the CCR module is in light yellow. **E.** Co-expression of the lignin network in *J. ascendens* based on PTAL and PAL genes. Known lignin enzymes are in dark purple and labeled with their enzyme abbreviation. As we do not have pathway annotations for *J. ascendens*, other genes that are in unknown pathways are shown in gray. **F.** Overlapping gene network modules in *J. ascendens*. Genes in modules of the lignin network based around PTAL/PAL are in dark purple. Genes in modules based on other enzymes are in the following colors: TyrA1-2: light blue, HCT, CCR, CAD, F5H: light purple, C3ʹH: light yellow, FMT: green, MYB15/30: black, ADT and DHS: blue, CHS, CHI, FNSII, A3ʹH-C5ʹH: dark pink, C3H/APX: light green, TyrAnc: light orange. **Abbreviations:** PMT: *p*-Coumaroyl-CoA monolignol transferase; FMT: feruloyl-CoA monolignol transferase; CHS: chalcone synthase; 4CL: 4-coumarate:CoA ligase; COMT: caffeic acid *O*-methyltransferase; ADT: arogenate dehydratase; CCoAOMT: caffeoyl-CoA 3-*O*- methyltransferase; PAL/PTAL: phenylalanine ammonia lyase/ phenylalanine/tyrosine ammonia- lyase; CCR: cinnamoyl-CoA reductase; C4H: cinnamate 4-hydroxylase; DHS: 3-deoxy-D- arabinoheptulosonate-7-phosphate synthase; TyrA: arogenate dehydrogenase; A3ʹH-C5ʹH: apigenin 3ʹ-hydroxylase/chrysoeriol 5ʹ-hydroxylase; F5H: ferulate 5-hydroxylase; FNSII/F2H: flavone synthase II/ flavanone 2-hydroxylase; C3H/APX: coumarate 3-hydroxylase/ascorbate peroxidase; HCT: Hydroxycinnamoyl-CoA shikimate/quinate hydroxycinnamoyl transferase; C3ʹH: *p*-coumarate ester 3-hydroxylase; CHI: chalcone isomerase; CAD: cinnamyl alcohol dehydrogenase; CSE: caffeoyl shikimate esterase.

Genes of two core lignin enzymes, C4H and CCoAOMT, did not co-express with *PTAL* and *TyrA1*, nor did those of other grass-specific enzymes, including A3ʹH-C5ʹH and C3H/APX catalyzing the last step of tricin biosynthesis (*83*) and the “shortcut” synthesis to caffeic acid (*43*), respectively. However, co-expression can vary based on the size and cohesiveness of clusters and different components of a pathway may be regulated and co-express differently (*5*). We analyzed co-expression modules centered around these missing genes and asked whether any of these modules significantly overlapped with the *PTAL-TyrA1* module (see **Methods**). This analysis revealed that modules with *C4H* (bradi2g31510, bradi2g53470) and *CCoAOMT* (bradi3g39380) overlap significantly with the *PTAL-TyrA1* module (**Fig. 3B**, see **table S5** for overlap percentages). Similarly, the modules containing *C3H/APX* (bradi1g16510) and those of *A3′H-C5′H* (bradi4g16560) significantly but distinctly overlapped with the *PTAL-TyrA1* modules (**Fig. 3B, table S5**). The *A3′H-C5′H* modules contained genes encoding C3′H and CHIL, the known transcriptional activators of lignin biosynthesis, MYB42/85 and MYB58/63-like (MYB58/63L, **Fig. 3B**), as well as the specific DHS1b isoform that is insensitive to feedback inhibition and contributes to the efficient production of aromatic amino acid precursors (*41*) (**Fig. 3B**). The modules containing feedback-insensitive, non-canonical TyrA (TyrAnc) (*41*), but not a feedback-sensitive isoform TyrA2, also significantly overlapped with the *PTAL-TyrA1* modules (**Fig. 3B**). Another transcriptional activator of lignin biosynthesis, MYB103 (*80*, *81*), was found in the *C3H/APX* module. The phenylalanine biosynthetic enzyme ADTs, FMT, and the transcription activator involved in secondary cell wall biosynthesis, MYB46/83 (*84*), were, however, still missing from all of these modules. Overall, these module comparisons identify additional isoforms of both core and grass-specific lignin pathway genes that are closely associated with the *PTAL- TyrA1* network.

To ascertain the conservation of the lignin network among grasses, a *Zea mays* co-expression network was constructed using expression data from similar tissue types and developmental stages to those in *B. distachyon* (*85*). Because we used cultivar B73 (*65*) and PH207 (*86*) genomes for the gene expression and phylogenetic mapping, respectively, we identified the reciprocal best match between genes across the two different maize genomes using BLAST similarity scores (see **Methods**). The network built around the three *Z. mays PTALs* (Zm00008a016750, Zm00008a022367, and Zm00008a06867) using the Spearman’s mutual rank confirmed that all three *PTALs* were highly co-expressed with core-lignin pathway genes and the *PTALs* and *TyrA1* co-expression modules highly overlapped with each other also in *Z. mays* (99.5th percentile of the overlap of all modules). As in *B. distachyon*, the *PTAL-TyrA1* module of *Z. mays* included genes of most core lignin pathway enzymes (*e.g.*, PAL, 4CL, HCT, C3′H, CCoAOMT, CCR, CAD, COMT) and grass-specific tricin lignin enzymes (*e.g.*, CHI, FNSII, A3′H-C5′H, COMT) as well as DHS and the MYB transcription activators, MYB46/83, MYB42/85, MYB58/63L, MYB55/61, MYB103, and SWAM1. Again, as in *B. distachyon*, core lignin enzymes C4H and CSE were missing in the *PTAL-TyrA1* network (**Fig. 3C,D, fig. S3B**, **table S6**). Unlike *B. distachyon*, however, the *Z. mays* network lacked the grass-specific *PMT*, *CHS*, and *CHIL* genes but included *CCoAOMT*, *A3′H-C5′H*, as well as *ADT* involved in phenylalanine biosynthesis (**Fig. 3C**,**D**, **table S6**).

Among genes missing in the *Z. mays PTAL-TyrA1* network, *C4H* isoforms *1a* and *1b* were co-expressed in the same module that overlaps significantly with the *PTAL*, but not TyrA1, modules (**table S5**, **Fig. 3D**). The *C4H* module of *Z. mays* includes its immediate up and downstream enzymes, 4CL and PAL. The *PMT* modules significantly overlapped with the *PTAL-TyrA1* modules and contained the core lignin genes *4CL*, *HCT*, *CCoAOMT*, *CAD*, and *MYB58*/63 (**Fig. 3D**, **table S6**). *PMT* was also found in the module clustered around *A3′H-C5′H* (**Fig. 3D**). Unlike in *B. distachyon*, the modules of *TyrAnc* and *C3H/APX* did not overlap with the *PTAL-TyrA1* module (**table S5**), while *F5H* was not found in the *Z. mays* transcriptome dataset used. None of the three *Z. mays C3H/APX* isoforms co-expressed with any lignin pathway genes and instead co-expressed with aerobic respiration, cytokinin hormone biosynthesis, and other salvage/detox pathway genes (**table S6**), suggesting that C3H/APX may function differently between *B. distachyon* and *Z. mays.* These combined findings demonstrated that *PTAL*, *TyrA1*, tricin biosynthetic genes, and *PMT* are co-expressed with core lignin genes in *B. distachyon* and *Z. mays*, highlighting conserved and distinct sets of genes present in the lignin co-expression network of two different grasses.

### *Joinvillea* co-expression analysis reveals a stepwise evolution of the grass lignin network

To elucidate the evolutionary trajectory of the grass lignin co-expression network, we captured the transcriptome of *J. ascendens*, a closely related non-grass graminid that is a direct sister to the entire grass family. Despite previously reported challenges in seed germination and seedling establishment (*87*), our improved germination protocol (see **Methods**) yielded a sufficient number of seedlings for RNA-sequencing across seven different tissue types and developmental stages of *J. ascendens* (**Fig. 1C**). Spearman’s mutual rank was again used to capture lignin co-expression modules using *J. ascendens PAL* and *PTAL* orthologs (Joascv11021328m and Joascv11021323m, respectively) (*40*) as bait genes. Although *PAL* and *PTAL* were not in the same module, their individual modules significantly overlapped (**table S5**). The combined *PAL* and *PTAL* module encompassed the core lignin genes, *C4H*, *4CL*, *CSE*, *CCoAOMT*, and *COMT*, in addition to *PMT* (**Fig. 3E**); the *4CL*, *CCoAOMT*, and *COMT* genes were also co-expressed in the grass *PTAL-TyrA1* networks. The module containing *CCR1a* was significantly overlapped with both *PAL* and *PTAL* modules, whereas modules containing *HCT*, *C3′H*, *CCR1b*, *CAD1*, and *F5H* were adjacent to the PTAL module but only significantly overlapped with CCR1a, not with the PAL or PTAL modules in *J. ascendens* (**Fig. 3F**, **tables S5** and **S7**). The module containing *TyrA1* overlaps significantly with the modules containing *PTAL* and *CCR1a*, but not *PAL*, in *J. ascendens*. Quantitative reverse transcription PCR (qRT-PCR) analyses demonstrate that *JaPTAL* is more highly expressed than *JaPAL* in the internodes of *J. ascendens* (**fig. S4A**). Transcript levels of two *JaTyrA* genes, normalized to *JaUBI10*, showed strong expression in the internodes alongside other lignin pathway genes (**fig. S4B**). These findings substantiate the establishment of a functional tyrosine-derived lignin pathway within this non-grass graminid species.

*J. ascendens* exhibited some distinct features compared to the grass lignin networks, notably the absence of co-expression of the orthologs of tricin biosynthetic genes (*CHS*, *CHIL*, *CHI*, *FNSII*, and *A3′H/C5′H*), *C3H/APX*, and *TyrAnc*, with any of the above lignin modules (**Fig. 3F**, **table S7**). Conversely, two core lignin pathway genes *C4H* and *CSE* were co-expressed in the *PAL/PTAL* module in *J. ascendens* (**Fig. 3E**), but not with *PAL/PTAL* in grasses (**Fig. 3A**,**C**). Unlike in grasses, *FMT* significantly overlapped with *CCR1b* and *CAD1* modules, suggesting the active involvement of *FMT* in lignification in *J. ascendens*. MYB activator genes, including *MYB46/83, MYB42/85, MYB55/61, MYB103*, and *SWAM1*, were in the adjacent module, but *MYB58/63* was absent from all the modules. Instead, *MYB15/30*, which is the ortholog of rice *MYB30* functioning in stress-induced lignification (*88*, *89*), was present in the *FMT* module. These findings imply that different MYB factors were recruited to regulate lignin biosynthesis over the course of grass evolution. Overall, the co-expression analyses of *J. ascendens* revealed that some lignin pathway genes previously considered to be grass-specific (*e.g.*, *PTAL*) are present in the core lignin network of this non-grass graminid, alongside *PMT* that likely functions in broader commelinid monocots (*46*). However, other genes were specifically recruited to (*e.g.*, *MYB58/63*, tricin biosynthetic genes) or potentially lost (*e.g.*, *CSE*) from the lignin network in grasses (**fig. S3**).

### Multiple lignin pathway genes are syntenic across Poales and Poaceae genomes

Syntenic analyses can highlight conserved genetic regions across multiple genomes that can reveal functional relationships (*e.g.*, involvement in the same metabolic pathway) and underlying genetic mechanisms (*e.g.*, gene duplication) (*90*, *91*). Here we examined syntenic regions for genes encoding critical core (*i.e.*, PAL/PTAL, TyrA) and specialized (*i.e.*, PMT, A3′H-C5′H, FNSII) lignin enzymes, across the genomes of multiple Poales and grass species, including *Ananas comosus, J. ascendens, Streptochaeta angustifolia, Pharus latifolius, B. distachyon, Sorghum bicolor,* and *Z. mays* (see **Methods**). The *TyrAnc* gene was syntenic across *J. ascendens* and grasses (*42*), whereas *TyrA1* and/or *TyrA2* was syntenic only within grasses and underwent a tandem duplication within core grasses (**fig. S5A**). The *PMT* (**fig. S5B**) and *PAL/PTAL* (**Fig. 4**) genes were also syntenic across all Poales species analyzed, consistent with our recent report that the *PTAL* gene resulted from a tandem duplication of the *PAL* gene in the common ancestor of *J. ascendens* and grasses (*40*). Additionally, the *PAL*-*PTAL* pair duplicated by the whole-genome duplication (ρWGD) within grasses and then again by the *Z. mays*-specific WGD (*92*) (**Fig. 4**). Thus, the *PAL* copy number expansion within grasses (**Fig. 2**) took place via repeated tandem duplications in one specific synteny block, whereas the other block maintained the “ancestral” state with one *PTAL* and one *PAL* (**Fig. 4**). Finally, although *A3ʹH-C5ʹH* did not have any significant synteny, *FNSII* involved in tricin biosynthesis was syntenic within grasses, including *P. latifolius*, but not with the non-grass Poales *J. ascendens* and *A. comosus* (**fig. S6**). Some lignin pathway genes (*e.g.*, *PAL/PTAL*, *TyrAnc*, *PMT*) therefore have conserved synteny between *J. ascendens* and grasses, whereas others like *TyrA1* and *FNSII* were only in grass-specific synteny, consistent with the lack of their tight co-expression with the *PAL/PTAL* genes in *J. ascendens* (**Fig. 3E**,**F**).

**Fig. 4.**
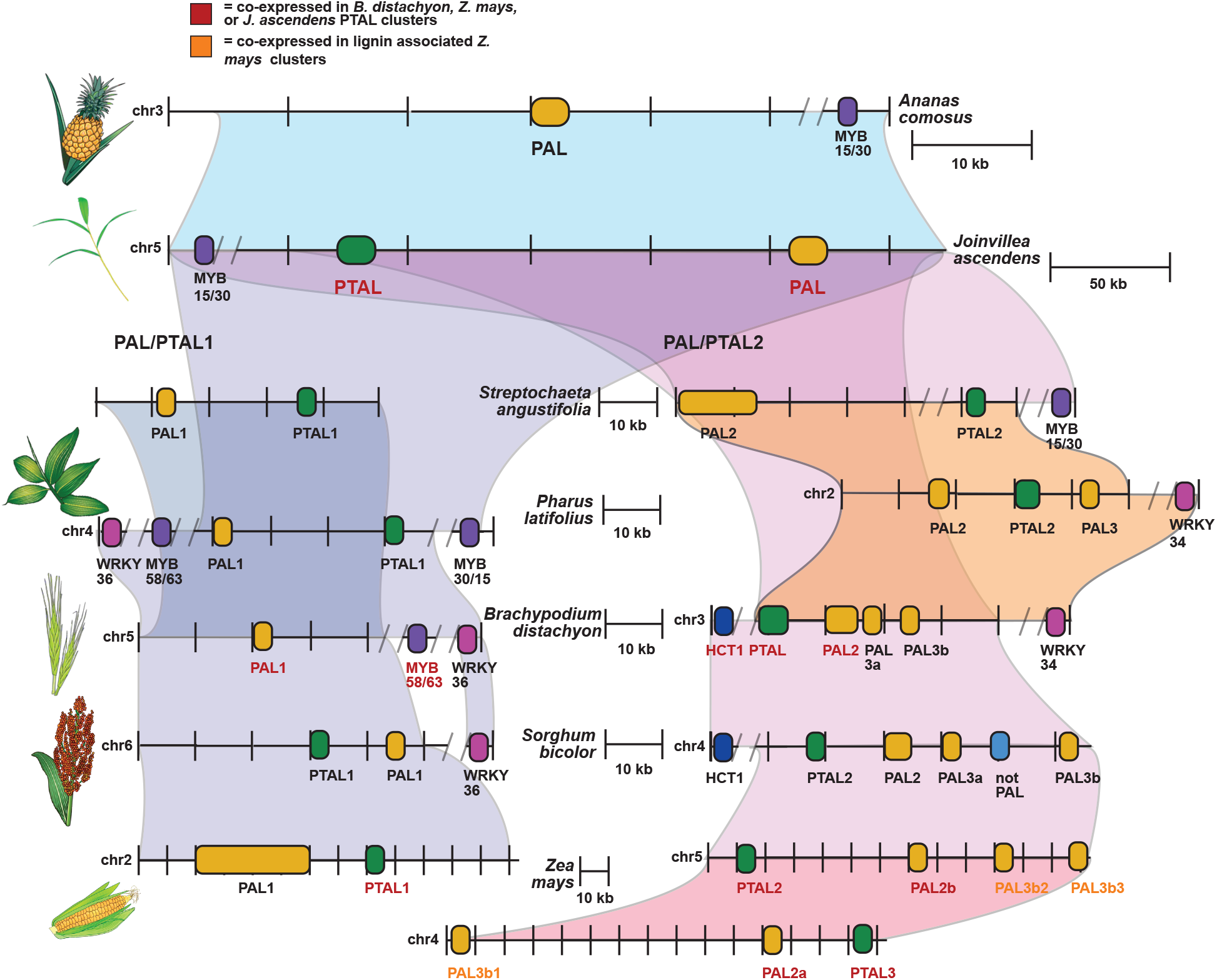
PTAL, resulting from a tandem duplication of PAL before the _ρ_WGD, is syntenic with MYB and WRKY transcriptional factors. Synteny analysis, performed using the CoGe database, revealed a tandem duplication of PAL/PTAL in *J. ascendens*, followed by a duplication of the PAL/PTAL pair within grasses coinciding with the ρWGD in grasses. Color scheme: PAL genes are yellow, PTAL genes are green, MYB transcriptional factors (MYB TF) are dark purple, WRKY transcriptional factors (WRKY TF) are dark pink. Relative distance on each chromosome is given by the scale for each species. Genes labeled in red indicate they are co-expressed with other lignin pathway genes in that species. PAL/PTAL: phenylalanine ammonia lyase/ phenylalanine/tyrosine ammonia-lyase; MYB: myeloblastosis.

We also found other lignin related genes in the same syntenic blocks as *PAL/PTAL* across species (**Fig. 4**). In *B. distachyon* and *S. bicolor,* their *PAL/PTAL* syntenic blocks contained *HCT*, which was found in the *B. distachyon* co-expression network. The *PAL/PTAL* syntenic blocks of two grasses, *P. latifolius* and *B. distachyon*, contained the gene of MYB58/63 that activates lignin production in *A. thaliana* and grasses (Rao and Dixon, 2018) and was co-expressed with lignin pathway genes in *B. distachyon* and *Z. mays* (**Fig. 3**, **fig. S3**). In contrast, *MYB15/30* (*88*, *89*) was found in the *PAL/PTAL* syntenic block in the non-grass Poales, *J. ascendens* and *A. comosus*, as well as the non-core grasses, *S. angustifolia* and *P. latifolius*. This coincides with the presence of MYB15/30, rather than MYB58/63, within the lignin co-expression network of *J. ascendens* (**Fig. 3F)**. WRKY transcriptional factors 34 and/or 36, which negatively regulate the lignin pathway (*93*, *94*), were also syntenic in the grasses *P. latifolius*, *B. distachyon*, and *S. bicolor*, but not in non-grasses, *J. ascendens* and *A. comosus* (**Fig. 4**). Overall, lignin pathway genes found in the co-expression network in both grasses and non-grasses frequently exhibited a syntenic relationship. Furthermore, genes that were found in the lignin co-expression network of *J. ascendens* but were absent in those of grasses—and *vice versa* (*e.g.*, FNSII, MYBs)—were often not syntenic, highlighting the presence or absence of genes in certain genomic regions may contribute to fine-tuning of the lignin metabolic network.

### Co-expression combined with phylogenies reveals recruitments of specific orthologous isoforms at different stages of lignin metabolic network evolution

To further investigate potential divergence and recruitment history of specific enzyme orthologs involved in the lignin metabolic network, gene family phylogenies were constructed and compared with their co-expression memberships. We first built green plant trees to identify the monocot clades (gray arrows in the inserts of **figs. S7** to **S27**) and then generated the extensive monocot gene trees (**Fig. 5**; **figs. S7** to **S27**). Consistent with taxonomic relationships, orthologs of non-grass graminids, including *J. ascendens* and *Ecdeiocolea monostachya*, nested to the clade of grass orthologs, including ones from non-core grasses, *S. angustifolia* and *P. latifolius*. Gene duplication events were detected for many genes near the base of Poaceae, potentially through ρWGD (*e.g.*, *PMT*, *PAL, DHS1*, *ADT1a*, **Fig. 5, figs. S8, S14, S15, S25**). Notably, *4CL1*, *CCoAOMT2*, and *CCR2* underwent multiple rounds of gene duplications before grass evolution within Poales (**figs. S8, S23, S24**), while extensive gene losses were noted within core grasses for CSE (**fig. S26**) in agreement with the copy number and co-expression analyses (**Figs. 2** and **3**).

**Fig. 5.**
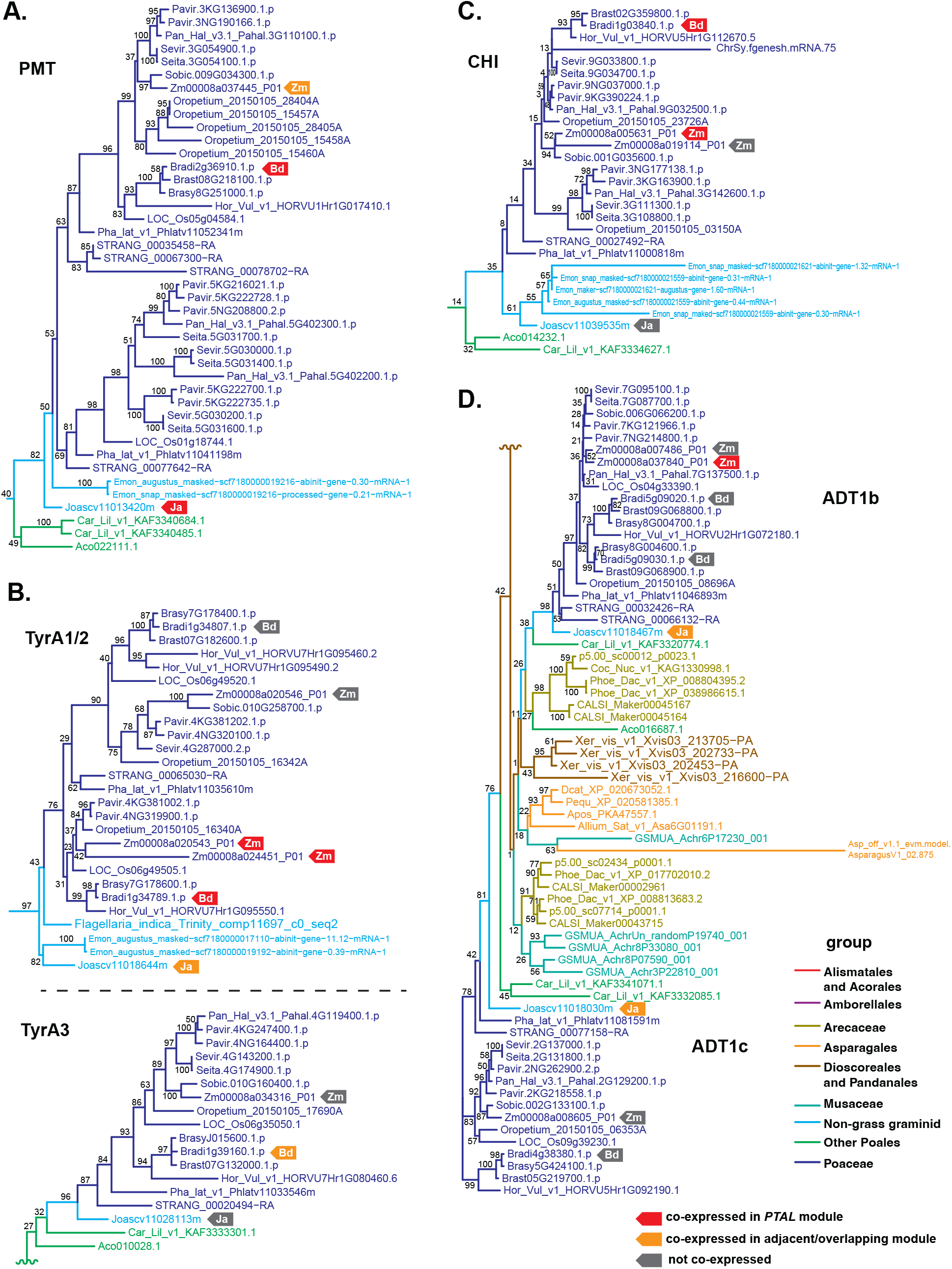
Some isoforms of lignin pathway genes diverged differently in expression patterns. All phylogenetic trees include monocot species from the species tree in **Fig. 2**. Groups are color coded according to their family or order(s). Red and yellow circles indicate the gene is co-expressed in a PTAL module and an adjacent or overlapping lignin gene module, respectively, whereas gray and open circles indicate the gene is not co-expressed and found in the dataset, respectively. PTAL: phenylalanine/tyrosine ammonia-lyase. **A.** A part of the phylogenetic tree of the PMT clade (**fig. S7**), an example of maintaining co-expression in the lignin pathway across species. PMT: *p*-Coumaroyl-CoA monolignol transferase. **B.** CHI phylogeny (full tree **fig. S21**), which shows a gain of co-expression in grasses whereas the genes are not co-expressed with lignin pathway genes in *J. ascendens* but are in *B. distachyon* and *Z. mays*. CHI: chalcone isomerase. **C.** TyrA phylogeny (full tree **fig. S16**), as an example of expansion of expression in grasses where TyrA1 is already co-expressed in *J. ascendens*, then TyrAnc is additionally recruited to the lignin network in *B. distachyon*. TyrA: arogenate dehydrogenase. **D.** ADT phylogeny (full tree **fig. S24**) as an example of co-expression loss from *J. ascendens* to grasses. Both ADT1b and 1c are co-expressed in *J. ascendens*, but none of the ADT isoforms are co-expressed in *B. distachyon* and only ADT1b is co-expressed in *Z. mays*. ADT: arogenate dehydratase.

We subsequently mapped the co-expression data of *B. distachyon, Z. mays,* and *J. ascendens* (**Figs. 3** and **S3**, **tables S3**, **S6**, and **S7**) onto these phylogenies. Four general patterns emerged. First, as anticipated, the same orthologous isoforms were co-expressed (within the *PTAL* module or an overlapping adjacent module) among *J. ascendens, B. distachyon*, and *Z. mays*. This pattern occurred for many core genes, such as *PMT* (**Fig. 5A**, **fig. S7**), *HCT* (**fig. S8**), *C3′H* (**fig. S9**), *CAD* (**fig. S10**), *F5H* (**fig. S11** – not co-expressed in *Z. mays*), *COMT* (**fig. S12**), *PAL/PTAL* (**fig. S13**), as well as *DHS*, though recently duplicated isoforms, *DHS1a* and *1b*, were present in the lignin co-expression network of *Z. mays* and *B. distachyon*, respectively (**fig. S14**). This pattern was also observed for the MYB activators SWAM1, MYB103, and MYB55/61 (**fig. S15** – MYB55/61 was not found in *B. distachyon*). *TyrA1/2*, *CCoAOMT1*, and *CCR1a/b* isoforms also showed conserved co-expression across *J. ascendens* and grasses, although other isoforms (*i.e.*, *TyrAnc*, *CCoAOMT2*, *CCR2b*) were co-expressed only in specific grasses (**Fig. 5B**, **figs. S16** to **S18**). Second, some gene families showed co-expression in grasses, but not in *J. ascendens*. This second case mostly included specialized lignin enzymes like the tricin pathway enzymes CHS, CHIL, CHI, A3′H- C5′H and FNSII (**Fig. 5C**, **figs. S19** to **S22**), or C3H/APX (**fig. S23**). Third, some genes were co-expressed with lignin pathway genes in *J. ascendens*, but not in grasses, which were seen for two of the four *ADT* isoforms (ADT1c and ADT2) (**Fig. 5D**, **fig. S24**), *FMT* (**fig. S7**), as well as *CSE* that were lost in many core grasses (**fig. S25**). Fourth and the most intriguing pattern was that one orthologous isoform was co-expressed in *J. ascendens*, while a different isoform was co-expressed in grasses. For example, the isoform *C4H2* was co-expressed with *PAL/PTAL* in *J. ascendens*, whereas the isoforms *C4H1a/1b* were in co-expressed modules adjacent to *PAL/PTAL* in grasses (**fig. S26**). For *4CL*, the isoforms in clades *1a* and *1b* were co-expressed with *PTAL* in grasses but were co-expressed in a different adjacent module in *J. ascendens*, while the *4CL1d* isoform was co-expressed with *PAL/PTAL* in *J. ascendens* but not in grasses; **fig. S27**). Similar patterns also occurred in *MYB58/63* and *MYB15/30*, which were co-expressed only in grasses and *J. ascendens*, respectively (**fig. S16**), consistent with the synteny results (**Fig. 4**). Overall, comparative analyses of co-expression and phylogenetic data suggest that extensive divergence and recruitment of different orthologous isoforms shaped the establishment of the lignin metabolic network during the evolution of Poales and grasses.

### *Joinvillea* possess FNSII but lack C5′H activity in the limiting step of tricin biosynthesis

Our co-expression network analysis show that tricin biosynthetic genes are in the lignin network of both *B. distachyon* and *Z. mays* but absent in that of *J. ascendens* (**Fig. 3**, **Fig. 5C**), suggesting that these genes were recruited into the lignin network after the divergence between grasses and *J. ascendens*. To investigate when the tricin pathway emerged during the grass evolution, we examined the functional origin of two key CYP450 enzymes, FNSII and A3′H-C5ʹH, involved in tricin biosynthesis (*56*). FNSII produces apigenin from naringenin, which is derived from *p*CA- CoA with the catalysis of CHS, CHI, and CHIL, while A3ʹH-C5ʹH together with COMT catalyze the sequential hydroxylation and methoxylation steps, respectively, to synthesize tricin (**Fig. 6A**). FNSII and A3ʹH-C5ʹH belong to the CYP93G and CYP75B subfamily, respectively (*83*, *95*). Grasses have an additional pair of CYP93G and CYP75B enzymes, F2H and F3ʹH, respectively, which are involved in the production of flavone *C*-glucosides (*e.g.*, isovitexin, orientin, **Fig. 6A**) (*59*, *96*).

**Fig. 6.**
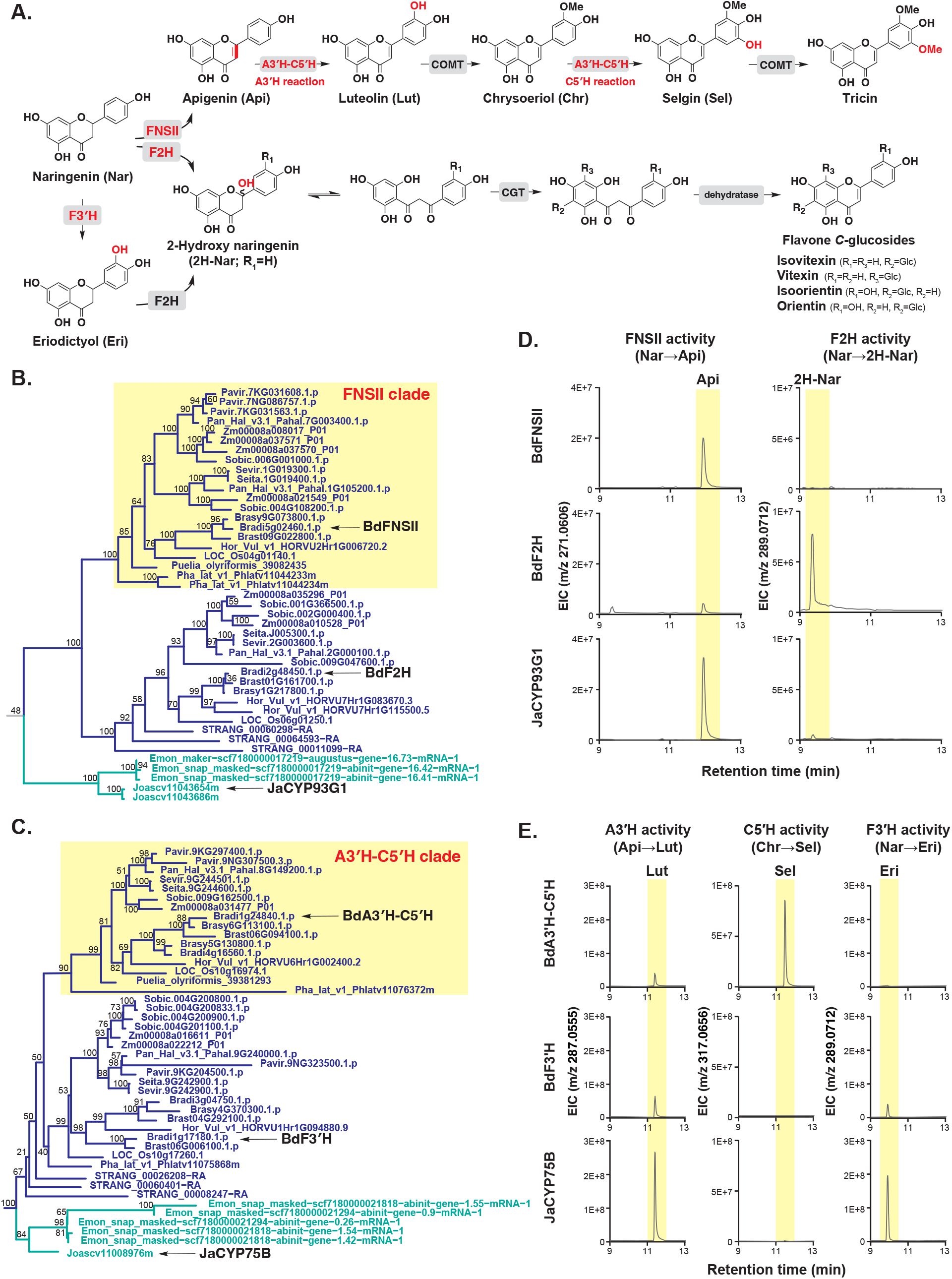
The *J. ascendens* CYP93G possess FNSII activity but the sole *J. ascendens* CYP75B lack C5′H activity. **A.** The proposed biosynthetic pathway and intermediates for tricin and flavone *C*-glucosides in grasses are presented, with the enzyme reaction steps whose activities were determined in this study highlighted in red. **B.** Phylogenetic tree of CYP93G enzymes that contains characterized FNSII and F2H from grasses. FNSII/F2H: flavone synthase II/ flavanone 2-hydroxylase. **C.** Enzyme assay of BdFNSII, BdF2H, and JaCYP93G1 for the conversion of naringenin into apigenin (FNSII reaction) and 2-hydroxy naringenin (F2H reaction). Extracted ion chromatograms (EIC) for the expected product mass are shown. Abbreviations for each compound are provided in **Fig. 6A**. FNSII/F2H: flavone synthase II/ flavanone 2-hydroxylase. **D.** Phylogenetic tree of CYP75B enzymes that contains previously characterized A3ʹH-C5ʹH and F3ʹH enzymes from grasses. A3′H-C5′H: apigenin 3′-hydroxylase/chrysoeriol 5ʹ-hydroxylase; F3ʹH: flavonoid 3′-hydroxylase. **E.** Enzyme assay of BdF3ʹH, BdA3ʹH-C5ʹH, and JaCYP75B for the conversion of apigenin into luteolin (A3′H reaction), chrysoeriol into selgin (C5′H reaction) and naringenin into eriodictyol (F3′H reaction). EIC for the expected product mass are shown. Abbreviations for each compound are provided in **Fig. 6A**. A3′H-C5ʹH: apigenin 3′-hydroxylase/chrysoeriol 5′-hydroxylase; F3′H: flavonoid 3′-hydroxylase.

Our phylogenetic analysis (**Fig. 6B**, **6C**, **fig. S22**) showed that grasses have two distinct clades for both CYP93G and CYP75B subfamily enzymes, one of which contain previously identified rice FNSII and A3′H-C5′H, respectively (*83*, *95*). However, CYP93G and CYP75B enzymes of *J. ascendens* were found in the outgroup clade, indicating that CYP93G and CYP75B duplicated within grasses after the divergence from *J. ascendens* and other non-grass graminids (**Fig. 6B**, **6C**, **fig. S22**). We hypothesized that the duplicated CYP93G and CYP75B functionalized after the duplication in grasses and were recruited into tricin and flavone *C*-glycoside biosynthesis.

To test this hypothesis, we cloned and investigated the activity of a CYP93G enzyme of *J. ascendens* (JaCYP93G1) together with two CYP93G paralogs of *B. distachyon* (BdFNSII and BdF2H, respectively, **Fig. 6B**). Cloning of the other CYP93G gene, *JaCYP93G2*, was not successful, likely due to its low expression. JaCYP93G1, BdFNSII, and BdF2H were expressed in yeast *Saccharomyces cerevisiae* and their potential FNSII and F2H activity were analyzed using the microsomal fractions and naringenin as a substrate. As expected, BdFNSII and BdF2H showed strong FNSII and F2H activities, respectively, while exhibiting very weak or undetectable reciprocal activities (**Fig. 6D**). Thus, FNSII and F2H enzymes have strict reaction specificity to produce apigenin and 2-hydroxynaringenin, respectively. Somewhat unexpectedly, JaCYP93G1 showed strong FNSII activity but only trace levels of F2H activity (**Fig. 6D**, **fig. S28**). This result suggests that FNSII activity, required for the committed step of tricin biosynthesis, was already present in the common ancestor of *J. ascendens* and grasses.

To investigate whether *J. ascendens* possesses a functional A3′H-C5′H enzyme capable of catalyzing the following two hydroxylation steps, converting apigenin into luteolin and chrysoeriol into selgin (**Fig. 6A**), we next cloned and characterized the sole CYP75B enzyme of *J. ascendens*, JaCYP75B, together with two CYP75B paralogs of *B. distachyon* (BdF3ʹH and BdA3′H-C5ʹH, respectively, **Fig. 6C**). Again, the yeast microsomal fractions expressing individual enzymes were analyzed for A3ʹH and C5ʹH activity, as well as F3ʹH activity involved in flavone *C*-glycoside biosynthesis (**Fig. 6A**). As anticipated, BdA3′H-C5′H showed A3ʹH and C5ʹH activities, whereas BdF3ʹH exhibited F3′H and A3′H activities (**Fig. 6E**), supporting a previous report in rice orthologs (*83*). Similarly to BdF3ʹH, JaCYP75B exhibited A3ʹH and F3ʹH activities but had no detectable C5ʹH activity (**Fig. 6E**, **fig. S29)**. C5′H activity therefore emerged after the gene duplication of CYP75B that took place within grasses. The finding suggests that the acquisition of C5ʹH activity in the CYP75B family was the pivotal step toward the evolution of the tricin biosynthetic pathway in grasses.

### Tricin biosynthesis emerged within the grass family

Inter-species co-expression and biochemical analyses suggest the emergence of the completed tricin biosynthetic pathway in core-grasses. To test if the biochemical evolution collaborates with the chemical evolution, we first conducted soluble metabolite analysis from core-grasses and their sister lineages. Core-grasses (*Sorghum bicolor*, *Setaria viridis*, and *B. distachyon*) were grown in an environmentally controlled growth chamber for 2-3 weeks before metabolite extraction. Fresh and green leaves of *J. ascendens* (8-months-old)*, S. angustifolia*, *Pharus lappulaceus*, and core-grasses *Sasa tsuboiana*, *Pleioblastus pygmaeus*, and *Phyllostachys aurea* were obtained from the University of Wisconsin Botany Greenhouse (**Fig. 7A**). Although we were unable to obtain the living plant of any species in the Puelioideae family, an herbarium specimen of *Puelia ciliata* was kindly provided by Missouri Botanical Garden (**Fig. 7A**). The obtained leaf tissues were freeze-dried, ground to powder, which were used for the metabolite extraction with 80% methanol, followed by LC-MS analysis (see **Methods**). Tricin intermediate chrysoeriol was detected from all species tested, including in *J. ascendens* (**Fig. 7A**), which aligns with the detection of FNSII activity in *J. ascendens* CYP93G1. Flavone *C*-glycosides, vitexin, isovitexin, orientin, and isoorientin (**Fig. 6A**), were also analyzed and detected in all species tested, although their accumulation was much lower in *J. ascendens* and *P. lappulaceus* compared to other species (**Fig. 7A**).

**Fig. 7.**
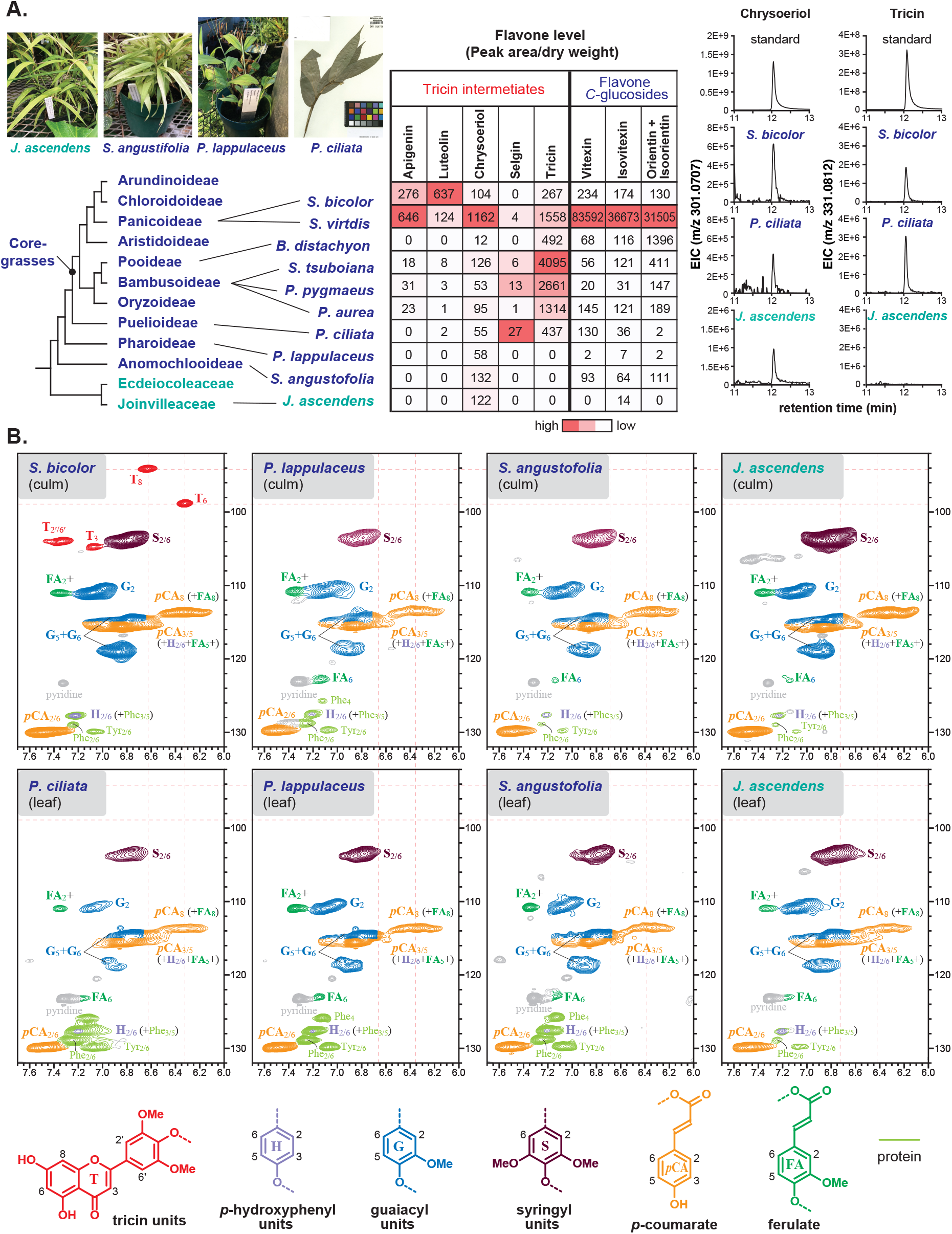
Tricin accumulation happened at the divergence of *Puelia* and core-grasses followed by with tricin-lignin formation event in core-grasses. **A.** Accumulation of tricin intermediates and flavone *C*-glucosides determined by LC-MS analysis of the acid hydrolysates. Heat maps coloring was applied independently to each flavone based on its normalized LC-MS peak intensity, whereas extracted ion chromatograms (EICs) show the expected product mass of chrysoeriol and tricin in positive-ion mode. See **fig. S30** for the LC-MS analysis of the MeOH extracts. **B.** Distribution of tricin-lignin determined by 2D HSQC NMR analysis of purified lignin fraction, demonstrating the unique tricin-lignin formation event in a core-grass (*Sorghum bicolor*).

In contrast to the tricin precursor chrysoeriol, which was ubiquitous among the plants analyzed, soluble tricin was detected only in the non-core grass *P. ciliata* and the core-grasses, but not in *P. lappulaceus*, *S. angustifolia*, or *J. ascendens* (**fig. S30**). Although some flavones may be glycosylated, soluble tricin was still not detectable from *P. lappulaceus*, *S. angustifolia*, or *J. ascendens* even after the methanolic extracts were acid-hydrolyzed to convert flavone *O*- glycosides into their free forms (**Fig. 7A**). These chemical data combined with the biochemical data (**Figs. 6** and **7**) support our contention that, although flavone *C*-glycosides and the earlier steps of tricin biosynthesis were already present in non-grass species, the complete pathway for tricin biosynthesis likely emerged in the common ancestor of *P. ciliata* and core-grasses. Although the full length FNSII and A3′H-C5′H homologs could not be obtained in *P. ciliata* for functional analyses, their partial gene fragments obtained from Shotgun genome sequencing (*97*) nested within the FNSII clade (*Puelia_olyriformis39082435* in **Fig. 6B**) and the A3ʹH-C5ʹH clade (*Puelia_olyriformis39381293* in **Fig. 6C**), further collaborating with our chemical analysis that *P. ciliata* possesses an active tricin biosynthetic pathway.

To further investigate the distribution of the lignin-bound tricin in comparison to soluble tricin, the lignin structures of a core grass *S. bicolor* and four core-grass sister lineages, *P. ciliata*, *P. lappulaceus*, *S. angustifolia*, and *J. ascendens,* were analyzed using the two-dimensional heteronuclear single-quantum coherence (HSQC) NMR method. The ground powders from these dried tissues were extracted with water, methanol, and chloroform to remove the extractives. The obtained cell wall residues were treated with cellulase to obtain lignin-enriched cell wall residues. The HSQC spectra of the lignin-enriched cell walls from all four sister species and *S. bicolor* displayed signals of canonical G, S, and H lignin units (**Fig. 7B**). As previously reported (*46*, *49*), all species that belong to commelinids exhibited clear and strong signals for *p*CA and FA residues (**Fig. 7B**). The signals for tricin residues were detected in the spectrum of *S. bicolor,* but not in those from all four sister species, *P. ciliata*, *P. lappulaceus*, *S. angustifolia,* and *J. ascendens* (**Fig. 7B**). Although *P. ciliata* accumulated soluble tricin (**Fig. 7A**), this non-core grass did not have a cell wall-bound tricin (**Fig. 7B**). These chemical analyses showed that the distribution of tricin- lignin in the cell wall is restricted to core-grasses.

## Discussion

Previous studies investigated the evolutionary history of lignin biosynthetic enzymes during land plant evolution (*98*, *99*). These studies focused on core genes in the lignin pathway, such as *PAL*, *4CL*, and *COMT*, that were also co-expressed in grass and Poales species we tested (**Fig. 2**). However, further evolution and diversification of this core and essential metabolic network in specific lineages remained unclear. Based on our integrated phylogenomic, biochemical, and chemical analyses, combined with prior knowledge of lignin biosynthesis, we can now trace the complex evolutionary history that reshaped the lignin metabolic network of grasses (**fig. S31**). The common ancestor of monocots already possessed deregulated TyrAnc and elevated tyrosine precursor availability (*42*). Within monocots, *p*-coumaroylation of G- and S-lignin subunits appeared in commelinids, followed by the emergence of the tyrosine-derived lignin biosynthesis just before the evolution of grasses and prior to ρWGD (*40*). Within the grass family, the flavone tricin was synthesized and subsequently incorporated into the cell wall lignin, as confirmed by biochemical and chemical analyses (**Fig. 6**). CSE was lost within many core-grasses (**fig. S25**) that likely acquired the C3H/APX shortcut pathway (**fig. S23**). Most recently, the first enzyme of the shikimate pathway, DHS, became deregulated in BOP (Bamboos, Oryzoideae, and Pooideae) grass species, directing carbon flux toward the dual lignin pathway (**fig. S14**) (*42*). The recruitment of different MYB transcription factors (**Fig. 4**, **fig. S15**) further refined the transcriptional regulation of lignin metabolic network during Poales and grass evolution. Extant grasses consequently possess unique lignin compositions and deposit substantial amounts of lignin into their scattered vascular bundles, likely providing critical mechanisms for some of the fastest-growing plant species (*e.g.*, bamboo, elephant grass, miscanthus). Although additional functional studies are needed to further validate the model (**fig. S31**), different network modules emerged at various stages of monocot evolution, remodeling and refining the lignin metabolic network through diverse mechanisms including tandem-gene and whole-genome duplications, gene losses, enzyme functionalization, and altered gene expression.

### Lignin *p*-coumaroylation and feruloylation originated in early monocots

*p*CA/FA-lignin is found across commelinid monocot orders, including Zingiberales, Arecales, and Poales (*46*, *49*). This coincides with the expansion of the *PMT/FMT* gene family (**Fig. 2**). The phylogeny suggests that an ancestral gene was duplicated within Poales (**fig. S7**) and sub- functionalized into PMT and FMT, which preferentially utilize *p*CA-CoA and FA-CoA substrates, respectively, in grasses (*100*). We speculate that the ancestral PMT/FMTs of commelinids were promiscuous enzymes capable of catalyzing both *p*-coumaroylation and feruloylation. *PMT* is incorporated into the lignin co-expression network in both *J. ascendens* and grasses, whereas *FMT* was co-expressed only in *J. ascendens* but not in grasses (**Fig. 3**), consistent with the lower feruloylation than *p*-coumaroylation observed in grass lignin (*46*). Together, *p*CA/FA-lignin and PMT/FMT function evolved early in monocot evolution, prior to the divergence of grasses and other Poales species. As *p*CA/FA-CoA substrates are broadly available in vascular plants, *p*CA/FA-lignin likely evolved relatively easily, and repeatedly (*28*, *46*, *47*), without necessarily requiring major changes in the upstream precursor pathways.

### Grasses shifted from phenylalanine to tyrosine-derived lignin pathways after the PTAL evolution

The PTAL enzyme responsible for the dual entry pathway from phenylalanine and tyrosine evolved through tandem gene duplication followed by neofunctionalization just before the emergence of grasses (**Fig. 4**, **fig. S13**) (*40*). The introduction of the second entry pathway via tyrosine was likely facilitated by the existing “high-tyrosine background” due to the deregulated TyrAnc, which is found across all Poales, including those lacking PTAL (**Fig. 5**; **fig. S16**) (*42*). This study additionally found that *PTAL* and *PAL* are both co-expressed with core lignin genes in grasses as well as in *J. ascendens*, suggesting that the newly evolved PTAL became integrated into the lignin metabolic network in the common ancestor of *J. ascendens* and grasses. Within grasses, *PALs*, some of which are co-expressed with *PTAL* (**fig. S13**), further duplicated multiple times hinting at their coordinated function.

The *J. ascend*ens lignin network also contained *C4H* and three of four *ADTs* (**figs. S26**, **S24**) whereas canonical *TyrA*, which is not duplicated into *TyrA1* and *2*, was only loosely co-expressed (**Fig. 5B**, **fig. S16**). Despite the presence of functional PTAL enzymes (*40*), *J. ascendens* therefore appears to still rely more heavily on phenylalanine-derived lignin biosynthesis than grasses. In contrast, the grass lignin co-expression network contained *TyrAnc* and *TyrA1* (**Fig. 5B**, **fig. S16**) but had only one or no *ADT* (**Fig. 5D**, **fig. S24**) and loose association with *C4H* (**fig. S26**). This finding suggests that the acquisition of PTAL likely reduced the reliance on ADT and C4H that work together with PAL (**Fig. 1**). Thus, subsequently to the PTAL evolution in graminids, grasses likely shifted from phenylalanine to energetically more efficient tyrosine-derived entry pathways (*39*) for lignin biosynthesis.

### Tricin-lignin evolved within grasses through the emergence of C5′H activity

Tricin-lignin is reported to be a unique characteristic of grasses and some specific monocot species (*55*, *101*). However, its evolutionary origin and the molecular mechanism underlying its emergence remained unknown. By integrating network analysis with biochemical and chemical analyses, this study revealed stepwise evolution of tricin-lignin formation in the grass lineage (**Fig. 7**; **fig. S31**). The grass CYP75B subfamily duplicated likely because of the ρWGD (**Fig. 6**; **fig. S22**), resulting in two separate clades. C5ʹH activity emerged from one clade at the divergence of *Puelia* and core-grasses and was the key innovation that led to the acquisition of the tricin

biosynthetic pathway (**Fig. 6**). Subsequently, within core-grasses, soluble tricin became incorporated into lignin polymer (**Fig. 7**). The precise mechanism for this incorporation remains unknown; it may involve coordinated expression of the tricin pathway and lignin biosynthetic genes or, alternatively, require additional component(s) such as transporters or polymerization enzymes. Future comprehensive genome and transcriptome analysis of *Puelia* species could shed light on the molecular mechanisms underlying the tricin incorporation into lignin. Our findings demonstrate that tricin biosynthesis and its subsequent incorporation into lignin are features acquired within core-grasses; however, soluble or lignin-bound tricin have been detected in some monocots outside of grasses, such as vanilla (Orchidaceae), palms (Arecaceae), and papyrus (Cyperaceae) (*55*, *101–103*). The tricin biosynthetic pathway and its currently unknown incorporation mechanism into the lignin polymer therefore likely evolved independently multiple times across these monocot and other plant lineages.

### Gene losses in grasses reshaping G and S lignin monomer synthesis

The hydroxylation of *p*CA at its 3-position is required for G/S lignin synthesis (**Fig. 1A**). *p*CA- CoA, generated by 4CL, is initially converted by HCT into *p*-coumaroyl shikimate (*104*). C3ʹH catalyzes the subsequent 3-hydroxylation producing caffeoyl shikimate, which is then converted to caffeic acid or caffeoyl-CoA by CSE or the reverse reaction of HCT, respectively (*74*, *105*) (**Fig. 1A**). Our data revealed that multiple genes were recruited or lost to further refine this key metabolic connection within the grass lignin metabolic network. For example, *CSE* was present and co-expressed within the lignin network in *J. ascendens*, but *CSE* was absent in the genomes of many grasses, including *B. distachyon* and *Z. mays* (**Fig. 2**, **fig. S25**) (*27*, *75*). Conversely, *HCT* is duplicated into two copies in all grasses (**fig. S8**), tightly co-expressed with other lignin pathway genes in both *B. distachyon* and *Z. mays* (**Fig. 3**), and present in one of the *PAL/PTAL* synteny in *B. distachyon* (**Fig. 4**), whereas *J. ascendens* HCT exhibits loose association with the lignin co-expression network (**Fig. 3**). A previous study demonstrated that *B. distachyon* possesses a C3H/APX enzyme that directly produces caffeic acid from *p*-coumaric acid (*43*), bypassing four steps catalyzed by 4CL, HCT, C3ʹH, and CSE (**Fig. 1A**). *C3H/APX* orthologs are present in all grasses and non-grass graminids (**fig. S23**) but were co-expressed with lignin pathway genes only in *B. distachyon* but not in *Z. mays* and *J. ascendens* (**Fig. 3**). This contrasts with *C3′H* orthologs, which were found in the lignin co-expression network in all three species (**Fig. 3**). This was also the case for *4CL* orthologs, although the specific *4CL1* isoforms were co-expressed with lignin pathway genes in grasses, whereas three isoforms from both *4CL1* and *2* were identified in the *J. ascendens* lignin network (**fig. S2A**). These findings indicate that the first three steps catalyzed by 4CL, HCT, and C3′H are conserved for G and S lignin synthesis (**Fig. 1A**), but the functional divergence or loss of additional components, such as C3H/APX and CSE, further reshaped this critical gateway for G/S lignin biosynthesis in various grasses.

### Various MYB activators fine-tuned the lignin transcriptional network during grass evolution

Multi-layered transcriptional regulatory mechanisms fine-tune lignin biosynthesis and deposition (*80*, *81*, *106*). Considering the substantial modifications observed within the lignin metabolic network during grass evolution, MYB evolution may be associated with changes in transcriptional regulation (**fig. S3**). Although our combined phylogenetic and co-expression analyses revealed that well-known MYB transcriptional factors (*e.g.*, *MYB46/83, MYB42/85, MYB55/61, MYB103, SWAM1*) activating the lignin biosynthesis are well co-expressed with lignin biosynthetic genes in both grasses and *J. ascendens*, *MYB58/63* showed co-expression only in grasses, but not in *J. ascendens* (**fig. S15**). Instead, an *MYB15/30* ortholog, which is reported to induce stress-induced lignification (*88*, *89*, *107*), was found to be a part of the lignin biosynthetic network in *J. ascendens*. The *PAL/PTAL* syntenic block also contained *MYB58/63* in grasses and *MYB15/30* in *J. ascendens* (**Fig. 4**). These results suggest that different MYB transcription factors, linked to distinct upstream signals, perhaps stress-induced in *J.ascendens* while constitutive expression in grasses, were recruited to regulate and fine-tune the expression of various components of the grass lignin metabolic network.

### Conclusion: Evolution of the lignin metabolic network

In the context of the grass lignin metabolic network, certain steps and modules evolved at distinct times during monocot evolution, providing an illustrative example of the evolutionary dynamics of establishment and remodeling of such complex yet coherent metabolic networks. Prior modeling studies suggested that enzymes do not experience uniform constraints in their evolution; those intricately connected to many parts of the network or other networks, or those exhibiting high flux, are generally more constrained than their less-connected or lower-flux counterparts (*108*, *109*). This pattern was also observed in our present study. Tricin biosynthetic enzymes appear to have evolved multiple times through convergent evolution, as the tricin pathway is at a terminus of the lignin metabolic network and is not constrained by downstream pathways. This is also true for the PMT/FMT enzymes and the widespread *p*CA/FA-lignins across multiple lineages of monocots and eudicots (*46*, *47*, *49*). In contrast, PAL/PTAL enzymes carry high flux and are highly connected to other enzymes in multiple pathways that depend on the products of PAL/PTAL- catalyzed reactions (*e.g.*, phenylpropanoid and benzenoid biosynthesis) or that provide phenylalanine and tyrosine precursors (*e.g.*, TyrA, ADT, and enzymes of the shikimate pathway). This complexity likely contributes to the rare occurrence of *PTAL*, which evolved only once within the plant kingdom in the graminid Poale ancestor (*40*), and was likely pre-adapted (*18*) to have a favorable metabolic background (*e.g.*, high tyrosine availability). Historical contingency can also influence evolvability of certain metabolic alterations (*20*), such as the loss of *CSE* that coincides with the gain of *APX/C3H* or potential *HCT* alteration, or other changes that might be associated with the dual PTAL entry pathways. Gaining knowledge on these potential metabolic constraints and contingencies is crucial for engineering and redesigning the metabolism of various plants having different evolutionary history and resulting network structures. The generated dataset and workflow can be further utilized to discover unknown genetic components associated with the evolution of the grass lignin metabolic network and other complex traits uniquely evolved in this critical plant family.

## Methods

### Gathering expression data and performing RNA sequencing

We constructed 3 co-expression networks, in *B. distachyon*, *Z. mays*, and *J. ascendens*. For *B. distachyon* we used transcriptome data from (*76*). This is a developmental microarray data set with the following tissues/stages where DAG is days after germination, DAH is days after heading, DAF is days after fertilization: roots (10-35 DAG), etiolated shoots (3 DAG), de-etiolated shoots (3 DAG), first node (10, 17, 27, 60 DAG), first node and adventitious roots (35 DAG), last node (35 DAG), lower part and upper part of inclined node (42 DAG), first internode (10 DAG, 17, 27, 35, 60), second internode (17, 27 DAG), last internode (35, 60 DAG), leaf (10, 17, 27, 60 DAG), young leaf <6 cm (60 DAG), mature leaf fully expanded (60 DAG), Peduncle (42 DAG), Spikelet pedicel (42 DAG), Young spikelet (3 DAH), First spikelet internode (42 DAG), Last spikelet internode (42 DAG), Endosperm (11 DAF), Whole grain (11 DAF, 31 DAF, 2 years), Coleoptile (10 DAG, 17+27 DAG).

For *Z. mays* we used data from (*85*). This is a development RNA-seq data set with tissues of different developmental stages where DAS is days after sowing, DAP is days after pollination, VT is vegetative tasseling, V*n* is vegetative stage corresponding to the number of emerged leaves, and R1 and R2 refer to reproductive 1 and 2. Tissues include roots (3 DAS, 7 DAS, V7, V13), leaves (6 DAS, V1, V3, V5, V7, V9, VT, 0-30 DAP, R2) stem (V1, V3), internode (V5, V9, 0-30 DAP), reproductive tissues (tassels (V13, V18), anthers (R1), silk (R1), cob (V18, R1), and seed (whole seed- 2-24 DAP, endosperm- 12-24 DAP, embryo- 16-24 DAP, pericarp- 18 DAP).

Finally, we performed RNA-sequencing on *J. ascendens*. We isolated RNA from the following tissues: 9-month-old internode, 9-month-old leaf, 9-month-old root (3 replicates each), young shoot and root 2 months after germination (3 replicates each), flowers (2 replicates) and peduncle (1 replicate) (**Fig. 1C**). We were only able to obtain one flowering stem from the *J. ascendens* plant, which is why there are fewer replicates for the flower and peduncle tissues. Tissue was ground in liquid nitrogen, and total RNA was isolated from ground tissue using the Plant RNeasy Plant Kit (QIAGEN). For RNA extraction from the root, total RNA was extracted using the CTAB/LiCl method (*110*) with modifications (*111*) to improve the yield and quality. Paired-end RNA-sequencing was performed at Azenta Life Sciences (https://www.azenta.com/) using the Illumina platform, 2×150bp configuration, with a single index per lane. Quality scores and read count are reported in **table S12**. RNA sequencing reads were processed according to https://github.com/HiroshiLab/RNA-seq_data_processing. Briefly, reads were trimmed via Trimmomatic 0.39 (*112*), with a maximum mismatch of 2, palindrome clip threshold of 30, a simple clip threshold of 10, a sliding window size of 4 and required quality of 20, and a minimum length of 36. FastQC (*113*) (https://www.bioinformatics.babraham.ac.uk/projects/fastqc/) was then performed to ensure sequence quality >20. Reads were mapped using HiSAT2 version 2.2.1 (*114*) with no discordant reads mapped to the *Joinvillea ascendens* genome v1.1, genome ID 587 on Phytozome (https://phytozome-next.jgi.doe.gov/info/Jascendens_v1_1). Reads were sorted using SAMtools version 1.15.1 (*115*), and only unique reads counted via the python package HTSeq 0.11.1 (*116*). Finally median TPM (Transcript Per Kilobase Million) was calculated for each same using R version 4.2.1 (2022-06-23) and the scatter (version 1.0.4) package using calculateTPM.

### Construction of co-expression networks

Co-expression networks were calculated via Spearman’s mutual rank (MR-SP) as in (*5*), using the pipeline https://github.itap.purdue.edu/jwisecav/mr2mods. *PTAL* and *TyrA1* were first used as bait genes to find modules in the grass species *B. distachyon* and *Z. mays*. Different decay factors which determine the overall size of the modules were tested, and for *B. distachyon*, the decay factor = 25 was chosen to get module sizes that include the *PTAL* gene to range from ∼100 genes to ∼200 genes. This size module seemed to include many other lignin pathway genes but was not too large that it included distantly related genes. For *Z. mays*, a larger decay factor = 35 was chosen to get *PTAL/TyrA1* modules ranging from ∼50 to ∼300 genes in size. For *J. ascendens*, because it was unknown whether *PTAL* was involved in lignin biosynthesis in this species, both *PTAL* and *PAL* genes were used as bait genes to find modules. The decay factor = 45 was chosen to select module sizes for *PAL/PTAL* between ∼50 and ∼250 genes. Modules were created using the both the ClusterOne program (*117*), as well as with Cytoscape (https://cytoscape.org/) (*118*), the latter by loading MR-SP network into Cytoscape, the finding the gene of interest (*i.e.*, *PTAL*), and choosing all direct connections to the gene.

For lignin pathway genes that did not co-express in *PTAL/TyrA1* clusters for *B. distachyon* and *Z. mays*, or in *PTAL/PAL* modules for *J. ascendens*, we found the co-expression modules in which they occurred. For *B. distachyon* these genes included: *TyrA2* (bradi1g34807), *TyrAnc* (bradi1g39160), *A3′H-C5′H* (bradi4g16560), *CCoAOMT* (bradi4g33340, bradi3g39380, bradi3g39400, bradi3g39410), *C3H/APX* (bradi1g16510, bradi1g65820), and *C4H* (bradi2g31510, bradi2g53470, bradi3g43160) (**table S3**). We also found individual modules for *PMT* (bradi2g36910), even though this is co-expressed with *PTAL* in *B. distachyon*. *Z. mays* genes not found in the *PTAL* modules included: *PMT* (Zm00008a037445), *TyrAnc* (Zm00008a034316), and *C3H/APX* (Zm00008a001157, Zm00008a035553, Zm00008a010007), and individual modules were found for these genes. *TyrA2* was not found in the *Z. mays* expression data set. To be consistent with *B. distachyon*, we also found individual modules for *A3′H-C5′H* (Zm00008a031477), even though this is co-expressed with *PTAL* in *Z. mays* (**table S6**). For *J. ascendens*, genes not present in *PAL* or *PTAL* modules included *CCR* (Joascv11029948m.g, Joascv11052244m.g, Joascv11029426m.g, Joascv11060192m.g, Joascv11060195m.g), *C3′H* (Joascv11045673m.g), *CAD* (Joascv11029231m.g, Joascv11020448m.g, Joascv11008280m.g, Joascv11032347m.g, Joascv11059723m.g), *C3H/APX* (Joascv11004324m.g), *CYP75B* (Joascv11008976m.g), *CHI* (Joascv11039535m.g), *F5H* (Joascv11058161m.g), *FMT* (Joascv11024051m.g), *CYP93G* (Joascv11043654m.g, Joascv11043686m.g), *HCT* (Joascv11054939m.g), *TyrA1* (Joascv11018644m.g) and *TyrAnc* (Joascv11028113m.g). Additionally, we investigated modules containing upstream shikimate pathway genes *ADT* (Joascv11004322m.g, Joascv11018467m.g, Joascv11018030m.g, Joascv11058337m.g) and *DHS* (Joascv11000409m.g), as well MYB transcription factors *SWAM1* (Joascv11035514m.g), *MYB15/30* (Joascv11021278m.g) and *MYB103* (Joascv11014856m.g) (**table S7**).

Next, we determined module overlap via custom python scripts available on GitHub: https://github.com/HiroshiLab/GO-term-enrichment. For each pair of modules, A and B, percent overlap was calculated as the percent of overlapping genes from A and B divided by the sum of the total number of genes from both modules (union). We calculated percent overlap for all modules for each species and then used this distribution to calculate percentiles (95^th^, 99^th^, and 99.5^th^) of the population. The percent overlap cutoff for the 99.5 percentile was used to determine significance, where any overlap percentage over the 99.5 percentile was deemed significant at α=0.005. In *B. distachyon*, significant overlap was above 0%, in *Z. mays*, significant overlap was above 1.8%, and for *J. ascendens* significant overlap was above 1.8% (**table S5**). For all lignin modules we found whether they were significantly overlapping, first with *PTAL* and then with each other. Because *PTAL* and *TyrA1* genes *in B. distachyon* and *Z. mays* were either already in the same cluster (100% overlap), or in clusters that significantly overlapped compared to the 99.5^th^ percentile at high percentages (20-40% in *B. distachyon*, 1-12% in *Z. mays* except for 2 *TyrA1* modules in *Z. mays*) (**table S5**), we combined separate *PTAL/TyrA1* clusters together to make one consensus cluster. In *J. ascendens*, *PAL* and *PTAL* modules overlapped 9-15% except for one pair, and we thus combined *PAL* and *PTAL* modules to create one consensus module.

### GO Pathway enrichment analysis of gene networks

Gene ontology (GO) terms were downloaded from http://geneontology.org/, version 1.2. Gene associations were downloaded from Phytozome version 13, https://phytozome-next.jgi.doe.gov/. For each species, the following versions for gene annotation were used: *B. distachyon*, version 3.2; *Z. mays*, version PH207 version 1.1; *J. ascendens* PH207 version 1.1. Pathway annotations were downloaded from Plant Metabolic Network (https://plantcyc.org/), version 15. Pathways were annotated to *B. distachyon* version 3.2 and *Z. mays* AGP version 4. There is no pathway annotation for *J. ascendens*. Because both *B. distachyon* and *Z. mays* had different annotation versions for the expression data set, we used BLAST (NCBI Basic Local Alignment Search Tool) version 2.10.0 (https://blast.ncbi.nlm.nih.gov/) between the different versions to find the reciprocal best match. This was used to match the expression data gene version to the GO and pathway annotation version. To find GO and pathway enrichment, we used the Fisher’s exact test from python package fisher 0.1.10 on all modules and called significant enrichment for those modules enriched in a pathway or GO term with an FDR rate less than 0.05 (https://github.com/HiroshiLab/GO-term-enrichment).

### Finding Orthogroups and constructing phylogenies

First OrthoFinder (*119*) was used to get orthogroups for each enzyme family. Two OrthoFinder runs were used, the first with species selections from green plant lineages (see **table S8**), and then species from monocot lineages with *Amborella trichopoda* as an outgroup (see **table S9**). The following enzyme families were included and are annotated in the lignin pathway figure (**Fig. 1A**): DHS, TyrA, ADT, PAL/PTAL, C4H, 4CL, C3H/APX, HCT, C3′H, CSE, CCoAOMT, CCR, CAD, F5H, COMT, PMT/FMT, CHS, CHIL, CHI, A3ʹH-C5ʹH. Additionally, orthogroups from known lignin pathway regulators were included: the transcription factors MYB. To obtain the correct lignin pathway orthogroups, known genes from *B. distachyon* were used as bait genes. For *Z. mays*, the genome used for phylogeny construction (PH207 (*86*)), was a different genome than what was available for co-expression (B73 (*65*)). Therefore, in order to compare the genes in the phylogeny to the genes in the co-expression network, we found the reciprocal best BLAST match based on percent similarity where a gene pair with the highest similarity was selected as the match for Z. mays genes across genomes. For CHI analysis, the orthogroup containing a *B. distachyon* gene lacked a functionally characterized rice CHI (*57*). Therefore, two orthogroups, initially identified using *B. distachyon* and rice genes as bait, were combined. Before building gene trees, genes from an orthogroup were filtered to remove duplicate genes and genes that were truncated (either less than 50 amino acids or less than 2 times the standard deviation of the mean). Genes from a given orthogroup were then counted for each species to build the heatmaps of gene copy number (**Figs. 2** and **S1**) and used to build gene trees. MAFFT v7.453 (2019/Nov/8; https://mafft.cbrc.jp/alignment/software/) was used to align sequences. ModelTest (*120*) was then used to select the appropriate evolutionary model for the orthogroup. Finally RAxML-ng (*121*) was used to build the phylogenetic tree with 10 parsimonious trees, 10 random trees, and the orthogroup tree from OrthoFinder as starting trees. 200 bootstrapping trees were run for each orthogroup with the FBP bootstrap metric. For monocot trees, Amborella genes were used as an outgroup. For each tree, the orthogroup and evolutionary model can be found in **table S10**. Overall methods for building the phylogenies can be found here: https://github.com/HiroshiLab/Building-gene-trees.

### Gene expression analysis by quantitative reverse transcription PCR (qRT-PCR)

Total RNA extracted above was reverse transcribed into cDNA using SuperScript IV reverse transcriptase (Invitrogen). Quantitative PCR analysis was conducted on Stratagene Mx3000P (Agilent Technologies) using GoTaq qPCR Master Mix (A6001, Promega) and primers listed in **table S11**. Cq values were calculated using LinRegPCR. For relative quantification, a ubiquitin gene (*JaUBI10*; Joasc.02G086900) was used as an internal control. To determine transcript, copy numbers, standard curves were constructed using serially diluted plasmids (pML94, Invitrogen) harboring corresponding qRT-PCR amplicons of *JaPTAL* and *JaPAL*.

### Finding syntenic regions across species

Synteny analyses were performed using COGE (https://genomevolution.org/coge/), across selected Poales species with the following options: DAGchainer: relative gene order, maximum distance between 2 matches: 20 genes, minimum number of aligned pairs: 5 genes, merged syntenic blocks with quota align, window size 100 genes, use all genes in the target genome, calculate synonymous substitution rates, and tandem duplication distance is 10. Synteny was calculated between the following species: *B. distachyon* (version 556, 3.0) and *J. ascendens* (version 1.1); *B. distachyon* (version 556, 3.0) and *Z. mays* (version PH207 UMN 1.0); *B. distachyon* (version 556, 3.0) and *P. latifolius* (version 1.0); *B. distachyon* (version 556, 3.0) and *S. bicolor* (version 454, 3.0.1); *P. latifolius* (version 1.0) and *S. angustifolia* (version 1.1); *S. angustifolia* (version 1.1) and *J. ascendens* (version 1.1); *J. ascendens* (version 1.1) and *A. comosus* (version 3); *J. ascendens* (version 1.1) and *P. latifolius* (version 1.0). For each species pair, syntenic blocks containing the following genes were searched for: PAL/PTAL, CYP93G (FNSII), CYP75B (A3′H-C5′H), TyrA, and PMT.

### Cloning of CYP93Gs and CYP75Bs

Total RNA used for the cloning was extracted from *B. distachyon* and *J. ascendens* using Plant RNeasy Mini Kit (QIAGEN). The obtained RNA was treated with DNase (QIAGEN) and cDNA was synthesized with Super Script IV VILO master mix (Invitrogen). The coding sequence of *CYP93G*s and *CYP75B*s from *B. distachyon* and *J. ascendens* were amplified by nested PCR using the corresponding cDNA, gene specific primers, and PrimeSTAR MAX DNA polymerase (Takara). The protein expression vectors in *S. cerevisiae* were generated by using yeast toolkit (*122*). The obtained PCR products were purified and used as templates for the other PCR reaction with primers harboring the overhang sequences with the donor vector pYTK001. The obtained PCR products were cloned into *Bam*HI-linearized entry vector pYTK001 by In-Fusion cloning (Clontech). After recovering the entry vectors with coding sequences of interest, the sequences were confirmed by sanger sequencing and subsequently subjected for the golden gate assembly using *Bsa*I sites to generate the final yeast expression vector. All the primers used in this study are listed in **table S11**. The final vector structure is illustrated in **fig. S32**.

### Recombinant protein expression and preparation of the microsomal fraction

For the recombinant protein expression, the cloned constructs were transformed into transgenic *S. cerevisiae* harboring *Arabidopsis thaliana* CYP450 reductase expressed (WAT11, (*123*) by following the protocol written in Yeast Protocol Handbook (Takara). The obtained colonies were grown in a synthetic medium lacking uracil (SD-Ura) containing 1% glucose at 30 ℃ and 225 rpm. After the culture was saturated (OD >1.5), the cells were collected by centrifuge, resuspended with SD-Ura medium with 1% galactose, and diluted into OD600=0.4. The cells were cultivated at 30 ℃ to induce the proteins, and after 16 h incubation, the cells were harvested with centrifuge and stored at -80 ℃ until use. The pellets were thawed, resuspended with the buffer containing 50 mM Tris-HCl (pH 7.4), 2 mM EDTA, and 100 mM 2-mercaptoethanol, and centrifuged. The recovered pellets were washed with 50 mM Tris-HCl (pH 7.4), 100 mM KCl, and 100 mM 2- mercaptoethanol, and centrifuged. The pellets were resuspended with 50 mM Tris-HCl (pH 7.4) containing 2 mM EDTA and 0.6 M sorbitol. Acid washed glass beads (450-600 um diameter, Sigma-Aldrich) were added, and the cells were ground using a genogrinder (Spex Sample Prep) at 1000 rpm for 10 min with occasional cooling and the supernatants were recovered by centrifuge. This step was repeated three times. The combined supernatant was submitted for centrifuge at 10000 g and 4 ℃ for 10 min, and the supernatant was further centrifuged with Ultracentrifuge (L8-80MR, Beckman) at 105,000 g and 4 ℃ for 90 min. The pellets were suspended with 50 mM Tris-HCl (pH 7.4) containing 1 mM EDTA and 20% glycerol, and the suspensions were used for the enzyme reaction. The protein concentration was determined using the BioRad protein assay dye (BioRad). The purity was determined by Western blotting with OctA-HRP antibody (sc- 166355, Santa Cruz) followed by the calculation of the signal intensity with ImageJ software.

### Enzyme assay of CYP93G and CYP75B enzymes

CYP93G enzymes were investigated for FNSII and F2H activities, and CYP75B enzymes were examined for A3ʹH and C5ʹH activities. NADP^+^ (1 mM), glucose 6-phosphate (10 mM), and glucose 6-phosphate dehydrogenase (1 unit) were mixed in 50 mM potassium phosphate (pH 7.4) buffer in a total volume of 450 μl and pre-incubated at 30 ℃ for 5 min to permit the generation of NADPH. The microsomal preparations (50 μl) were added and further incubated for 3 min. Five μL of 5 mM substrate (naringenin for FNSII and F2H, apigenin for A3ʹH, and selgin for C5ʹH activity) were added to start the reaction. After 30 min of the incubation, the reactions were terminated by adding ethyl acetate. The products were recovered by partition, and the combined organic phase was concentrated by speedvac. The pellets were suspended by 80% MeOH and subjected for the LC-MS analysis [Vanquish (LC) - Q Exactive (MS), Thermo Fisher]. Analytical conditions were as follows. LC part: ACQUITY UPLC HSS T3 column (1.8 μm, 2.1 × 100 mm, Waters); solvent system, solvent A (water including 0.1% [v/v] formic acid) and solvent B (MeOH); gradient program 99% A/1% B at 0 min, 95% A/5% B at 4.5 min, 75% A/25% B at 5 min, 65% A/35% B at 9.5 min, 50% A/50% B at 10 min, 30% A/70% B at 17.5 min, 1% A/99% B at 18 min, 1% A/99% B at 22 min, 99% A/1% B at 22 min, and 99% A/1% B at 26 min; flow rate 0.4 mL/min. The spectra were recorded using full-scan-ddMS^2^ positive-ion mode with the following setting. Full-MS: scan range *m/z* 60-900; resolution 70,000; maximum scan time 200 ms. dd-MS^2^: resolution 17,500; maximum scan time 50 ms; Isolation window 1.5 m/s; normalization collision energy (NCE) 20, 40, 80. The reaction products apigenin, luteolin and eriodictyol were identified by comparing the retention time and mass spectrum of the authentic standards (**fig. S28** and **S29**). For selgin and 2-hydroxy-naringenin, we compared the MS/MS fragmentation pattern with that in the literature (*83*, *96*) and detected the major peaks reported (m/z 163, 153, and 121 for 2-hydroxy-naringenin and m/z 302, 274, 203, and 153 for selgin).

### Metabolic analysis of flavones in graminid tissues

*Brachypodium distachyon* (Bd21), *Setaria viridis*, and *Sorghum bicolor* (RTx623) were grown for two to three weeks at 25 ℃ under the 12 h day/12 h night condition and 60% of humidity. The leaves were harvested from them and freeze-dried. Leaves from *J. ascendens* (1 year after germination), *S. angustifolia*, *P. lappulaceus*, *P. aurea*, *P. pygmaeus*, and *S. tsuboiana*, which were grown in a greenhouse of UW-Madison, Department of Botany, were harvested and freeze-dried. To minimize protein contaminants, we used well-matured or dead leaves for *J. ascendens* (5-years-old)*, S. angustifolia*, and *P. lappulaceus*, and the same herbarium specimen for *P. ciliata*. *S. bicolor* obtained from the Great Lakes Bioenergy Research Center, were harvested with a John Deere 7350 Self-propelled forage harvester/chopper and dried. The tissues were ground using a genogrinder (Spex Sample Prep). The powder was mixed with 80% MeOH containing ring ^13^C-labeled tyrosine as an internal control, sonicated for 20 min, and centrifuged to recover the supernatant. The steps were repeated twice. The obtained supernatant was combined and concentrated with speedvac. The pellet was resuspended with 80% MeOH and submitted for LC-MS analysis using the same analytical condition with enzyme reaction product. For the acid hydrolysis, the dried pellets of the extracts were suspended with 2 M HCl and heated at 85 ℃ for 2 h. The solvent was dried down on a speedvac and the obtained pellets were resuspended with 80% MeOH and subjected for LC-MS analysis with the same method with the enzyme assay (*96*). The compounds were identified by comparing the retention times and mass fragmentation patterns with authentic standards except for selgin (**fig. S30**). Selgin was identified by comparing its fragmentation pattern with that of the C5ʹH reaction product.

### 2D HSQC NMR-based structural analysis of graminid lignin

Freeze-dried materials (Internode or leave) were homogeneously pulverized with a ball mill (Retch MM400, 50 mL hardened steel jar, 1 × 15 mm hardened steel grinding ball, 30 Hz for 60 s), extracted with distilled water and 80% ethanol aqueous solution, and then freeze-dried to give CWRs. CWRs were further ball-milled using planetary ball-milled (Fritsch Pulverisette 7 with (500-600 mg CWR being placed in 20 mL agate jars with 10 × 10 mm agate grinding balls, the CWRs were milled for 22 grinding cycles at 600 rpm for 10 min, with 10 min rest time between cycles, and reversing direction each cycle). Aliquots of the ball-milled CWRs were directly swelled into DMSO-*d_6_*/pyridine-*d_5_* [4:1 (v/v), 600 μL] in a 5-mm external-diameter NMR tube, under sonnication, for whole cell wall NMR analysis (*124*, *125*). The remaining ball-milled CWRs were further digested with cellulases (Cellulysin, Calbiochem) according to the method previously described (*126*). The obtained lignin-enriched CWRs (∼40 mg) were then dissolved in DMSO- *d_6_*/pyridine-*d_5_* [4:1 (v/v), 600 μL] and submitted for NMR analysis. NMR spectra for **Fig. 7B** were acquired at 300 K on a Bruker Biospin (Billerica, MA) NEO 700 MHz spectrometer equipped with a 5 mm QCI ^1^H/^31^P/^13^C/^15^N cryoprobe with inverse geometry (proton coils closest to the sample). Adiabatic HSQC NMR experiments were carried out using standard Bruker pulseprogram hsqcetgpsisp2.2 (*125*). Spectra were acquired from 11.6 to -0.6 ppm in F2 (^1^H) with 3448 datapoints (acquisition time, 100 ms) and 214 to -5.7 ppm in F1 (^13^C) with 618 increments (F1 acquisition time, 8 ms) of 64 scans with a 1 s interscan delay; d24 was 0.86 ms (⅛_J_, J = 145 Hz); The total experiment time was 13.5 h. Data processing and analysis were performed with the Bruker TopSpin software (Bruker Biospin) and the central DMSO solvent peaks (δ_C_/δ_H_: 39.5/2.49 ppm) were used as an internal reference. Processing to 1k × 1k datapoints typically used Gaussian apodization (LB = −0.5, GB = 0.001) in F2 and cosine-bell-squared in F1 with one level of linear prediction (32 coefficients). NMR correlation peak assignments, labeling, and color attributes are from prior work (*127*, *128*), aided by data in the NMR Database of Lignin and Cell Wall Model Compounds (*129*).

## Acknowledgement

We thank the Botany Greenhouse of University of Wisconsin-Madison for providing tissues of *Joinvillea ascendens* and other Poales plants, Dr. Clint Chapple (Purdue University) for providing the yeast strain, Jordan Teisher and the Missouri Botanical Garden for providing the *Puelia ciliata* specimen, Dr. Hong Ma and Weichen Huang (Pennsylvania State University) for searching CYP93G and CYP75B homologs from their *Puelioideae* genomes, Dr. Rick Amasino and Junko Maeda (University of Wisconsin-Madison) for providing the sorghum seeds, Dr. Matthew Moscou (USDA-ARS and University of Minnesota) for providing the genome for *Ecdeiocolea monostachya*, and Dr. Sebastian Bednarek (University of Wisconsin-Madison) for the use of the ultracentrifuge machine. The NMR data were collected using the Bruker Avance NEO 700 MHz instrument at the NMR facility of the Great Lakes Bioenergy Research Center (GLBRC) and Wisconsin Energy Institute (WEI) of the University of Wisconsin–Madison, which is based upon work supported by the Great Lakes Bioenergy Research Center, U.S. Department of Energy, Office of Science, Office of Biological and Environmental Research under Award Number DE- SC0018409, which also supported S.D.K and J.R. This work was supported by the U.S. National Science Foundation (NSF) Plant Genome Research Program (IOS-1836824) to H.A.M. Y.T-K. was partly supported by the Overseas research fellow of the Japan Society for the Promotion of Science (JSPS).

## Supplementary Materials

**Fig. S1.** Enzyme and TF families related to phenylpropanoid biosynthesis evolved at different times over the course of lignin evolution.

**Fig. S2.** Full co-expression modules from PTAL/TyrA1 and A3’H-C5’H representing the lignin networks in *Brachypodium distachyon* and Zea mays. PTAL: phenylalanine/tyrosine ammonia-lyase; TyrA: arogenate dehydrogenase; A3’H-C5’H: apigenin 3’-hydroxylase/chrysoeriol 5’- hydroxylase.

**Fig. S3.** Positive regulatory network of MYB transcription factors in grass lignin formation. MYB: myeloblastosis.

**Fig. S4.** Gene expression analysis by qRT-PCR demonstrates the existence of tyrosine-derived lignin pathway in *J. ascendens*.

**Fig. S5.** PMT and TyrAnc are syntenic across Poales species, whereas TyrA1 becomes syntenic within grasses. PMT: *p*-Coumaroyl-CoA monolignol transferase; TyrA: arogenate dehydrogenase. Fig. S6. FNSII is syntenic across grasses, as well as between pineapple and Joinvillea, but synteny

breaks down between grasses and non-grass Poales. FNSII: flavone synthase II.

**Fig. S7.** Monocot phylogeny of PMT/FMT/BAHD acyltransferases shows the conservation of co-expression between *J. ascendens* and grasses in most clades. PMT: *p*-Coumaroyl-CoA monolignol transferase; FMT: feruloyl-CoA monolignol transferase; BAHD: benzylalcohol O-acetyltransferase (BEAT), anthocyanin O-hydroxycinnamoyltransferase (AHCT), anthranilate *N*-hydroxycinnamoyl/benzoyltransferase (HCBT), and deacetylvindoline 4-*O*-acetyltransferase (DAT); PAT: p-coumaroyl-CoA Arabinose Transferase; FAT: feruloyl-CoA Arabinose Transferase

**Fig. S8.** HCT phylogeny shows the conservation of co-expression between *J. ascendens* and grasses. HCT: Hydroxycinnamoyl-CoA shikimate/quinate hydroxycinnamoyl transferase

**Fig. S9.** C3’H phylogeny shows the conservation of co-expression between *J. ascendens* and grasses. C3’H: p-coumaroyl ester 3-hydroxylase.

**Fig. S10.** CAD phylogeny shows the conservation of co-expression between *J. ascendens* and grasses. CAD: cinnamyl alcohol dehydrogenase.

**Fig. S11.** F5H phylogeny shows conservation of co-expression between *B. distachyon* and *J. ascendens*, but its loss in *Z. mays*. F5H: ferulate 5-hydroxylase.

**Fig. S12.** COMT phylogeny shows the conservation of co-expression between *J. ascendens* and grasses. COMT: caffeic acid *O*-methyltransferase.

**Fig. S13.** PAL phylogeny shows maintenance of co-expression between *J. ascendens* and grasses, but expansion of isoforms in grasses. PAL: phenylalanine ammonia-lyase.

**Fig. S14.** DHS phylogeny shows the conservation of co-expression between *J. ascendens* and grasses. DHS: 3-deoxy-D-arabinoheptulosonate-7-phosphate synthase.

**Fig. S15.** MYB phylogeny shows changes and conservation of co-expression between *J. ascendens*

and grass isoforms in different clades. MYB: myeloblastosis.

**Fig. S16.** TyrA phylogeny shows the maintenance of co-expression between *J. ascendens* and grass isoforms in one clade, and expansion of co-expression in grasses in another clade. TyrA: arogenate dehydrogenase.

**Fig. S17.** CCoAOMT phylogeny shows change of co-expression between *J. ascendens* and grass isoforms. CCoAOMT: caffeoyl-CoA 3-O-methyltransferase.

**Fig. S18.** CCR phylogeny shows the conservation of co-expression between *J. ascendens* and grasses in one clade and the gain of co-expression in grasses in a separate clade. CCR: cinnamoyl-CoA reductase.

**Fig. S19.** CHS phylogeny shows a gain of co-expression in *B. distachyon*. CHS: chalcone synthase.

**Fig. S20.** CHIL phylogeny shows the expansion of co-expression in *B. distachyon* enzymes from *J. ascendens*. CHIL: chalcone isomerase-like.

**Fig. S21.** CHI phylogeny shows a gain of co-expression in grasses. CHI: chalcone isomerase. Fig. S22. CYP450 phylogeny shows the expansion of co-expression for two clades of tricin- producing enzymes from *J. ascendens* to grasses. CYP450: cytochrome P450

**Fig. S23.** C3H/APX phylogeny shows a gain of co-expression in *B. distachyon*. C3H/APX: coumarate 3-hydroxylase/ascorbate peroxidase.

Fig. S24A. ADT1 phylogeny shows both conservation of co-expression between *Z. mays* and *J. ascendens* and co-expression loss in grasses. ADT: arogenate dehydratase.

Fig. S24B. ADT2 phylogeny shows loss of co-expression in grasses. ADT: arogenate dehydratase.

**Fig. S25.** CSE phylogeny shows loss of genes in grasses. CSE: caffeoyl shikimate esterase.

**Fig. S26.** C4H phylogeny shows change of co-expression between *J. ascendens* and grass isoforms. C4H: cinnamate 4-hydroxylase.

Fig. S27A. 4CL1-2 phylogeny shows conservation of co-expression between grasses and *J. ascendens* as well as a loss of co-expression in grasses. 4CL: 4-coumarate:CoA ligase.

Fig. S27B. 4CL3 phylogeny has no co-expression in *J. ascendens* or grasses. 4CL: 4- coumarate:CoA ligase.

**Fig. S28.** FNSII and F2H activities of FNSII and F2H from *Brachypodium distachyon* and their closest homolog in *Joinvillea ascendens* (JaCYP93G1), suggesting the JaCYP93G1 possess FNSII activity. FNSII: flavone synthase II; F2H: flavanone 2-hydroxylase.

**Fig. S29.** A3’H, C5’H, and F3’H activities of A3’H-C5’H and F3’H from *Brachypodium distachyon* and their closest homolog in *Joinvillea ascendens* (JaCYP75B), suggesting single *J. ascendens* CYP75B enzyme lack C5’H activity. A3’H-C5’H: apigenin 3’-hydroxylase/chrysoeriol 5’- hydroxylase; F3’H: flavonoid 3’-hydroxylase.

**Fig. S30.** Accumulation of tricin intermediates and flavone C-glucosides determined by LC-MS analysis of the MeOH extract. Chromatograms of each compound for the selected species (A) and normalized LC-MS peak intensity (B) are shown.

**Fig. S31.** Grass lignin network evolved in a stepwise manner.

**Fig. S32.** Golden gate constructs for recombinant protein induction in *S. cerevisiae* WAT11.

**table S1.** Phenylpropanoid gene counts for green plants

**table S2.** Phenylpropanoid gene counts monocots

**table S3.** *B. distachyon* co-expression clusters

**table S4.** GO and pathway cluster enrichment

**table S5.** Percent cluster overlap across lignin co-expression clusters significance percentiles

**table S6.** *Z. mays* co-expression clusters

**table S7.** *J. ascendens* co-expression clusters

**table S8.** Green plant genome list

**table S9.** Monocot genome list

**table S10.** Orthogroup and tree model list

**table S11.** Primers used in this study

**table S12.** RNAseq read count and quality score

## Supplementary Materials for

**Figure S1.**
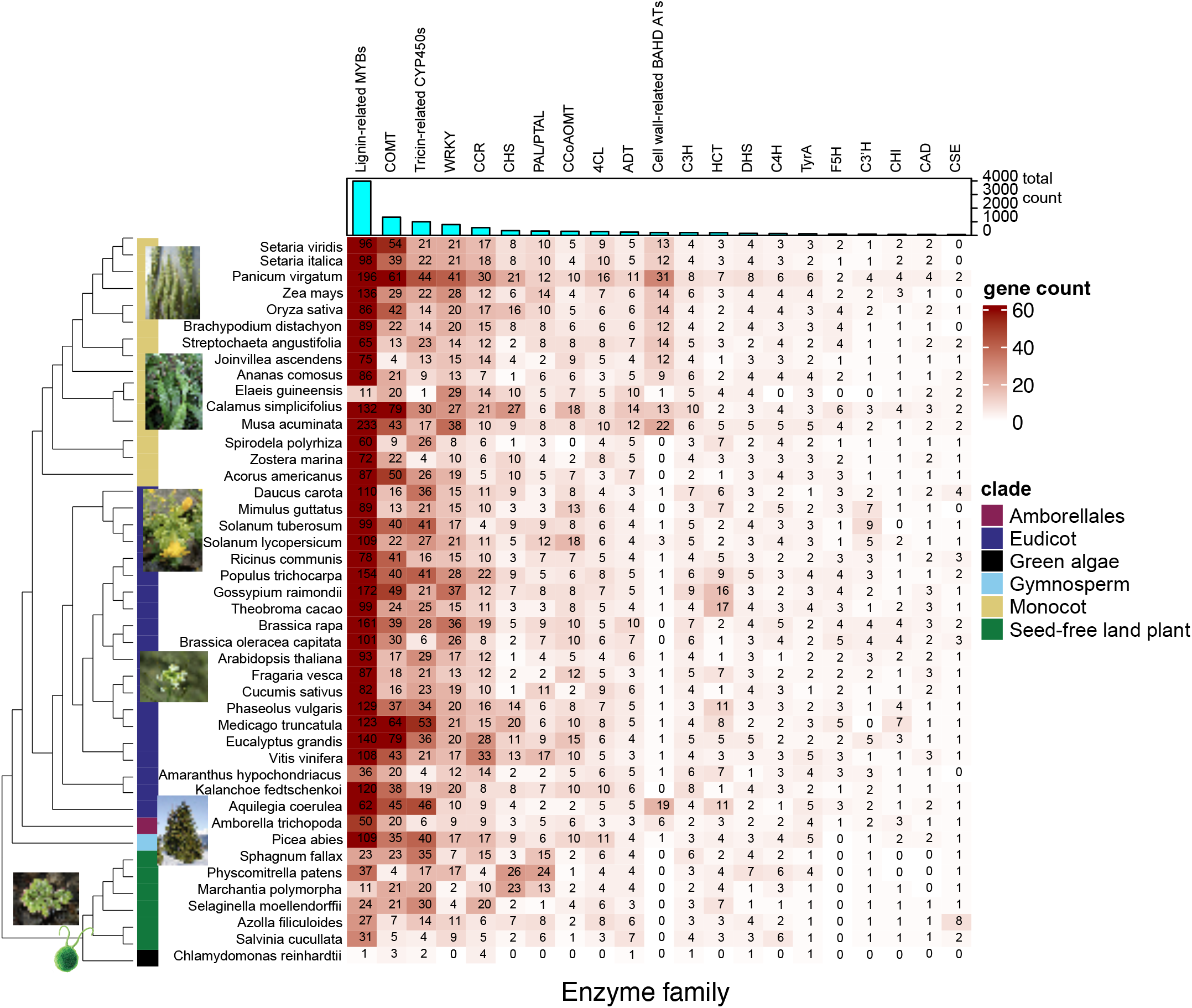
Enzyme and TF families related with phenylpropanoid biosynthesis evolve at different times over the course of lignin evolution. The heatmap represents the number of gene isoforms of an enzyme or transcription factor family belonging to the lignin pathway (column) in a given species (row). The darker the red color, the more isoforms are present. The bar graph at the top indicates the total number of isoforms for an enzyme family in all species. The color bars on the phylogeny denote different taxonomic clades of plants and algae as indicated.

**Figure S2.**
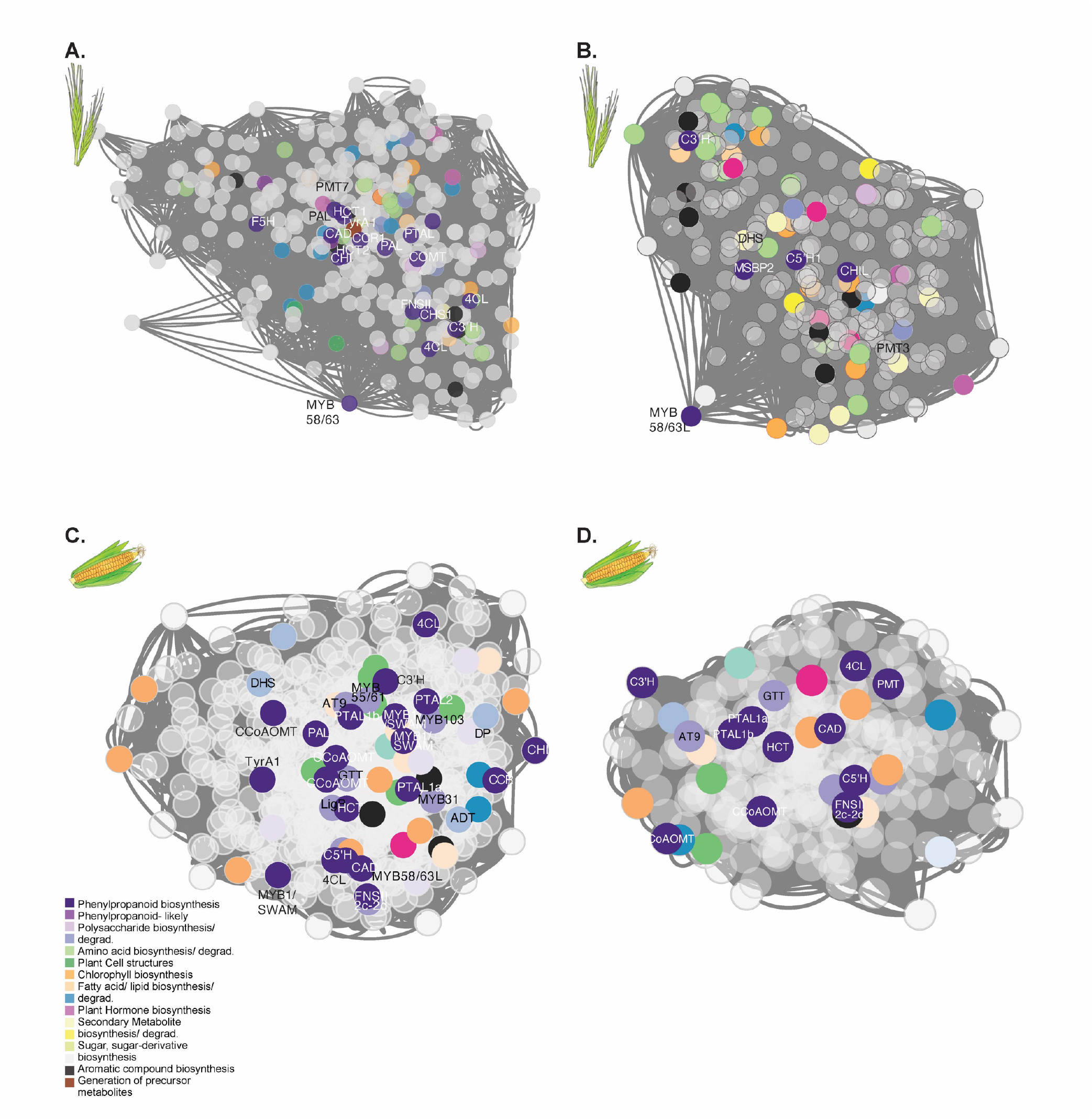
Full co-expression modules from PTAL/TyrA1 and A3’H-C5’H representing the lignin networks in *Brachypodium distachyon* and *Zea mays*. **A.** Co-expression of the lignin network in *B. distachyon* based on PTAL and TyrA1 genes, where each node is a gene and connections are representative of the distances between genes based on Spearman’s mutual rank. Known lignin enzymes are in dark purple and labeled with their enzyme abbreviation. Other colors correspond to pathways in different categories. Genes in grey belong to an unknown pathway. Module size is 460 genes. **B.** Co-expression of the lignin network in *B. distachyon* based on A3’H-C5’H. Nodes, distances, and colors are the same as above. **C.** Co-expression of the lignin network in *Z. mays* based on PTAL and TyrA1 genes. Each node is a gene and connections are representative of the distances between genes based on Spearman’s mutual rank. Known lignin enzymes are in dark purple and labeled with their enzyme abbreviation. Other colors correspond to pathways in different categories. Genes in grey belong to an unknown pathway. Module size is 600 genes. **D.** Co-expression of the lignin network in *Z. mays* based on A3’H-C5’H. Nodes, distances, and colors are the same as above.

**Figure S3.**
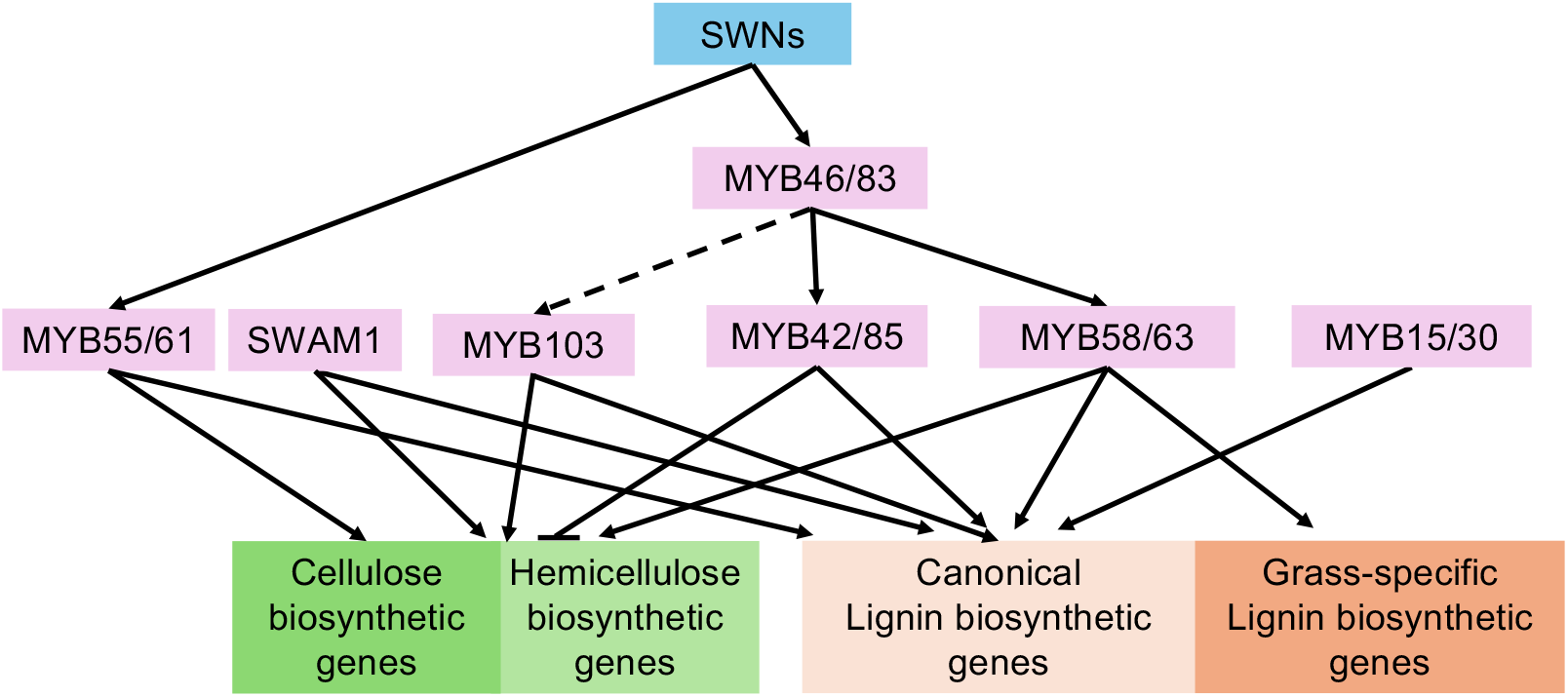
Positive regulatory network of MYB transcription factors in grass lignin formation. Positive transcriptional regulation is represented by arrows, and negative transcriptional regulation is shown by bars at the ends of lines. The scheme is based on Rao and Dixon (2018) and Miyamoto et al. (2020), with some modifications. SWN, secondary wall-associated NAC.

**Figure S4.**
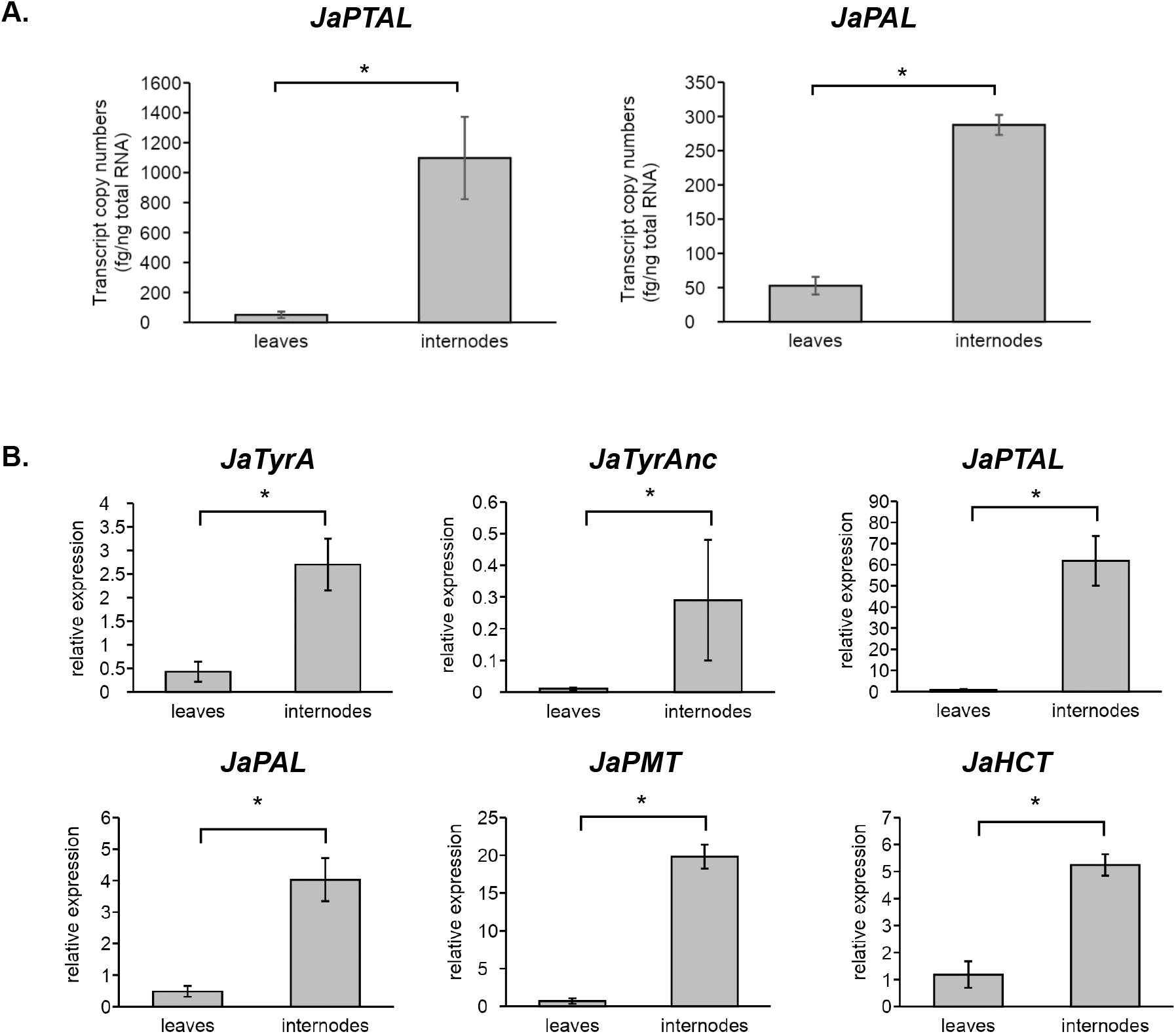
Gene expression analysis by qRT-PCR demonstrates the existence of tyrosine-derived lignin pathway in *J. ascendense*. **A.** Transcript copy numbers of *JaPTAL* and *JaPAL* in 8 month old leaves and internodes. **B.** RT-qPCR quantification, relative to the expression of *JaUBI10*, revealed that transcripts of JaTyrAs and other lignin pathway genes were much more abundant in the internodes compared to the leaves. Values are mean ± SD (*n* = 3). Asterisks indicate significant difference (Student’s *t*-test; *p* < 0.05)

**Figure S5.**
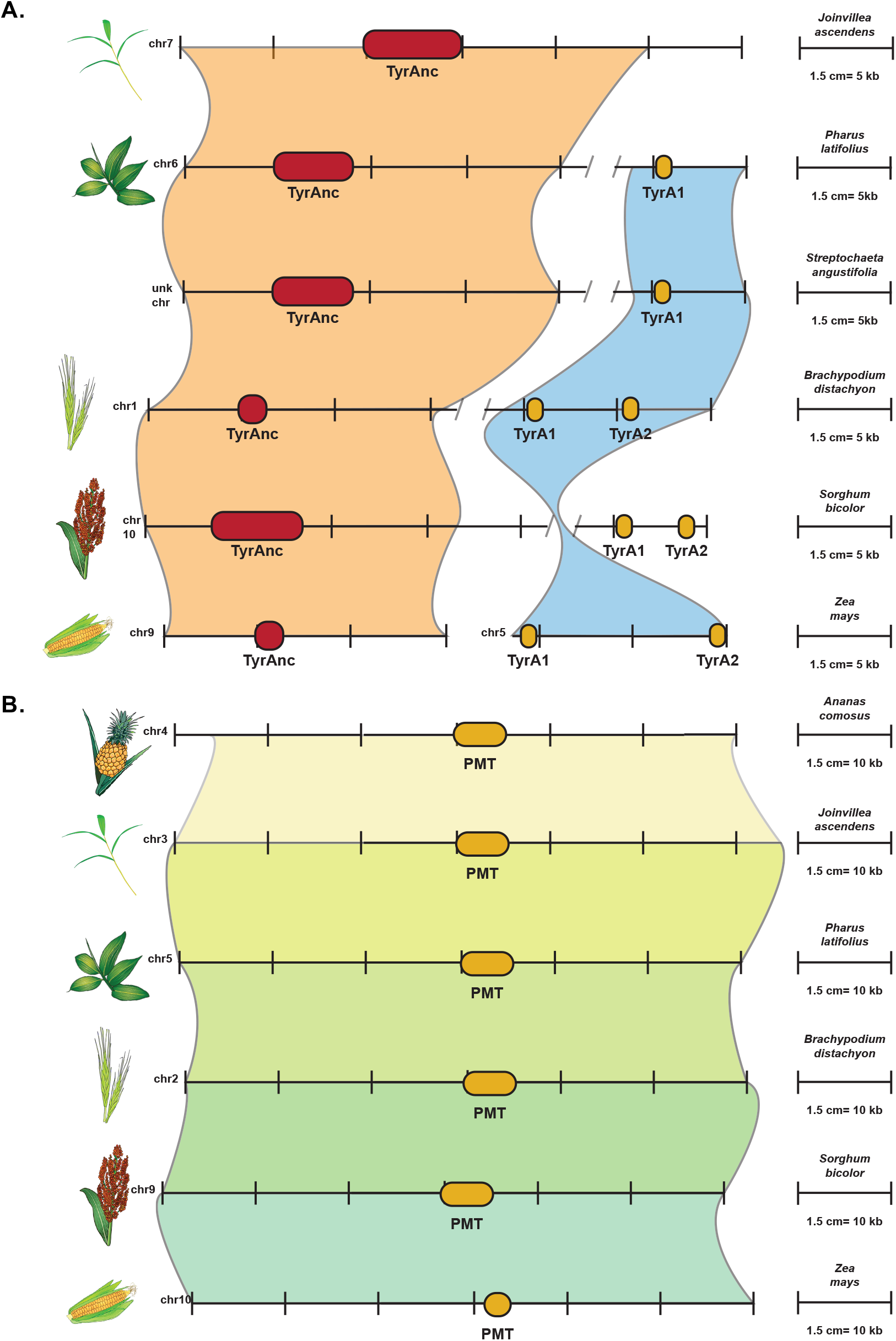
PMT and TyrAnc are syntenic across Poales species, while TyrA1 becomes syntenic within grasses. **A.** Synteny of TyrA across Poales species showing differences between the isoforms TyrA1 and TyrAnc. TyrAnc is shown in red, while TyrA1/2 is shown in yellow. Relative distance on each chromosome is given by the scale for each species. **B.** Synteny of PMT across Poales species. Relative distance on each chromosome is given by the scale for each species.

**Figure S6.**
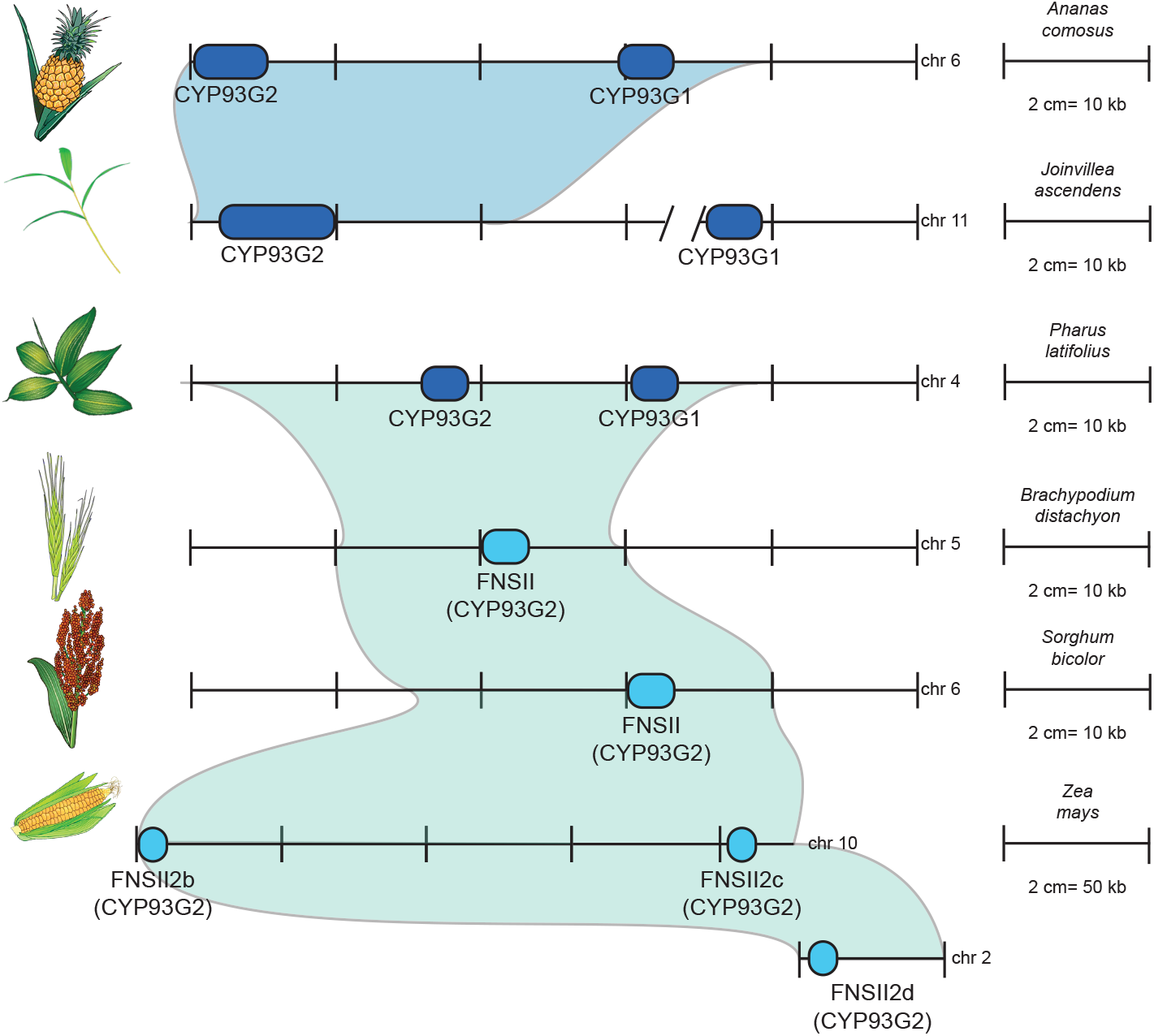
FNSII is syntenic across grasses, as well as between pineapple and Joinvillea, but synteny breaks down between grasses and non-grass Poales. Synteny of CYP93G across Poales species are shown. Dark blue genes are CYP93Gs with either unknown activity or dual FNSII/F2H activity. Light blue genes are CYP93Gs with only FNSII activity. Relative distance on each chromosome is given by the scale for each species.

**Figure S7.**
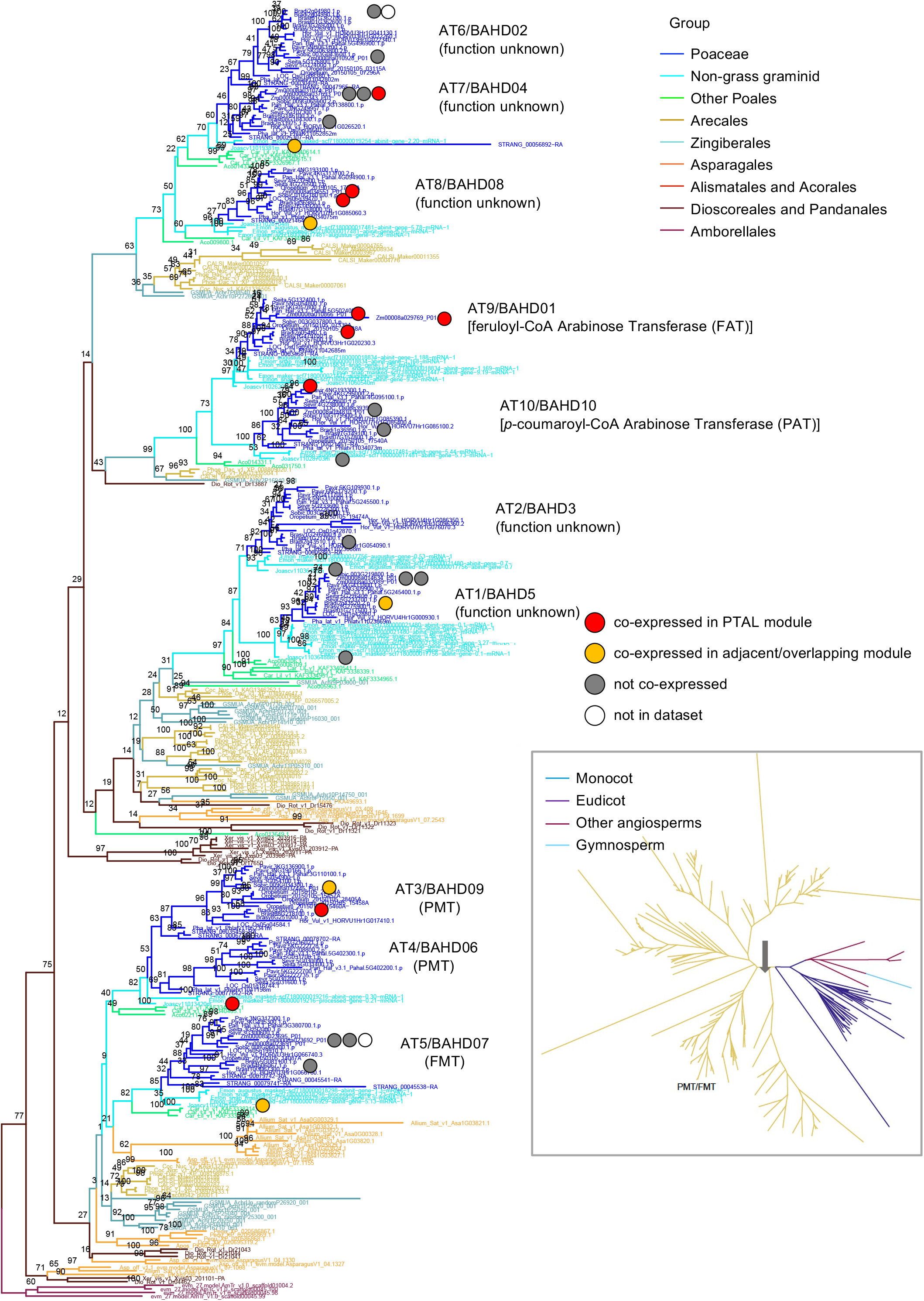
Monocot phylogeny of PMT/FMT/BAHD acyltransferases shows the conservation of co-expression between *J. ascendens* and grasses in most clades. The main tree is built using monocot species while the inset tree is built using species from all plant clades. The grey arrow in the inset tree shows where the base of the monocot tree starts.

**Figure S8.**
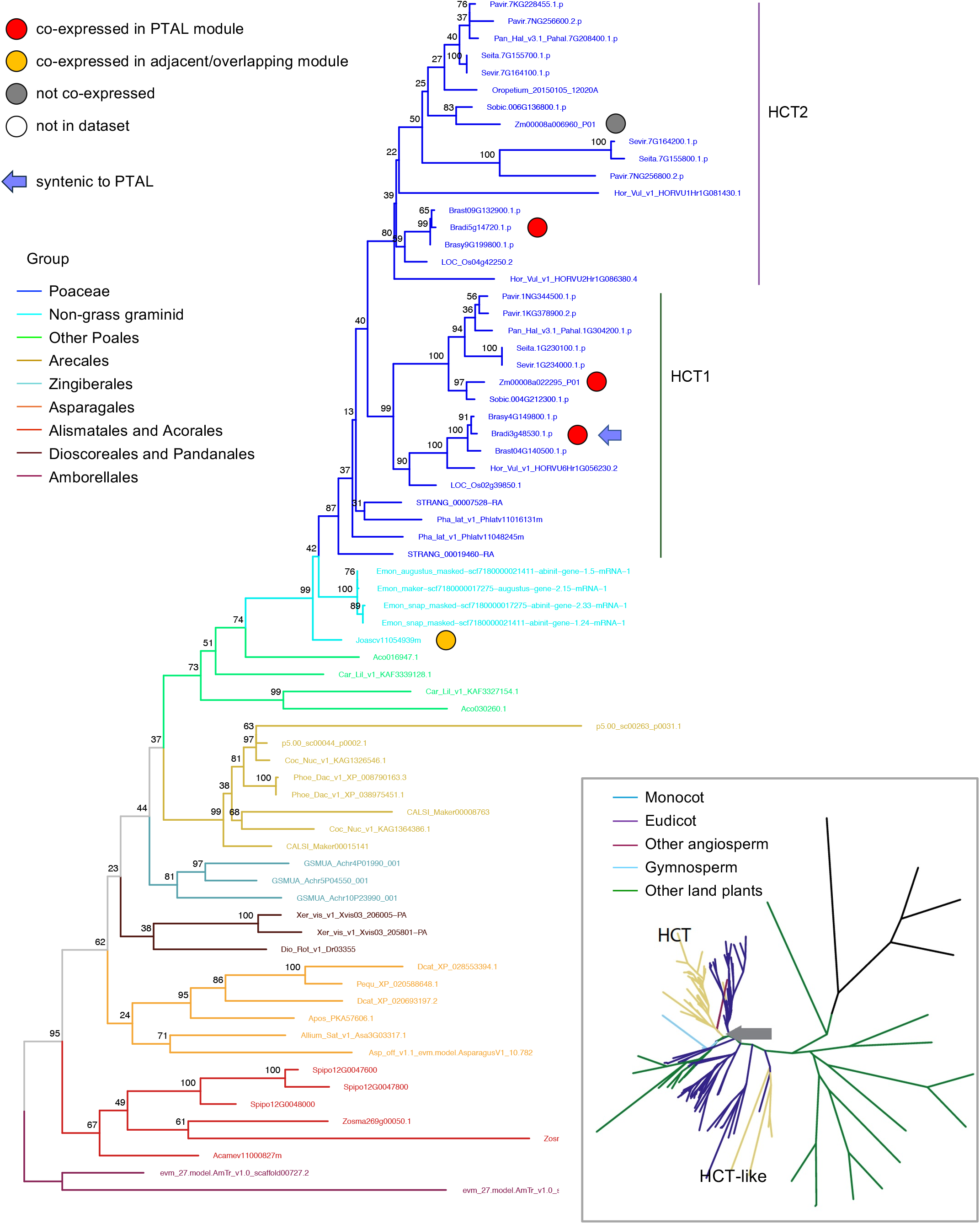
HCT phylogeny shows the conservation of co-expression between *J. ascendens* and grasses. The main tree is built using monocot species while the inset tree is built using species from all plant clades. The grey arrow in the inset tree shows where the base of the monocot tree starts.

**Figure S9.**
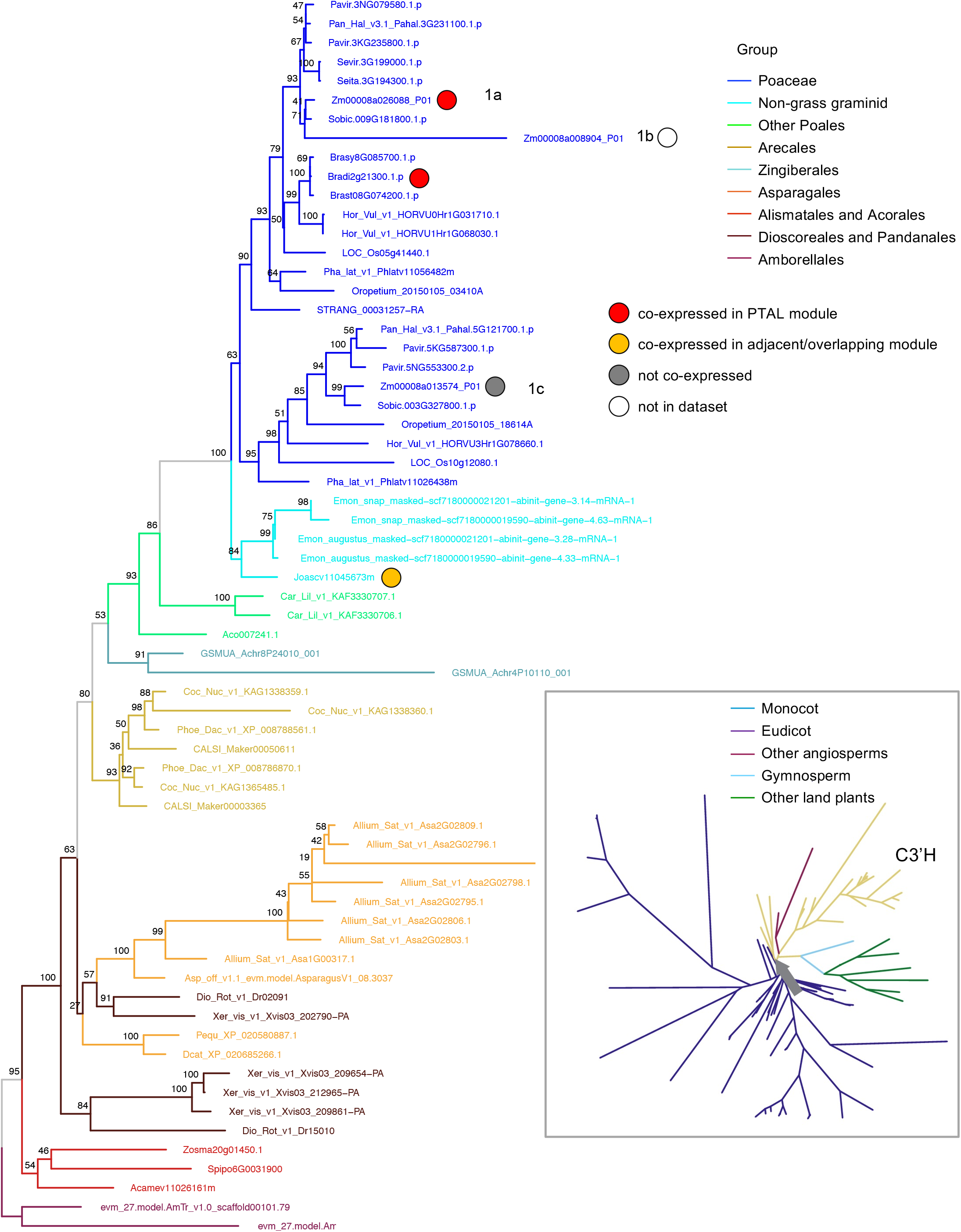
C3′H phylogeny shows the conservation of co-expression between *J. ascendens* and grasses. The main tree is built using monocot species while the inset tree is built using species from all plant clades. The grey arrow in the inset tree shows where the base of the monocot tree starts.

**Figure S10.**
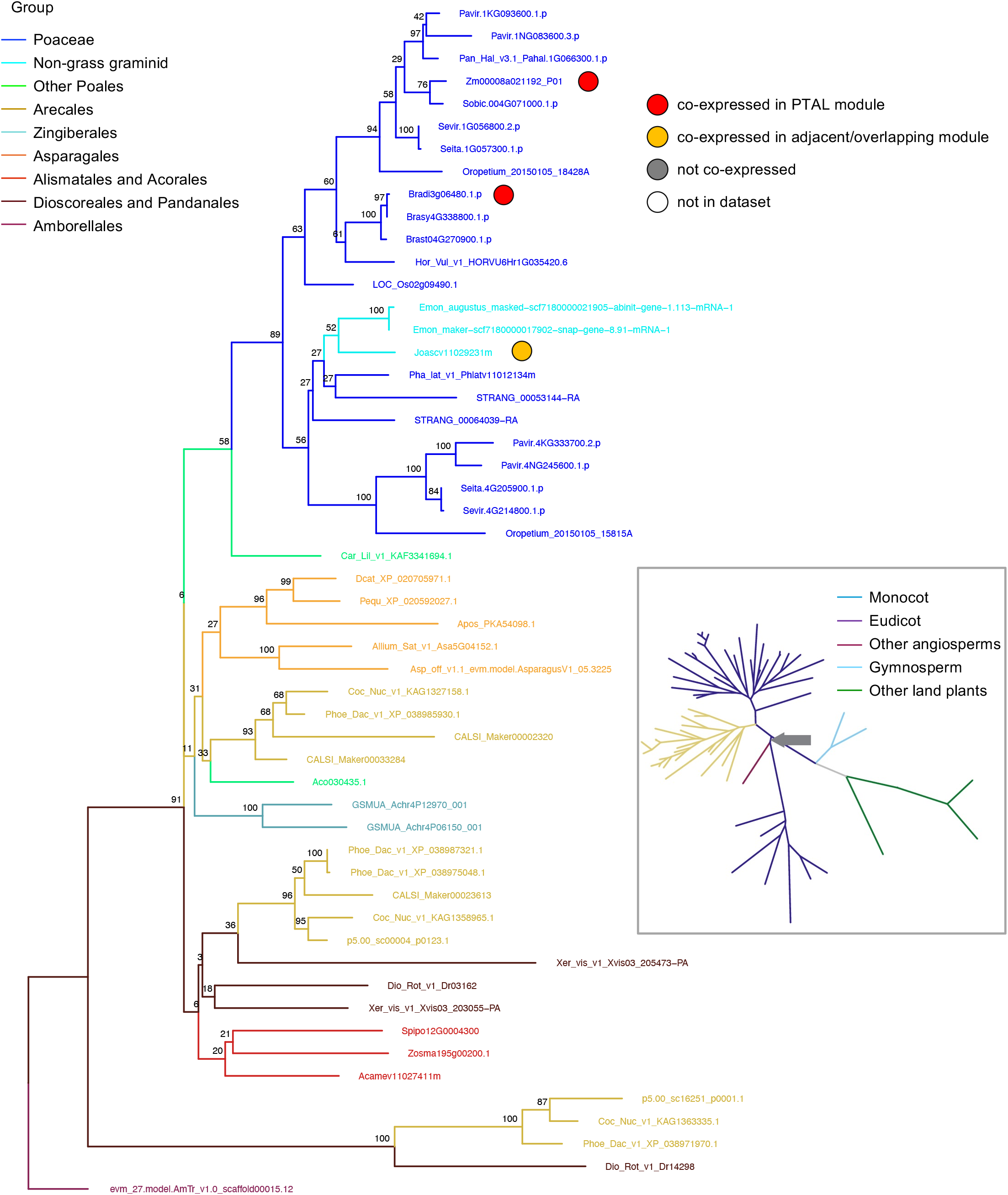
CAD phylogeny shows the conservation of co-expression between *J. ascendens* and grasses. The main tree is built using monocot species while the inset tree is built using species from all plant clades. The grey arrow in the inset tree shows where the base of the monocot tree starts.

**Figure S11.**
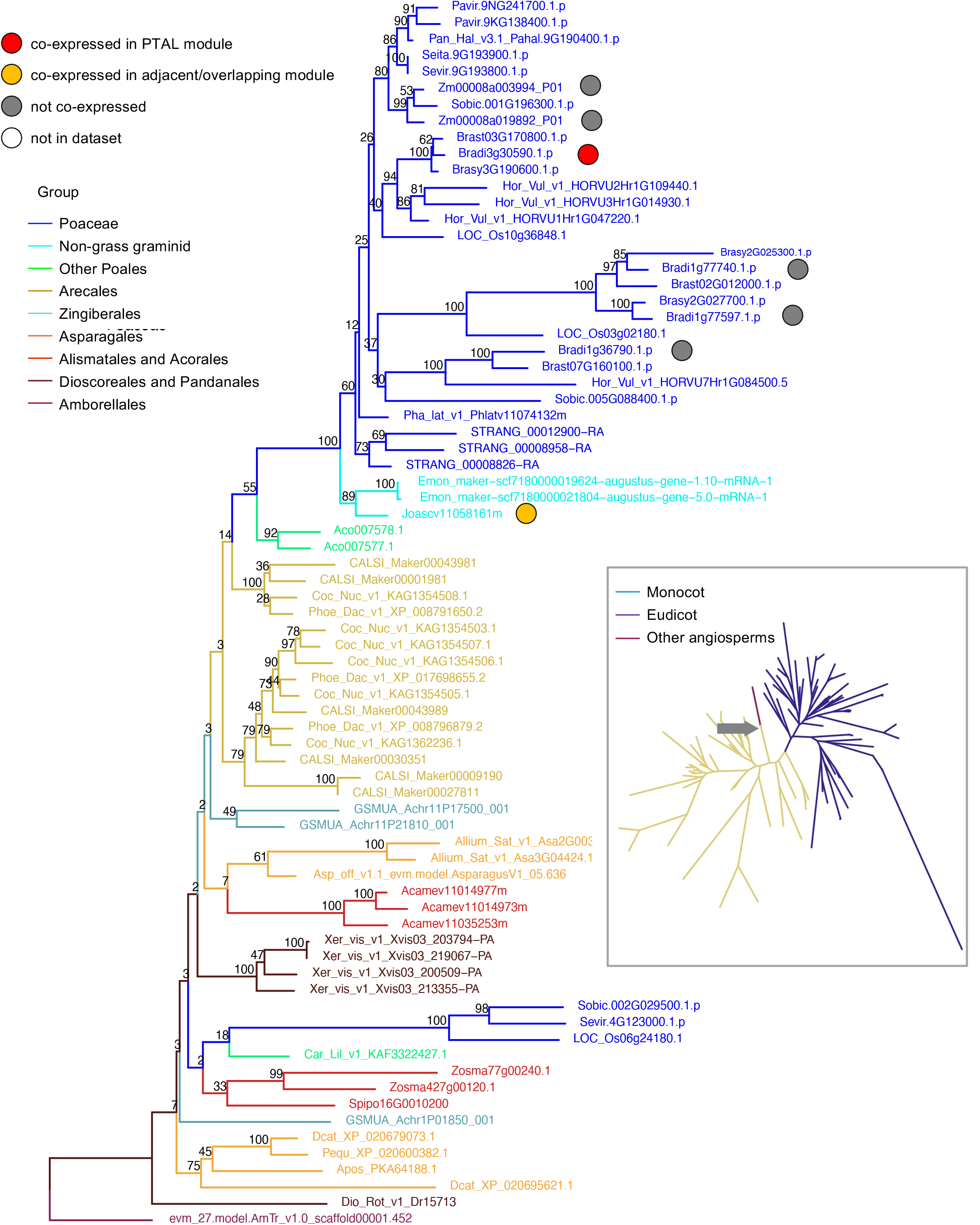
F5H phylogeny shows conservation of co-expression between *B. distachyon* and *J. ascendens,* but loss in *Z. mays*. The main tree is built using monocot species while the inset tree is built using species from all plant clades. The grey arrow in the inset tree shows where the base of the monocot tree starts.

**Figure S12.**
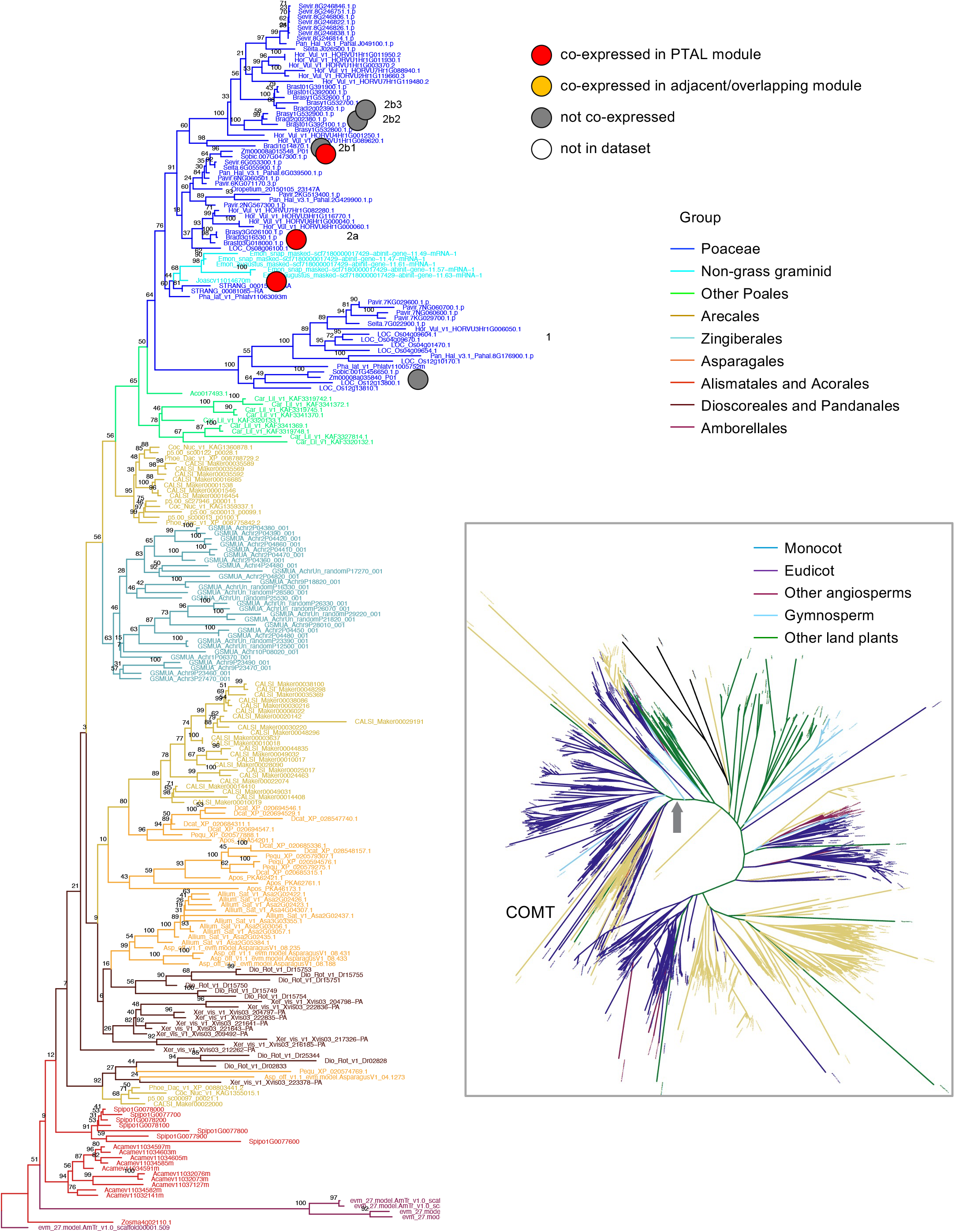
COMT phylogeny shows the conservation of co-expression between *J. ascendens* and grasses. The main tree is built using monocot species while the inset tree is built using species from all plant clades. The grey arrow in the inset tree shows where the base of the monocot tree starts.

**Figure S13.**
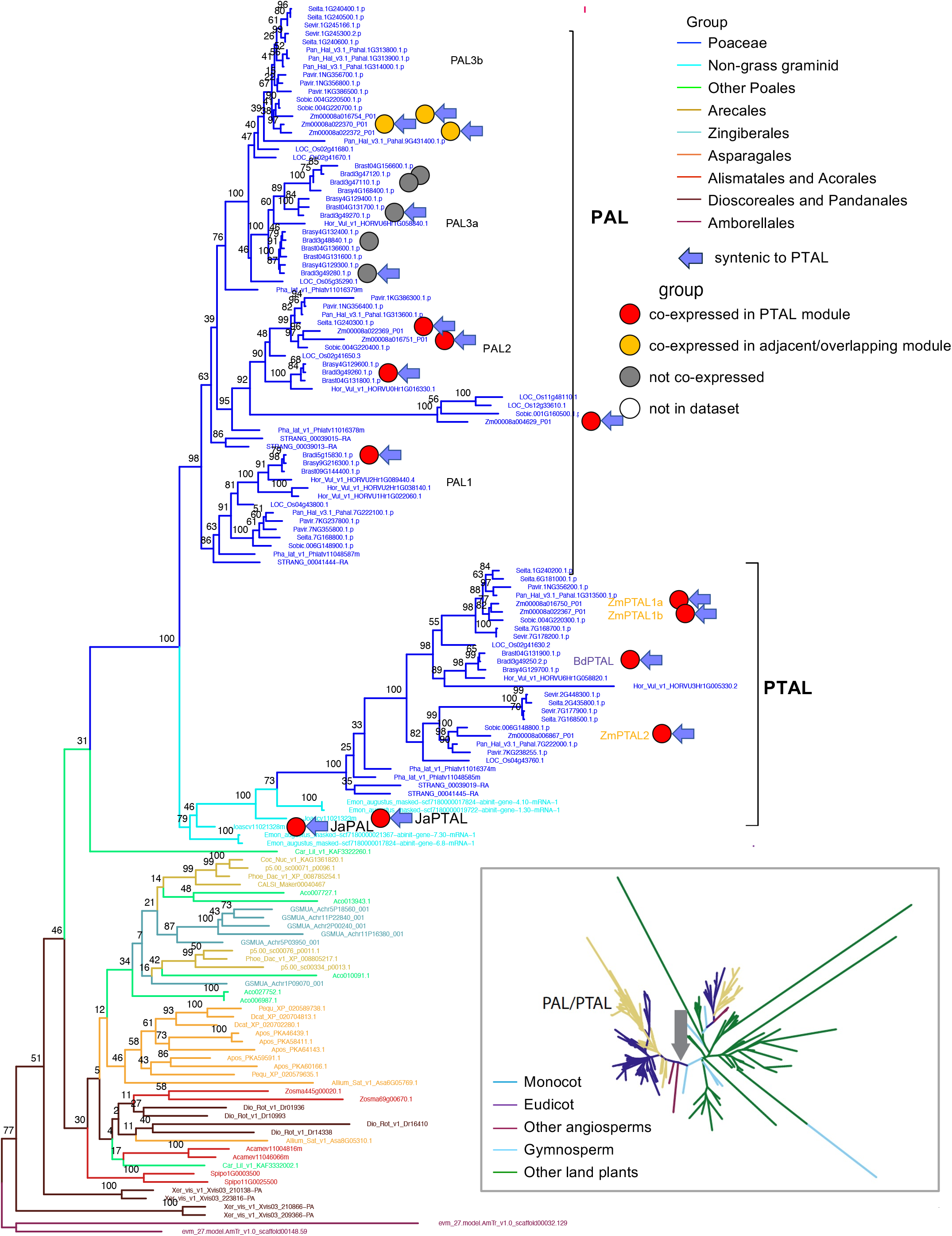
PAL phylogeny shows maintenance of co-expression between *J. ascendens* and grasses, but expansion of isoforms in grasses. The main tree is built using monocot species while the inset tree is built using species from all plant clades. The grey arrow in the inset tree shows where the base of the monocot tree starts.

**Figure S14.**
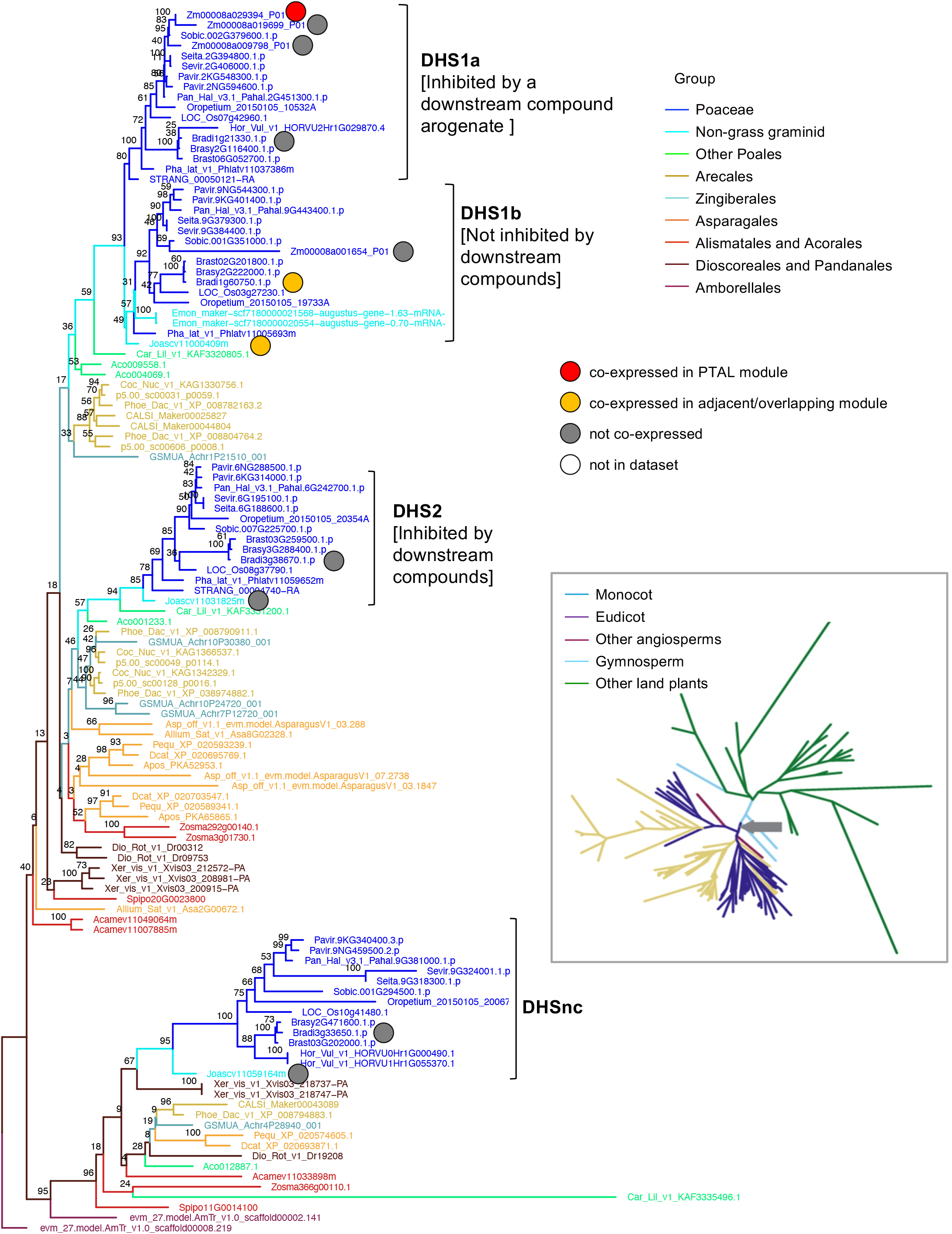
DHS phylogeny shows the conservation of co-expression between *J. ascendens* and grasses. The main tree is built using monocot species while the inset tree is built using species from all plant clades. The grey arrow in the inset tree shows where the base of the monocot tree starts.

**Figure S15.**
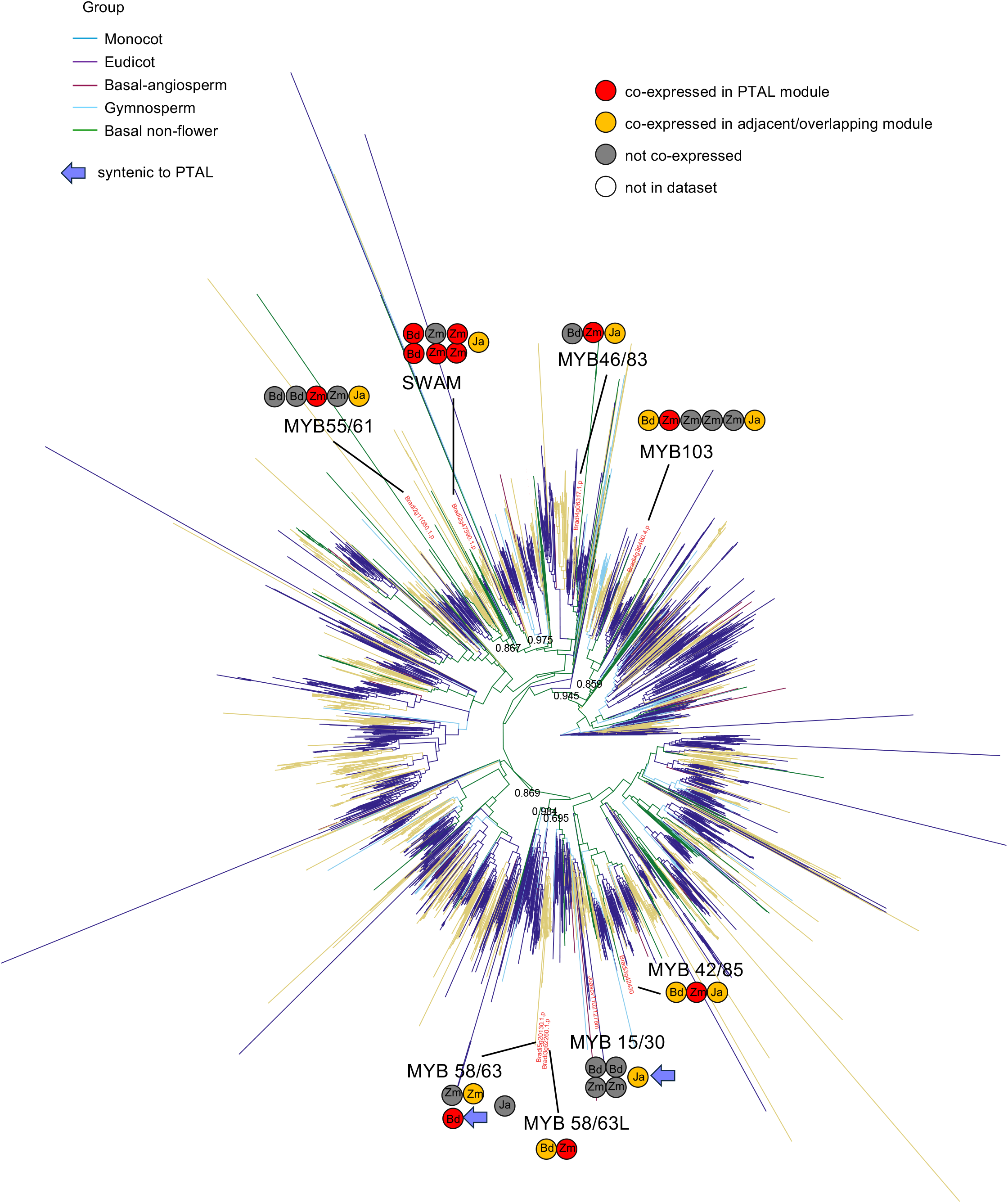
MYB phylogeny shows changes and conservation of co-expression between *J. ascendens* and grass isoforms in different clades.

**Figure S16.**
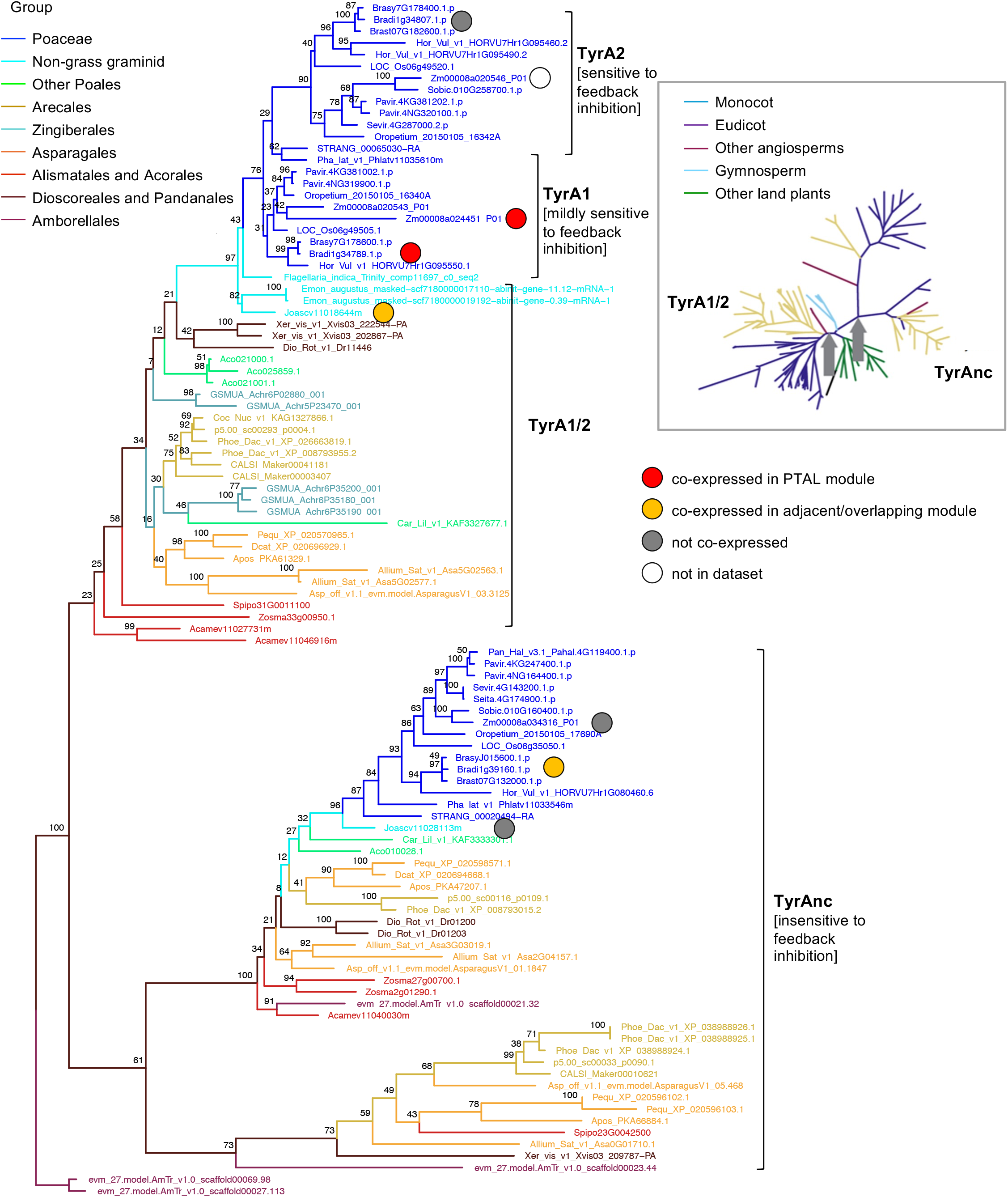
TyrA phylogeny shows the maintenance of co-expression between *J. ascendens* and grass isoforms in one clade, and expansion of co-expression in grasses in another clade. The main tree is built using monocot species while the inset tree is built using species from all plant clades. The grey arrow in the inset tree shows where the base of the monocot tree starts.

**Figure S17.**
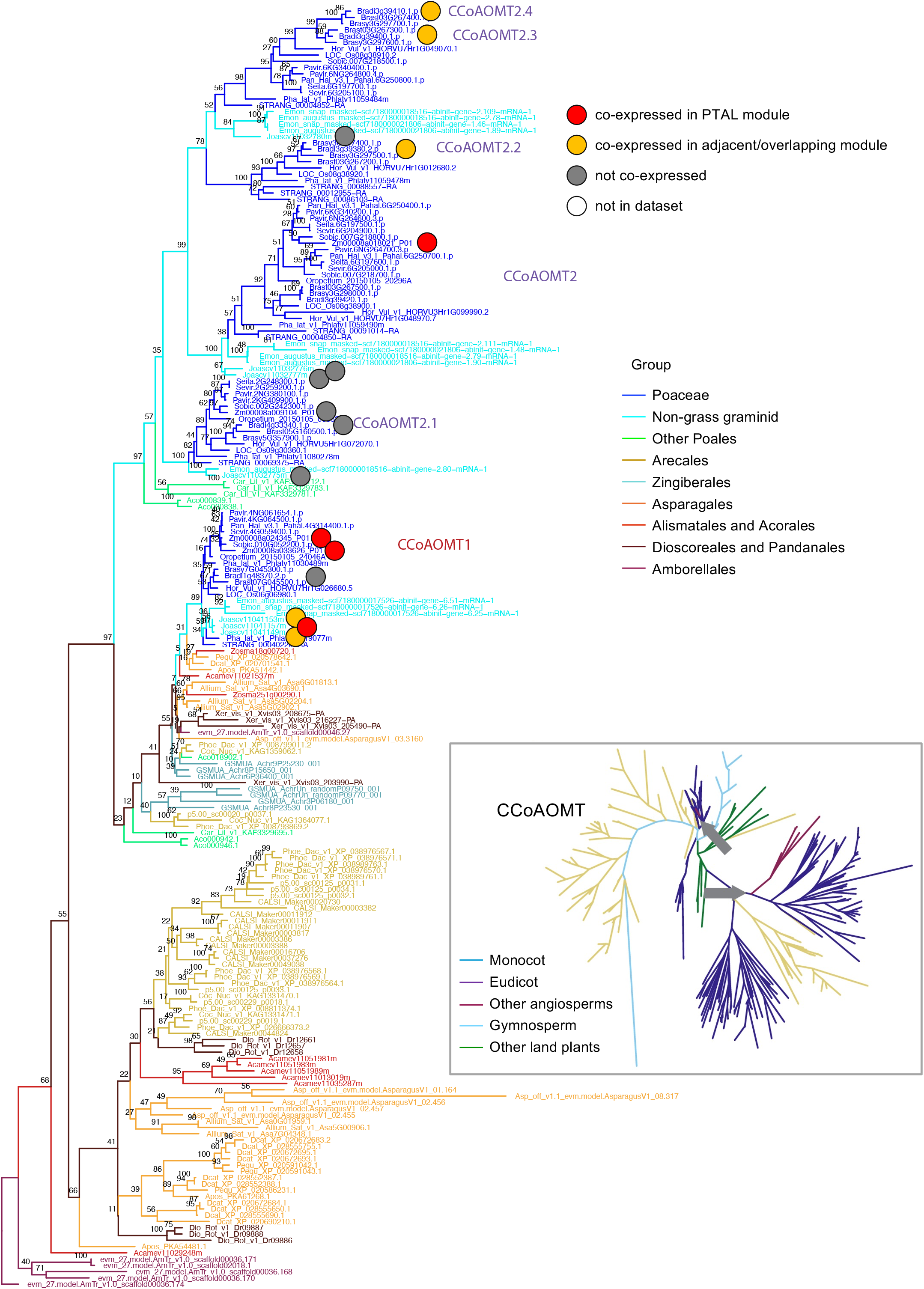
CCoAOMT phylogeny shows change of co-expression between *J. ascendens* and grass isoforms. The main tree is built using monocot species while the inset tree is built using species from all plant clades. The grey arrow in the inset tree shows where the base of the monocot tree starts.

**Figure S18.**
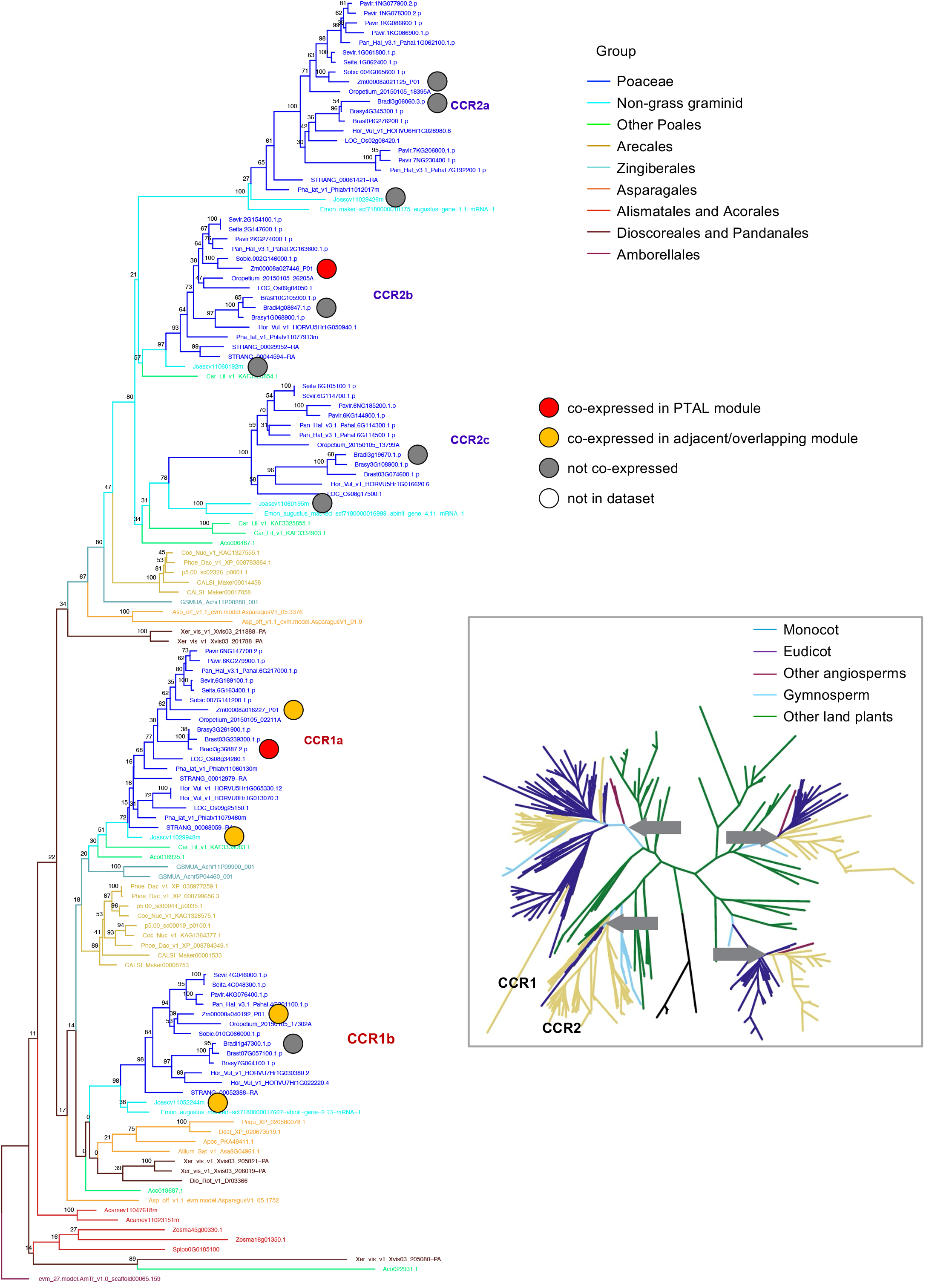
CCR phylogeny shows the conservation of co-expression between *J. ascendens* and grasses in one clade and the gain of co-expression in grasses in a separate clade. The main tree is built using monocot species while the inset tree is built using species from all plant clades. The grey arrow in the inset tree shows where the base of the monocot tree starts.

**Figure S19.**
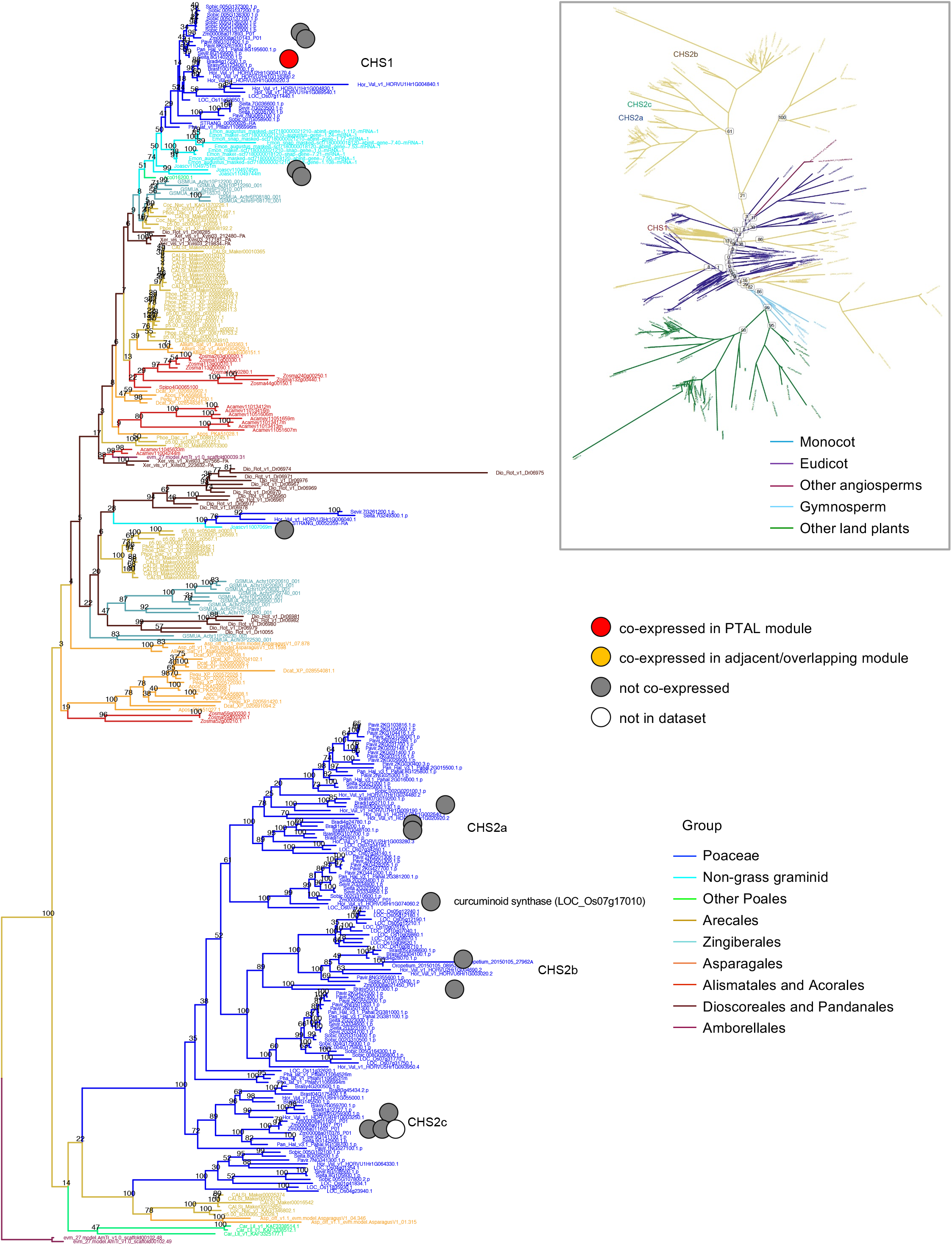
CHS phylogeny shows a gain of co-expression in *B. distachyon*. The main tree is built using monocot species while the inset tree is built using species from all plant clades. The grey arrow in the inset tree shows where the base of the monocot tree starts.

**Figure S20.**
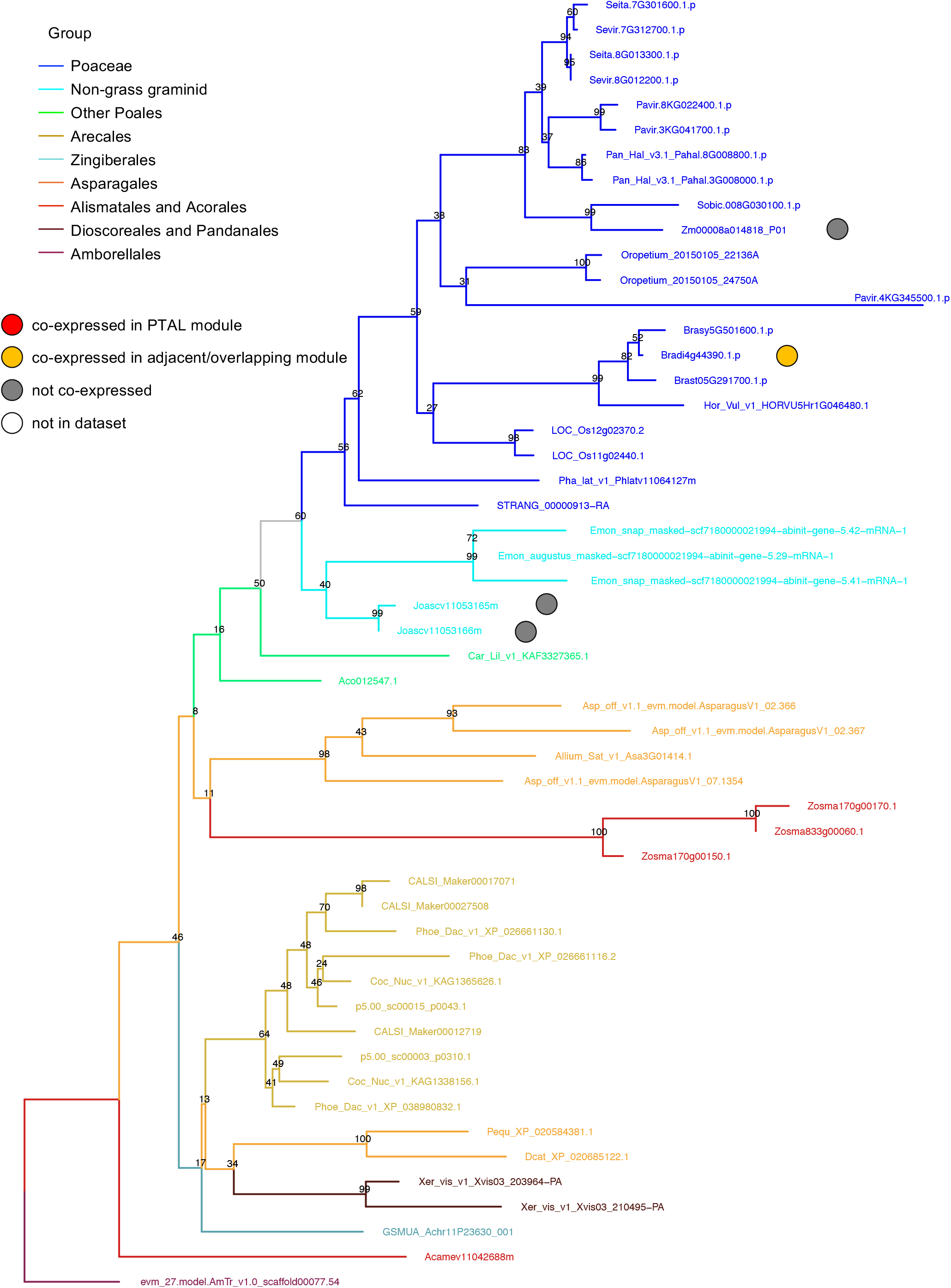
CHIL phylogeny shows the expansion of co-expression in *B. distachyon* enzymes from *J. ascendens*. The tree is built using monocot species.

**Figure S21.**
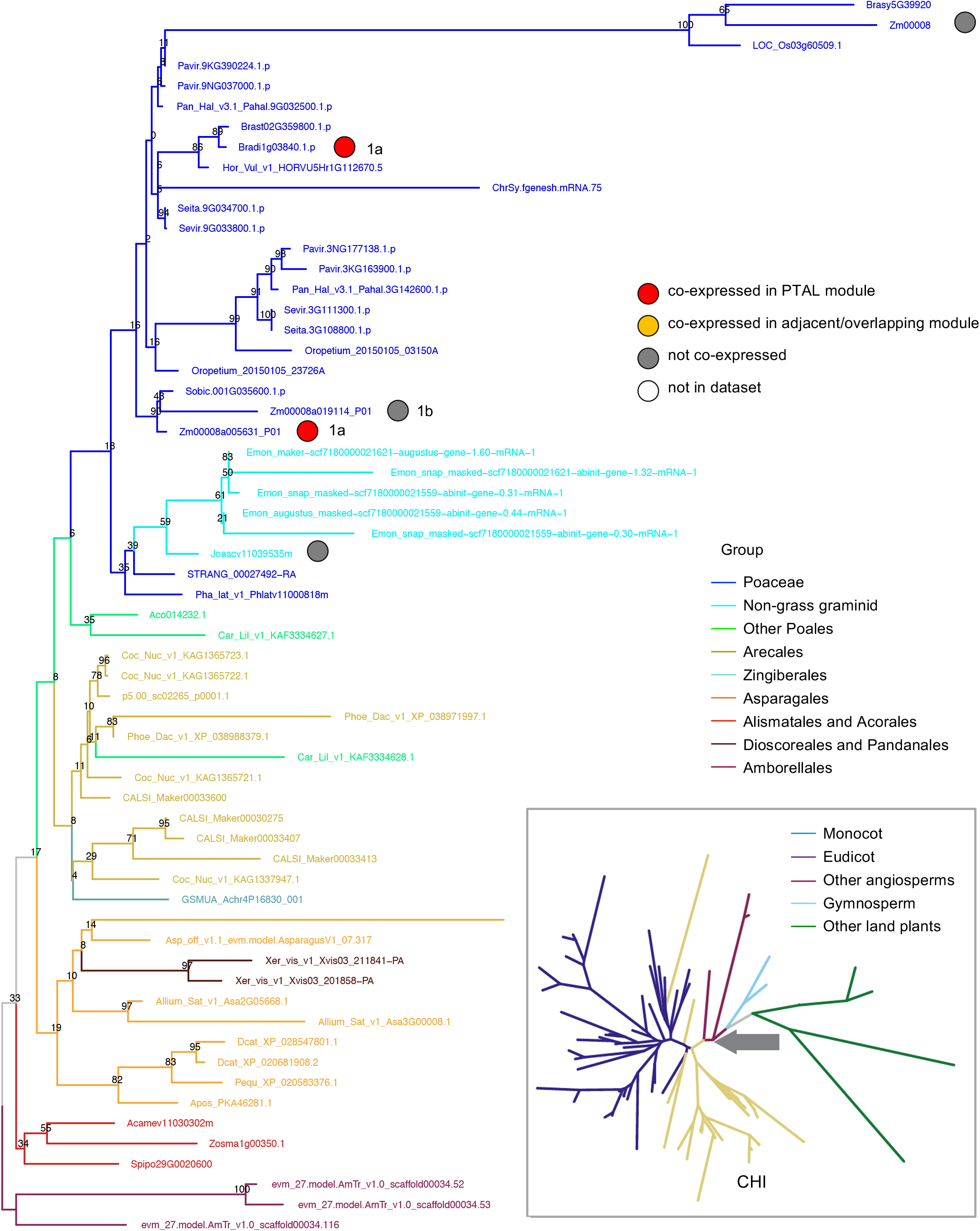
CHI phylogeny shows a gain of co-expression in grasses. The main tree is built using monocot species while the inset tree is built using species from all plant clades. The grey arrow in the inset tree shows where the base of the monocot tree starts.

**Figure S22.**
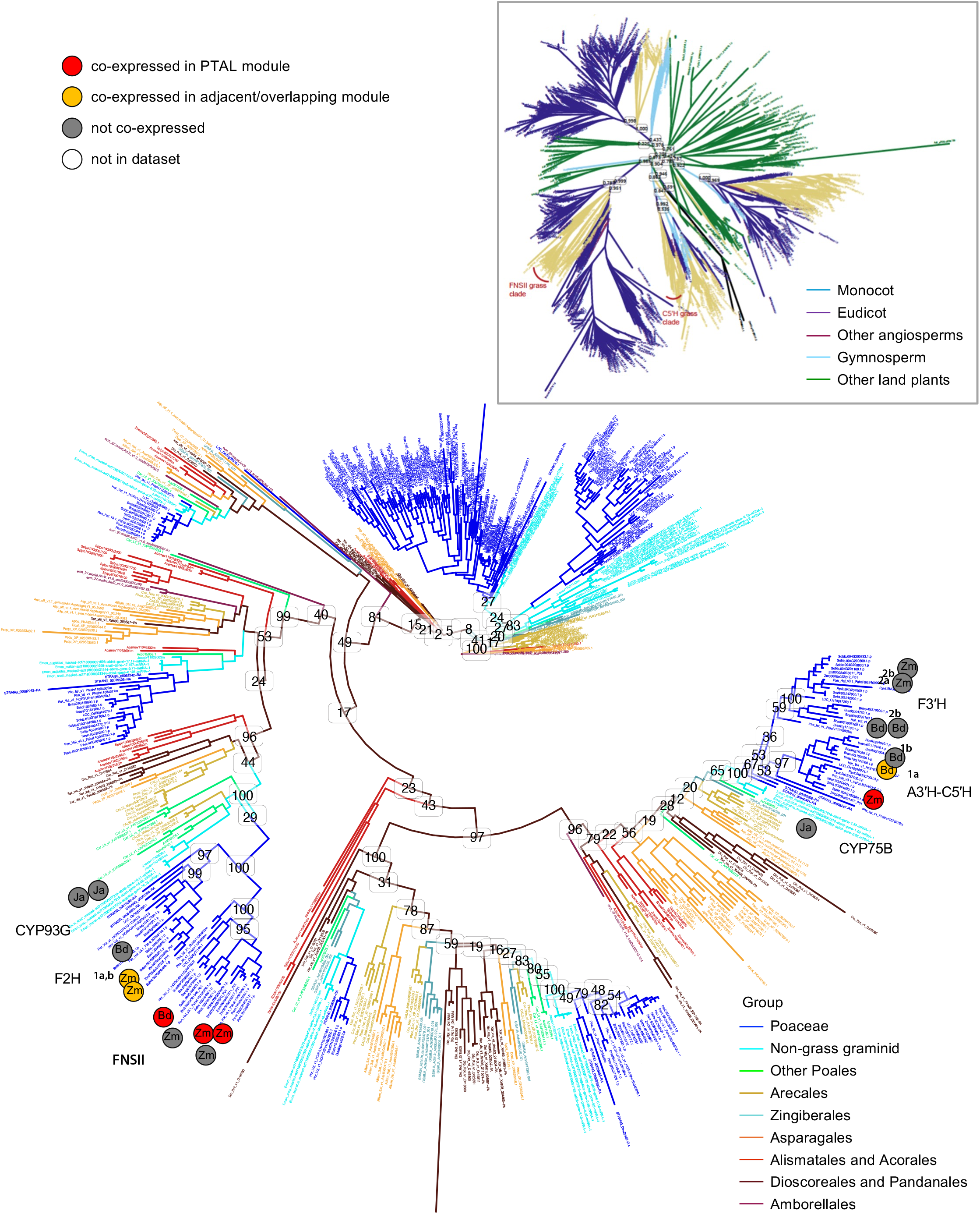
CYP450 phylogeny shows the expansion of co-expression for two clades of tricin-producing enzymes from *J. ascendens* to grasses. The main tree is built using monocot species while the inset tree is built using species from all plant clades.

**Figure S23.**
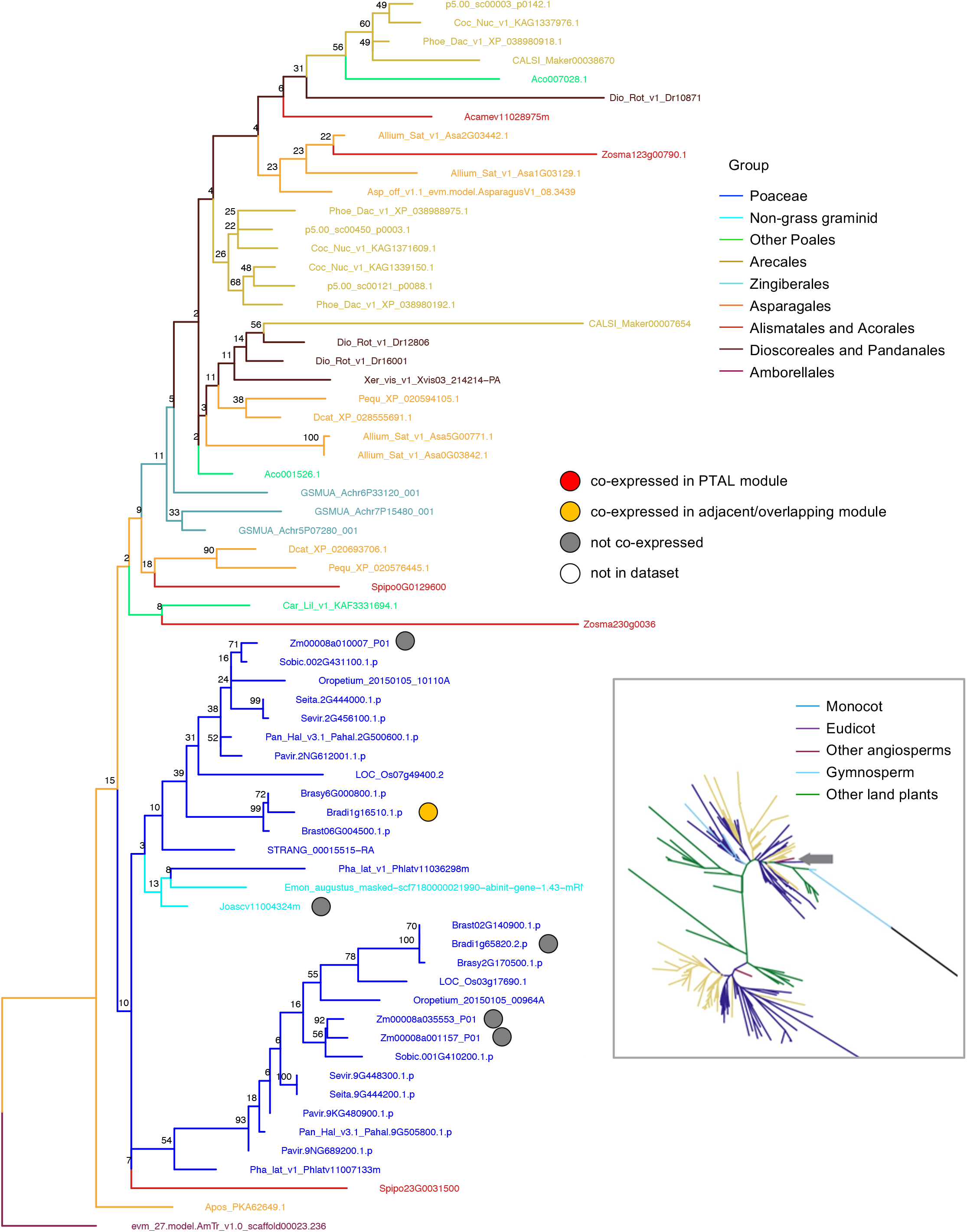
C3H/APX phylogeny shows a gain of co-expression in *B. distachyon*. The main tree is built using monocot species while the inset tree is built using species from all plant clades. The grey arrow in the inset tree shows where the base of the monocot tree starts.

**Figure S24A.**
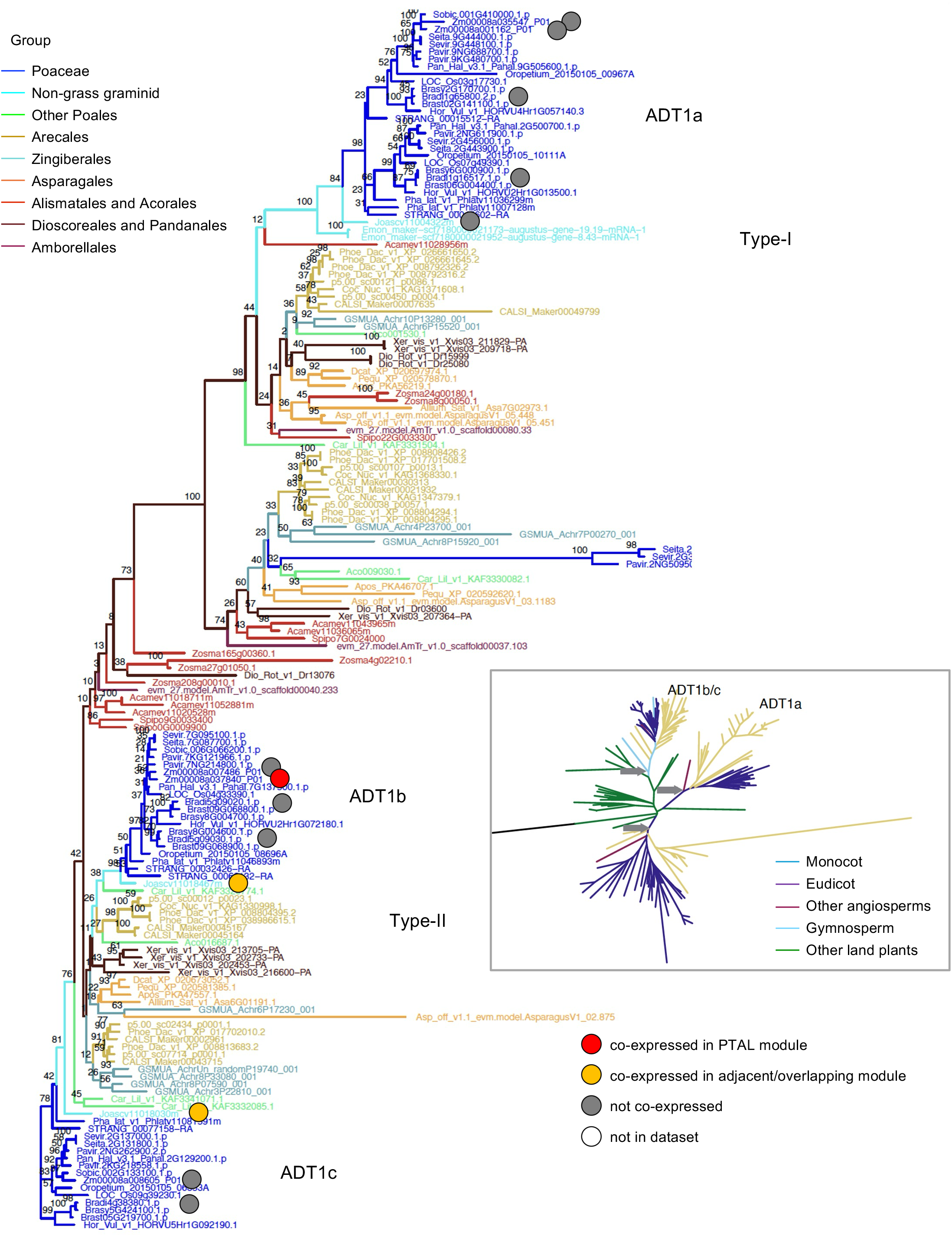
ADT1 phylogeny shows both conservation of co-expression between *Z. mays* and *J. ascendens* and co-expression loss in grasses. The main tree is built using monocot species while the inset tree is built using species from all plant clades. The grey arrows in the inset tree shows where the base of the monocot tree starts.

**Figure S24B.**
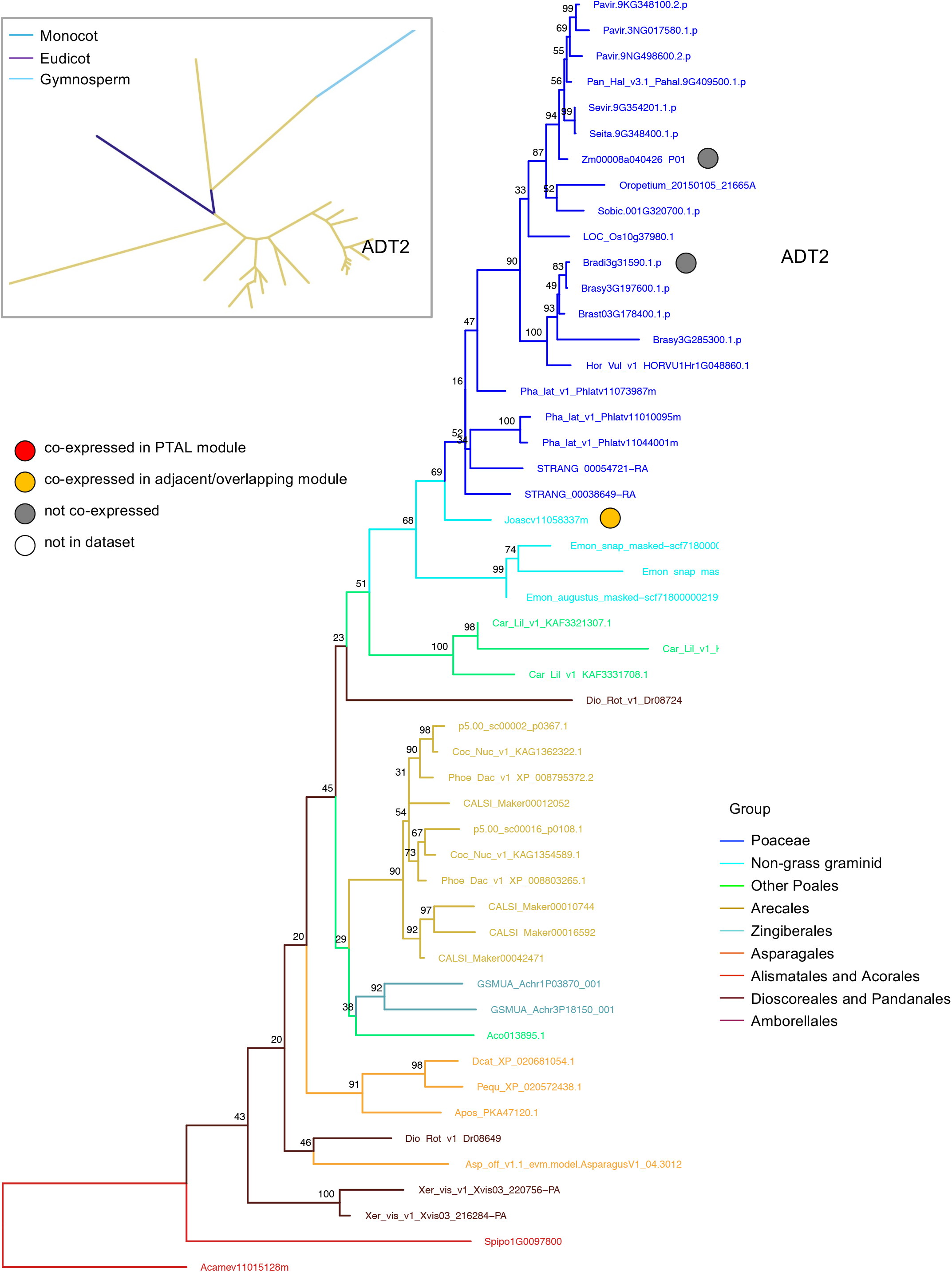
ADT2 phylogeny shows loss of co-expression in grasses. The main tree is built using monocot species while the inset tree is built using species from all plant clades.

**Figure S25.**
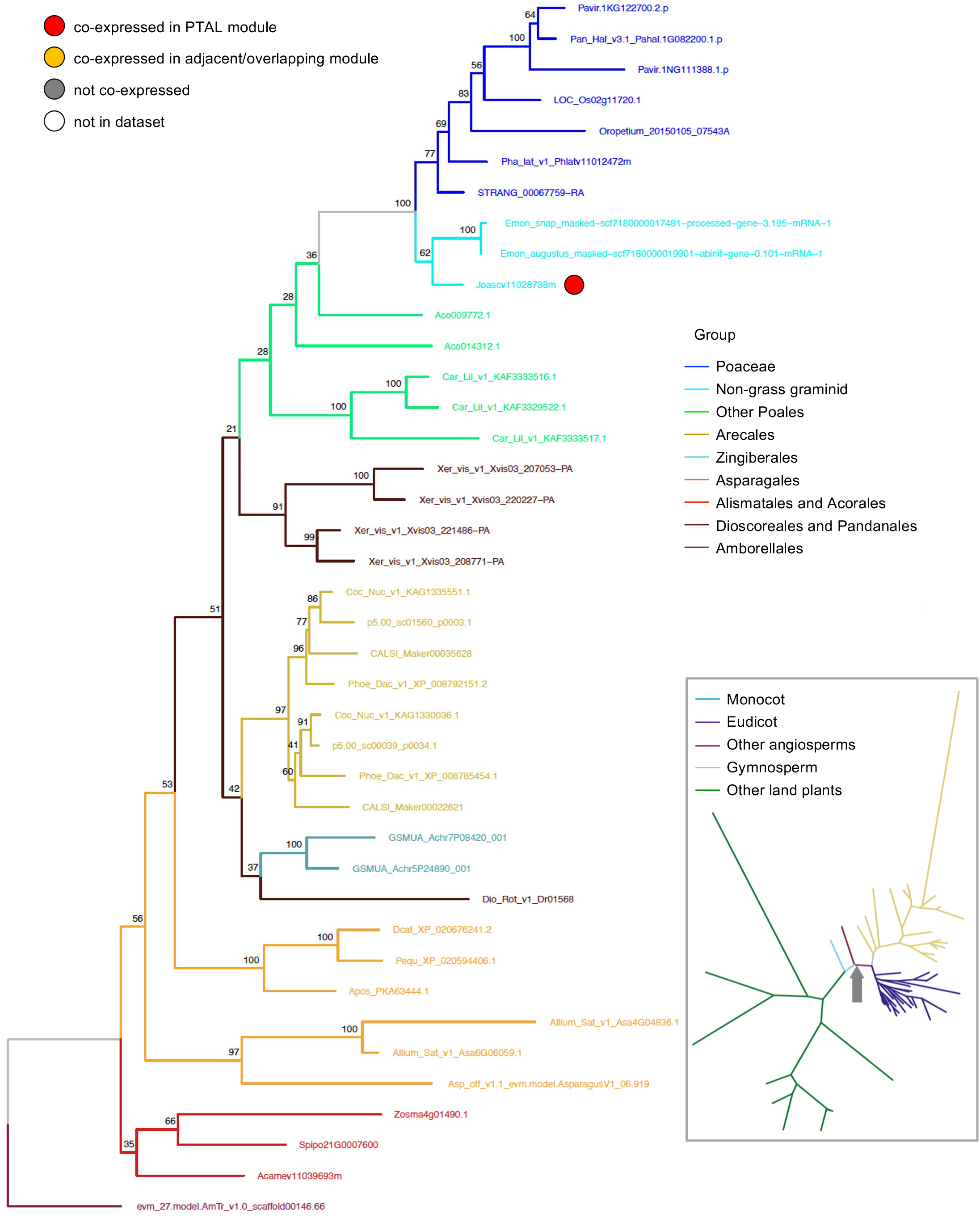
CSE phylogeny shows loss of genes in grasses. The main tree is built using monocot species while the inset tree is built using species from all plant clades. The grey arrow in the inset tree shows where the base of the monocot tree starts.

**Figure S26.**
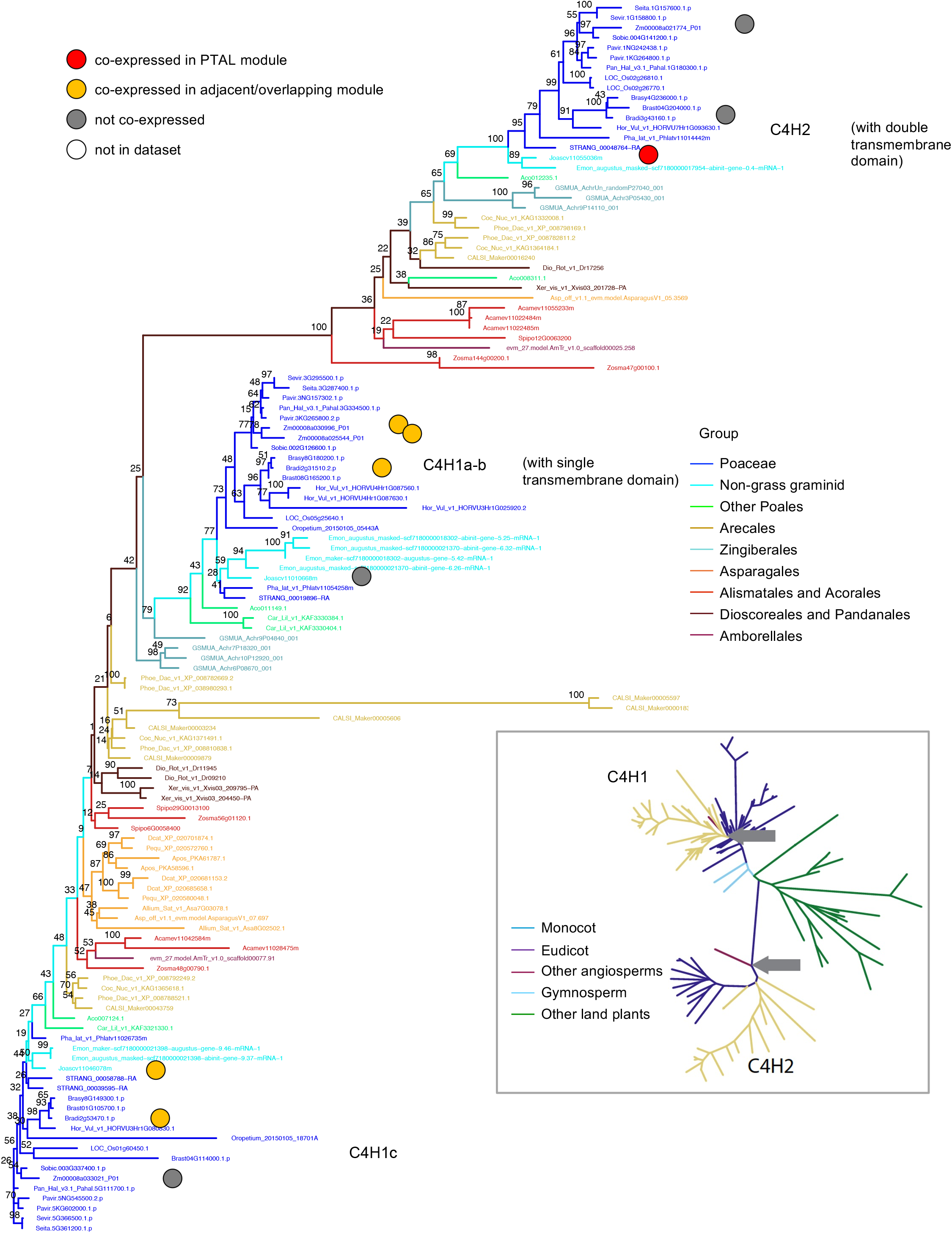
C4H phylogeny shows change of co-expression between *J. ascendens* and grass isoforms. The main tree is built using monocot species while the inset tree is built using species from all plant clades. The grey arrows in the inset tree show where the base of each C4H clade starts.

**Figure S27A.**
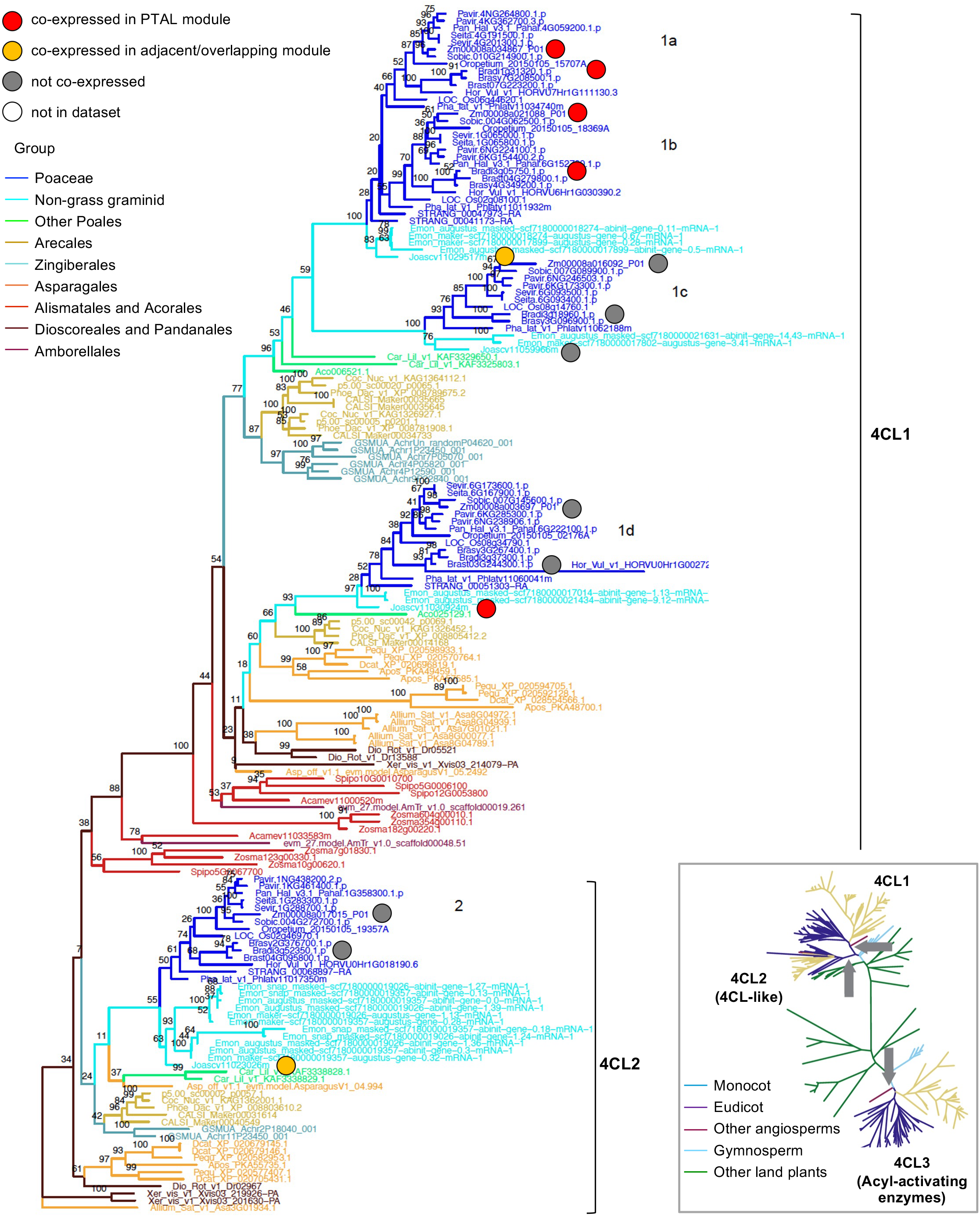
4CL1-2 phylogeny shows conservation of co-expression between grasses and *J. ascendens* as well as a loss of co-expression in grasses. Co-expression of isoform 4CL1a/b is maintained between grasses and *J. ascendens*, but isoforms 4CL1d and 4CL2 were co-expressed only in *J. ascendens*. The main tree is built using monocot species while the inset tree is built using species from all plant clades. The grey arrows in the inset tree show where the base of each 4CL clade starts.

**Figure S27B.**
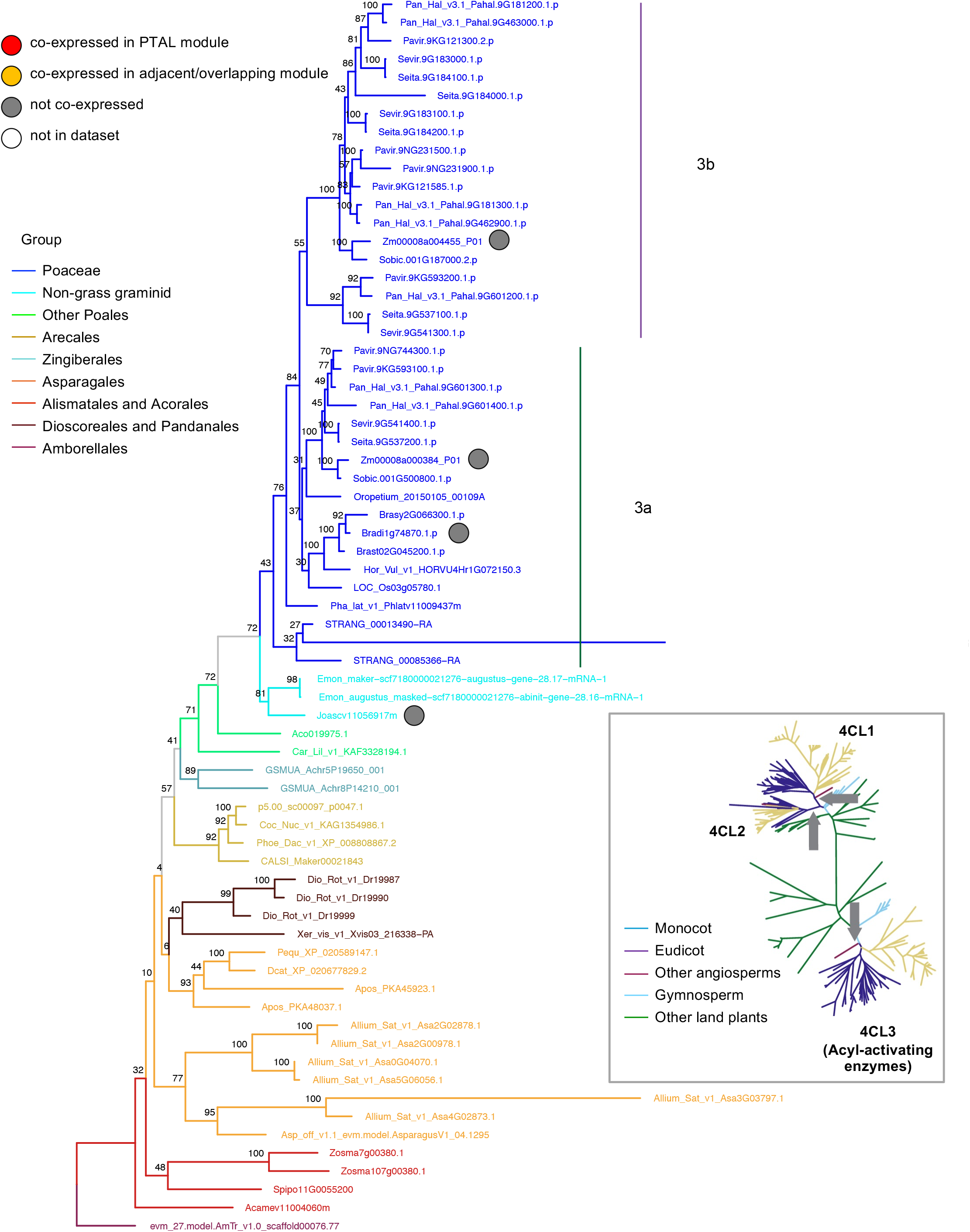
4CL3 phylogeny has no co-expression in *J. ascendens* or grasses. The main tree is built using monocot species while the inset tree is built using species from all plant clades. The grey arrows in the inset tree show where the base of each 4CL clade starts.

**Figure S28.**
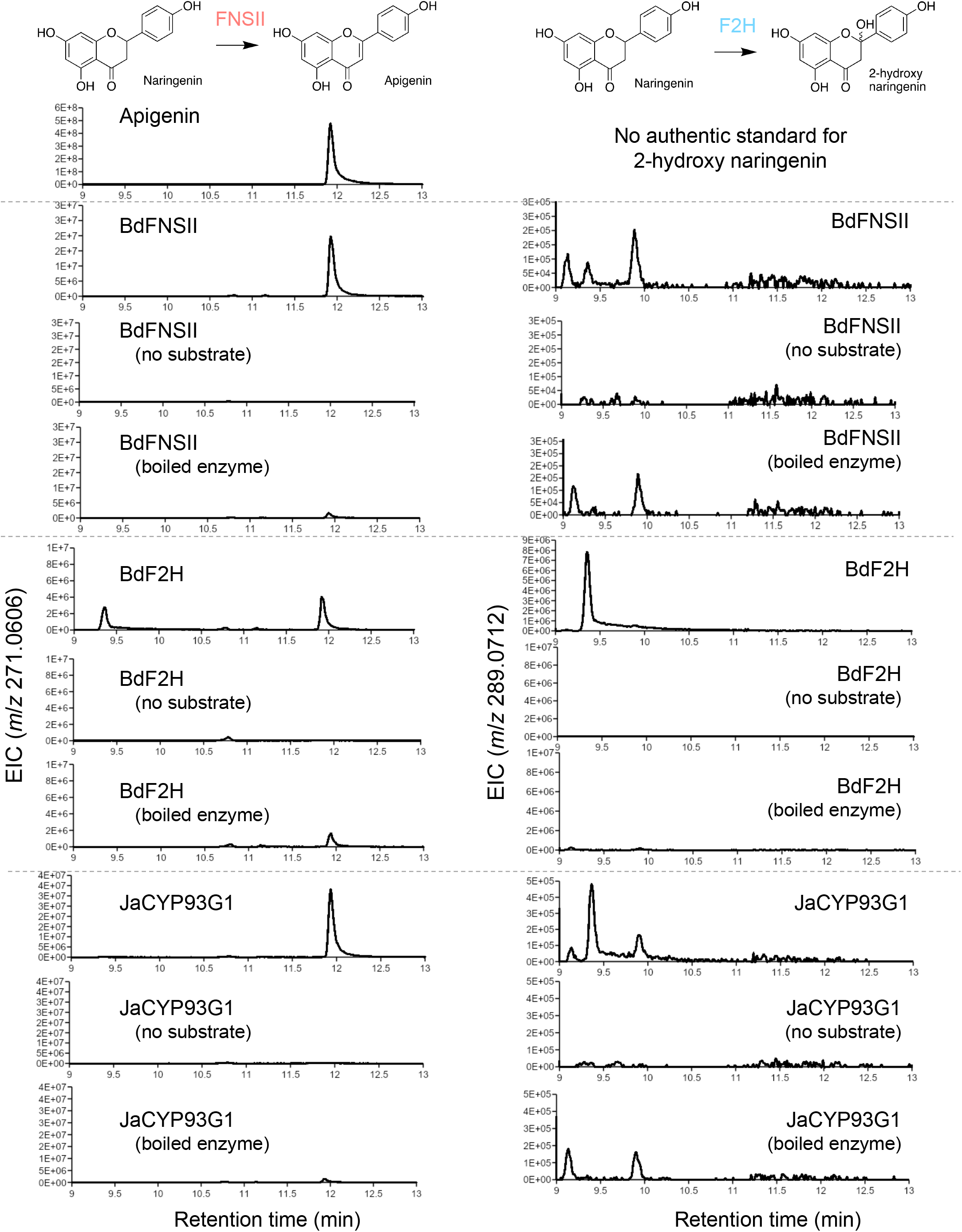
FNSII and F2H activities of FNSII and F2H from *B. distachyon* and their closest homolog in *J. ascendens* (JaCYP93G1), showing the JaCYP93G1 has FNSII activity.

**Figure S29.**
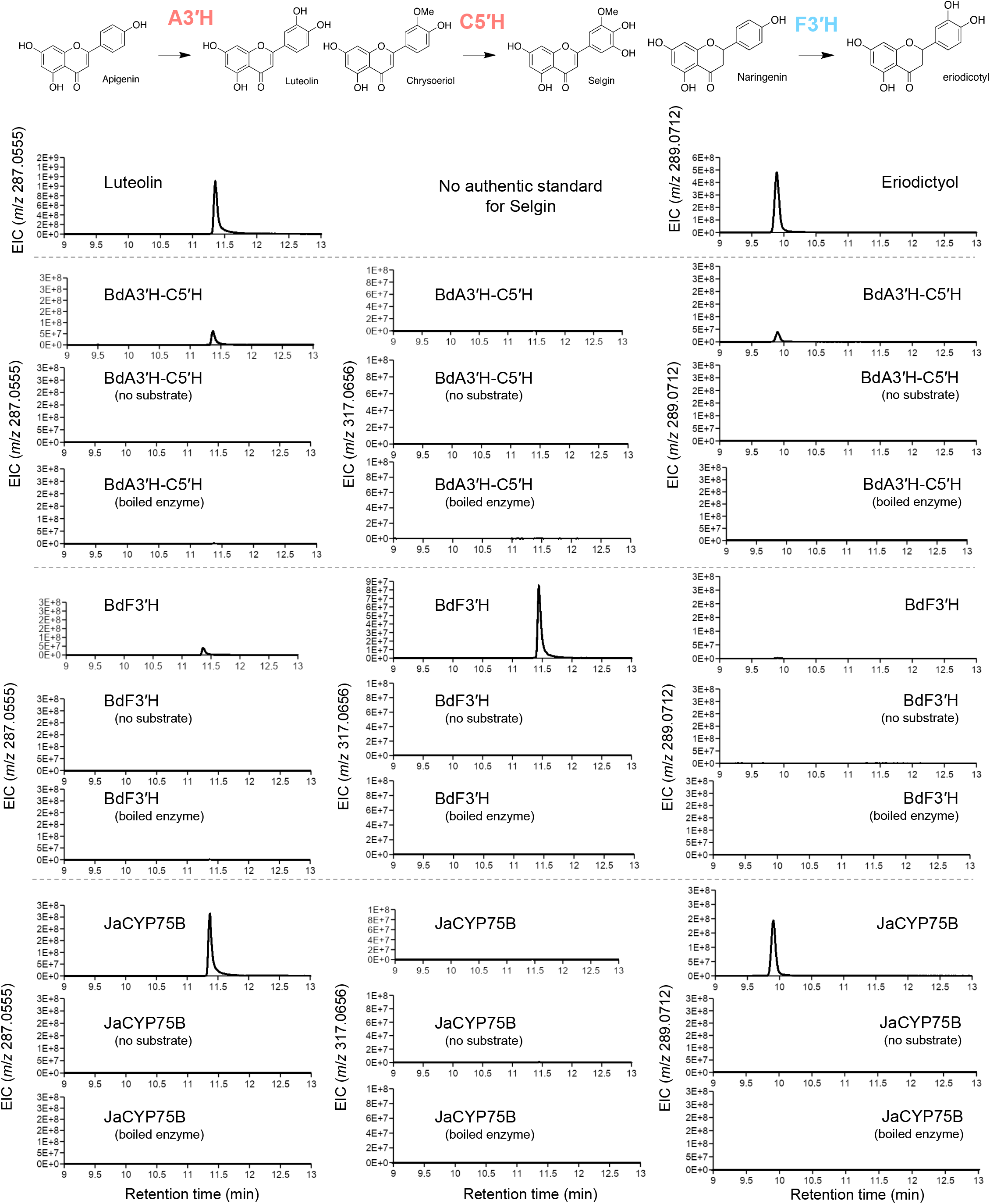
A3′H, C5′H, and F3′H activities of A3′H-C5′H and F3′H from *B. distachyon* and their closest homolog in *J. ascendens* (JaCYP75B), showing the *J. ascendens* CYP75B enzyme lack C5′H activity.

**Figure S30.**
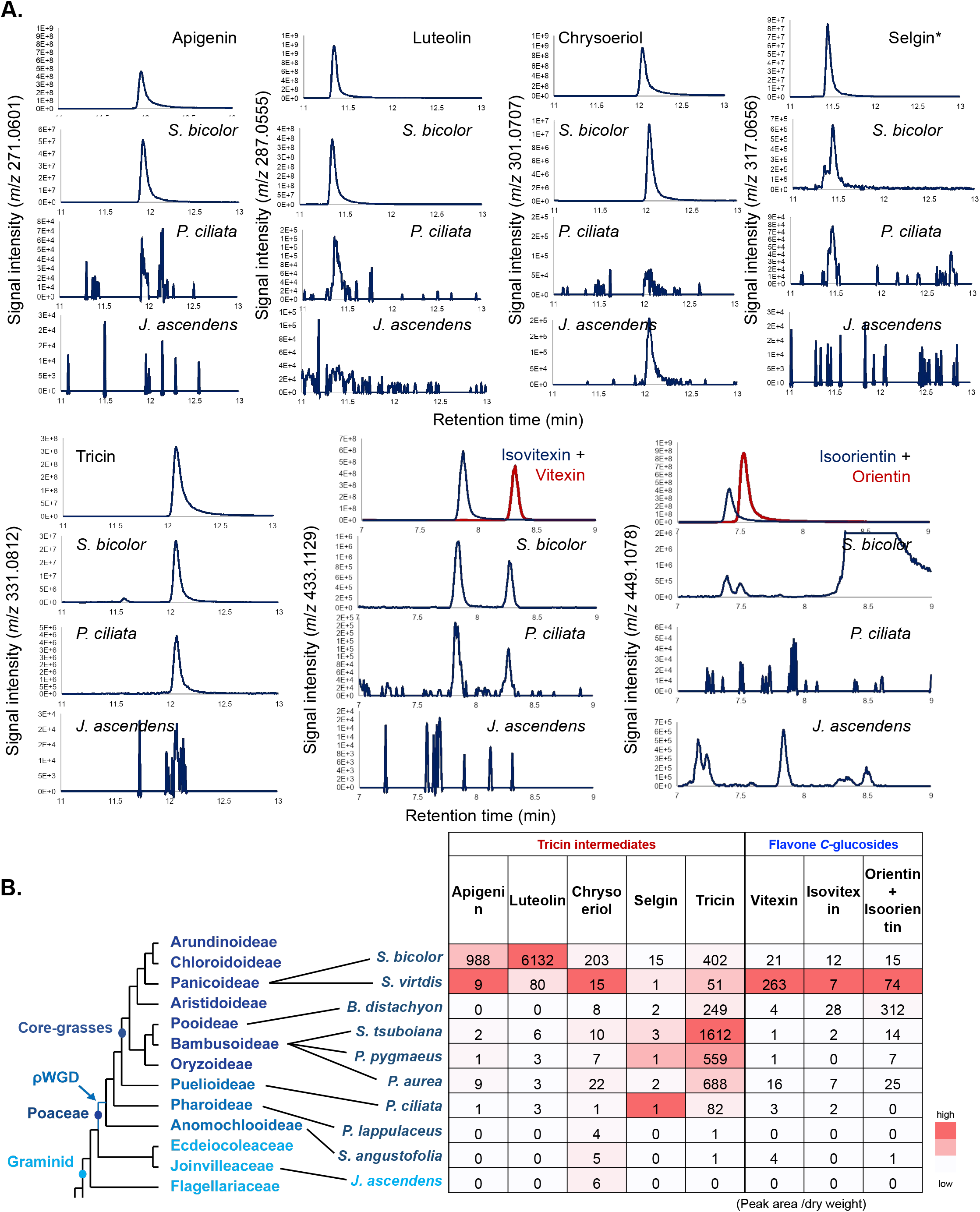
Accumulation of tricin intermediates and flavone *C*-glucosides determined by LC-MS analysis of the MeOH extract. Chromatogram of each compound for the selected species (A) and normalized LC-MS peak intensity (B) is shown. *Selgin standard peak was from C5′H enzyme reaction product.

**Figure S31.**
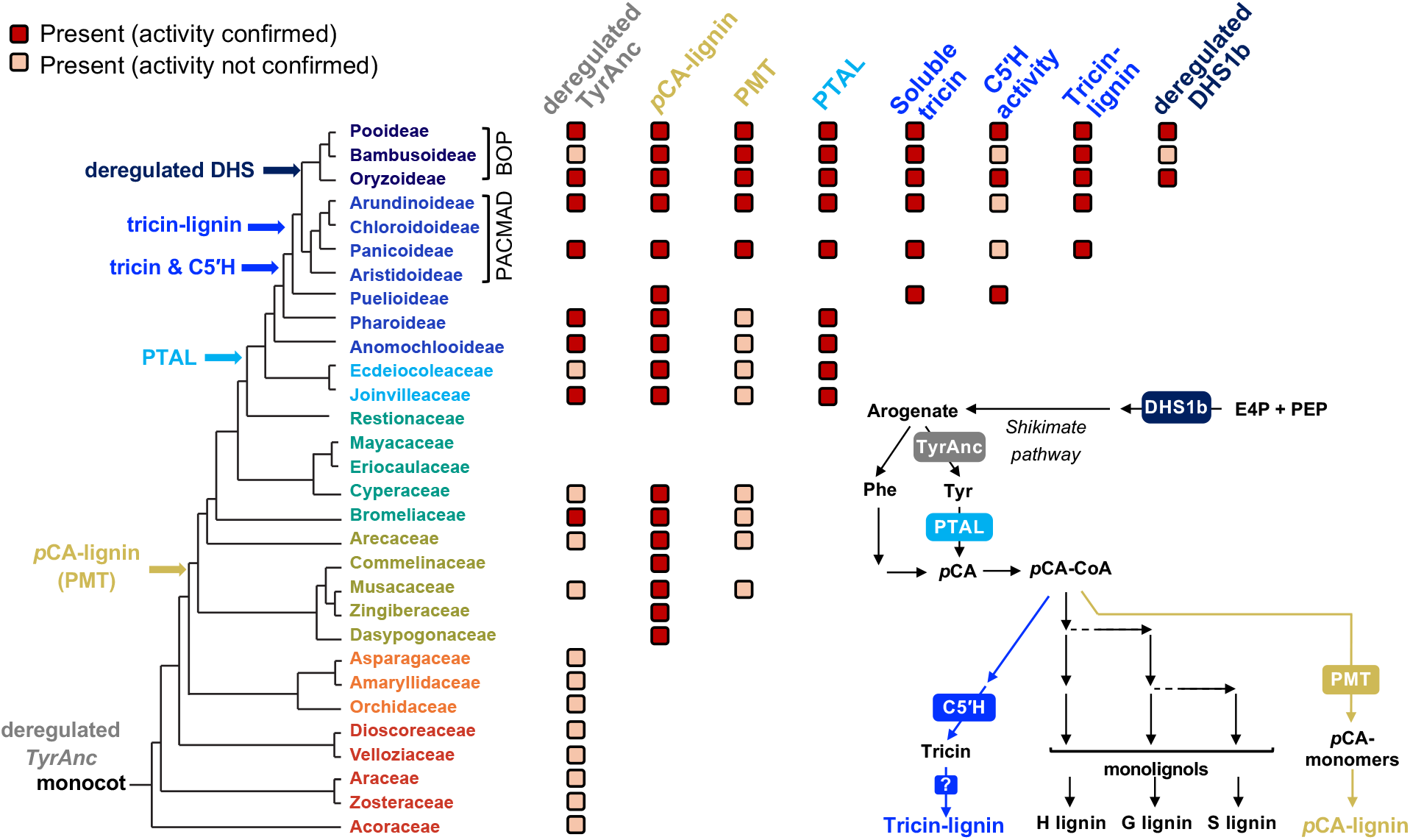
Grass lignin network evolved in a stepwise manner, in which *p*CA-lignin subunits occurred first, followed by tyrosine-derived lignin biosynthesis and then addition of tricin into the cell wall lignin. Monocot and Poales ancestors already had dregulated TyrAnc and PMT, which produce tyrosine and *p*CA- lignin, respectively. At the divergence of grasses and *J. ascendens*, PTAL emerged and generated two entry pathways for lignin biosynthesis. After the divergence of *Pharus* and *Puelia* C5’H activity emerged within the grass CYP75B family, leading to the acquisition of the tricin biosynthetic pathway, with soluble tricin later being incorporated into the cell wall lignin within core-grasses. The evolution of PTAL likely elevated the demand of tyrosine, which further facilitated the de-regulation of TyrAnc and the emergence of deregulated DHS1b isoform, which increased the upstream shikimate pathway activity in BOP grasses. Dark red boxes denote the presence of confirmed enzyme activity or chemical accumulation, while light orange boxes indicate the presence of orthologous genes whose activity have not ben confirmed.

**Figure S32.**
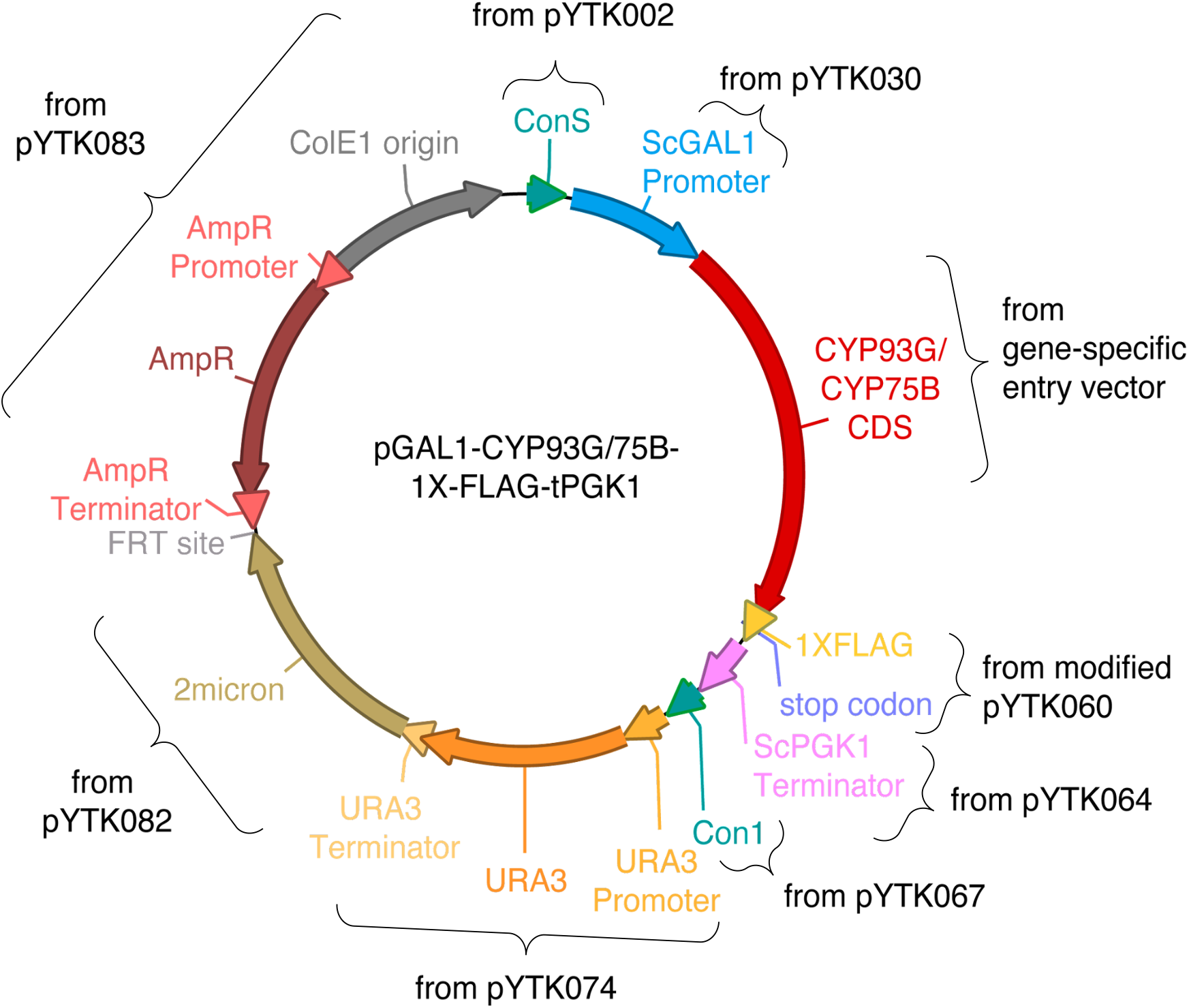
Golden gate constructs for recombinant protein induction in *S. cerevisiae* WAT11.

