## Supplemental Figures for "Establishment of the grass lignin metabolic network during monocot evolution"

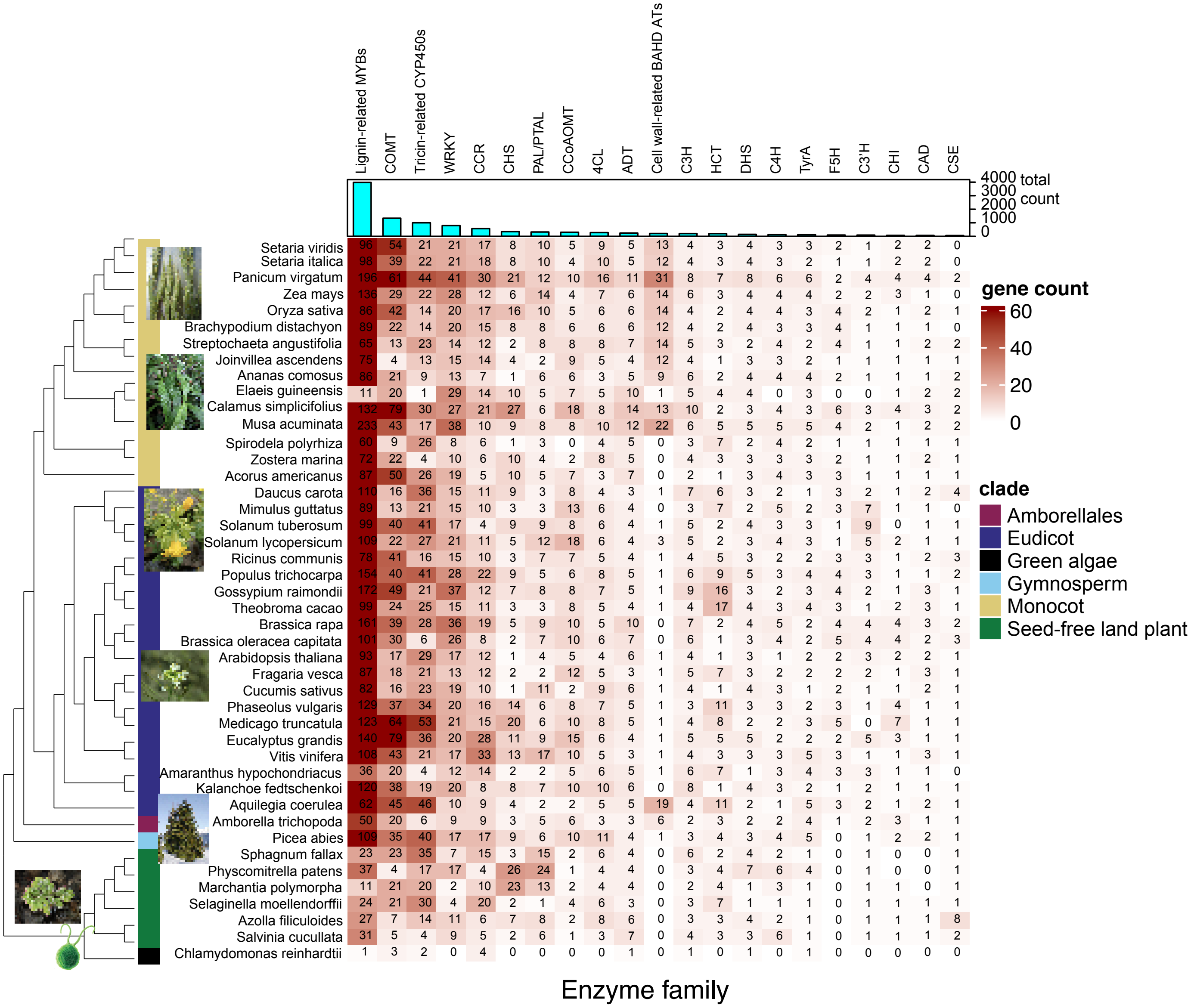


**Figure S1. Enzyme and TF families related with phenylpropanoid biosynthesis evolve at different times over the course of lignin evolution.**

The heatmap represents the number of gene isoforms of an enzyme or transcription factor family belonging to the lignin pathway (column) in a given species (row). The darker the red color, the more isoforms are present. The bar graph at the top indicates the total number of isoforms for an enzyme family in all species. The color bars on the phylogeny denote different taxonomic clades of plants and algae as indicated.

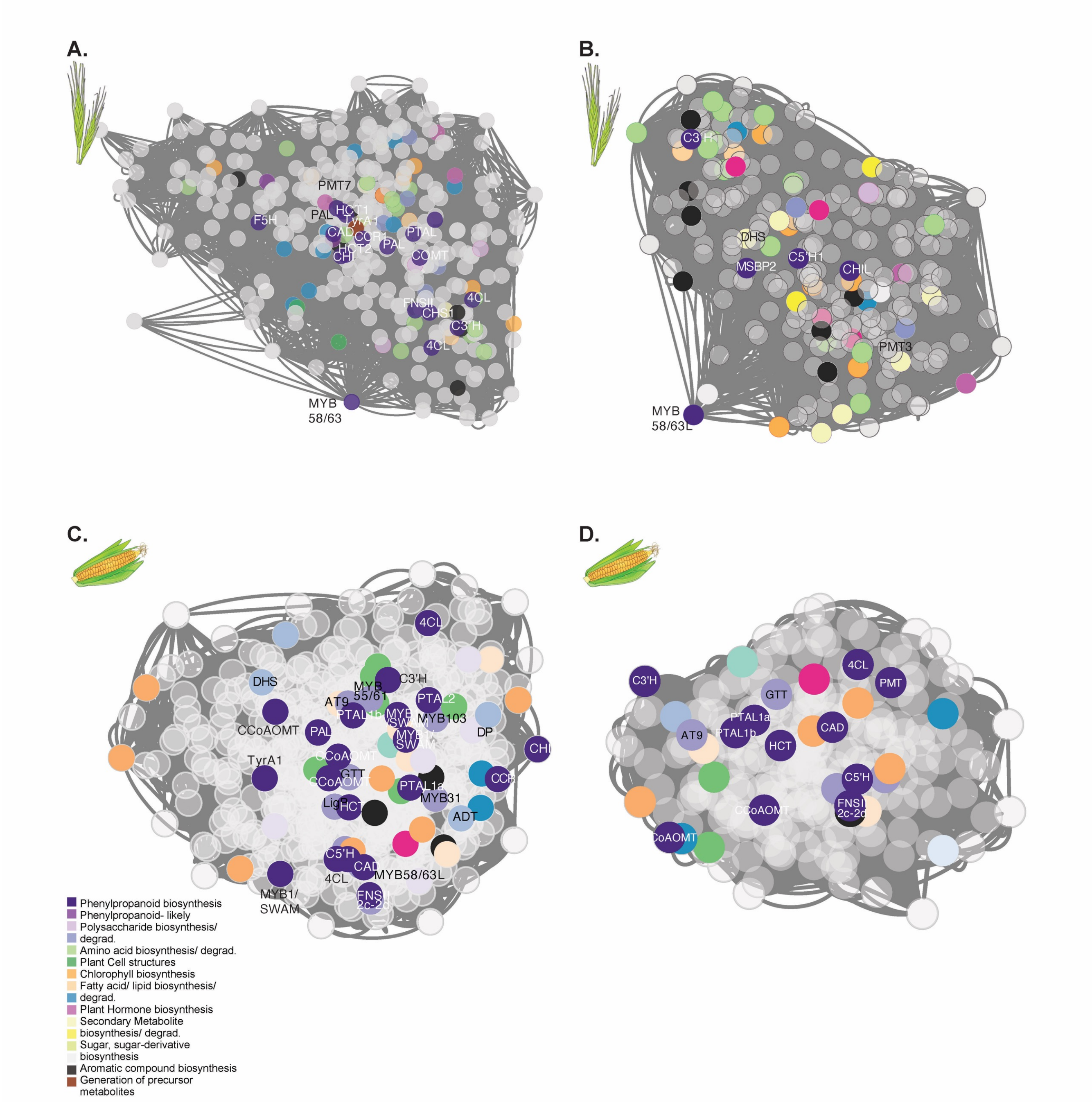


**Figure S2. Full co-expression modules from PTAL/TyrA1 and A3'H-C5'H representing the lignin networks in *Brachypodium distachyon* and *Zea mays.***

**A.** Co-expression of the lignin network in *B. distachyon* based on PTAL and TyrA1 genes, where each node is a gene and connections are representative of the distances between genes based on Spearman's mutual rank. Known lignin enzymes are in dark purple and labeled with their enzyme abbreviation. Other colors correspond to pathways in different categories. Genes in grey belong to an unknown pathway. Module size is 460 genes.

**B.** Co-expression of the lignin network in *B. distachyon* based on A3'H-C5'H. Nodes, distances, and colors are the same as above.

**C.** Co-expression of the lignin network in *Z. mays* based on PTAL and TyrA1 genes. Each node is a gene and connections are representative of the distances between genes based on Spearman's mutual rank. Known lignin enzymes are in dark purple and labeled with their enzyme abbreviation. Other colors correspond to pathways in different categories. Genes in grey belong to an unknown pathway. Module size is 600 genes.

**D.** Co-expression of the lignin network in *Z. mays* based on A3'H-C5'H. Nodes, distances, and colors are the same as above.

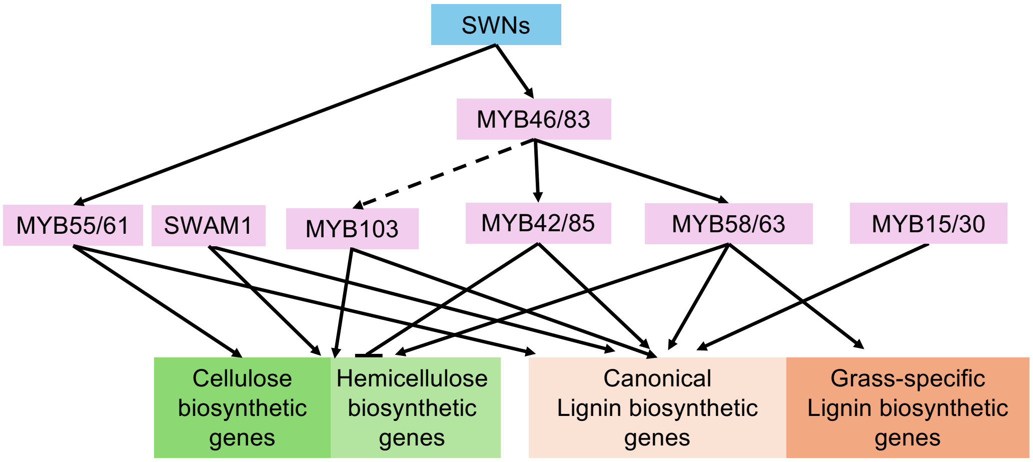


**Figure S3. Positive regulatory network of MYB transcription factors in grass lignin formation**
Positive transcriptional regulation is represented by arrows, and negative transcriptional regulation is shown by bars at the ends of lines. The scheme is based on Rao and Dixon (2018) and Miyamoto et al. (2020), with some modifications. SWN, secondary wall-associated NAC.

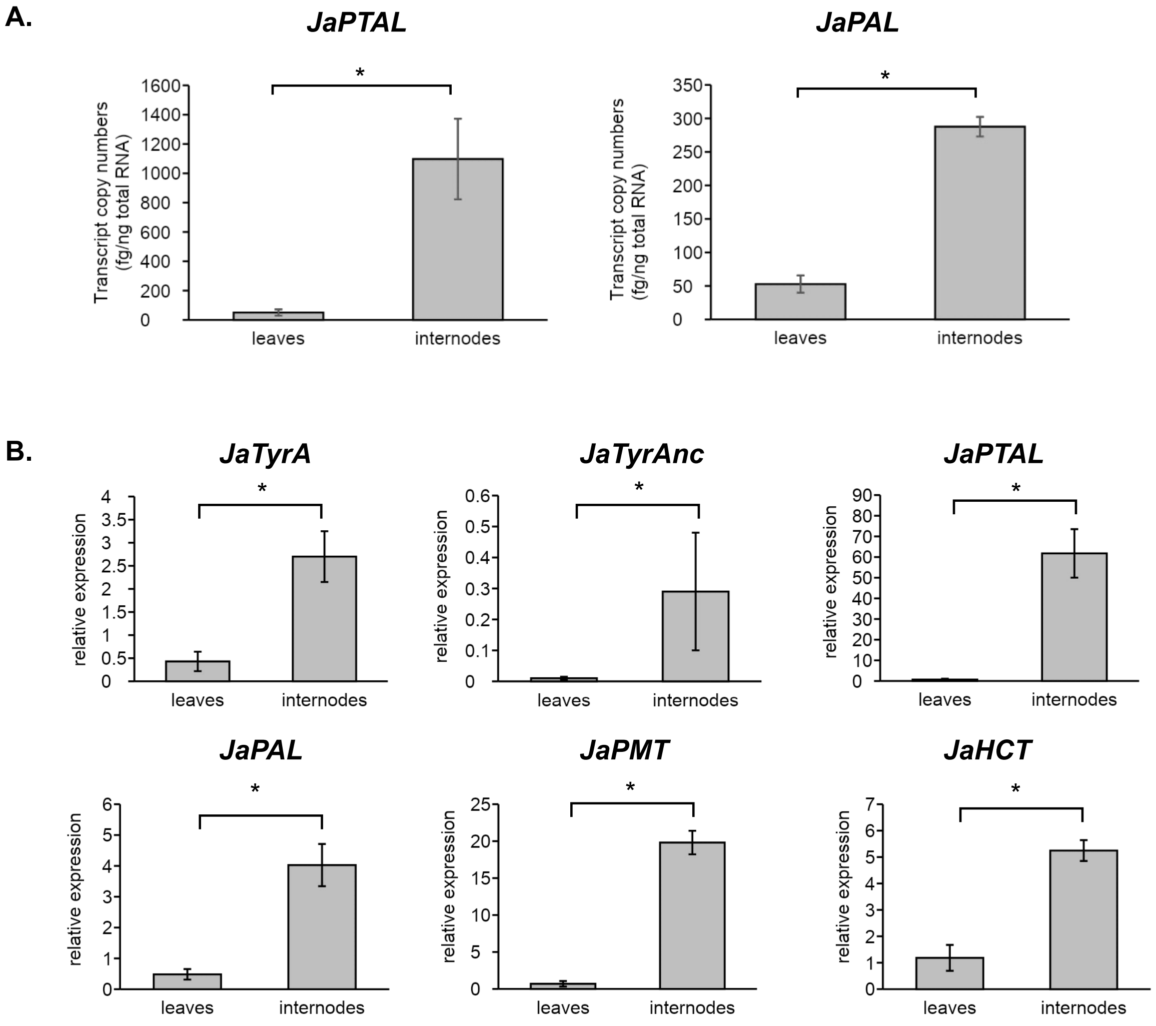


**Figure S4 Gene expression analysis by qRT-PCR demonstrates the existence of tyrosine-derived lignin pathway in *J. ascendense*.**
**A.** Transcript copy numbers of *JaPTAL* and *JaPAL* in 8 month old leaves and internodes.
**B.** RT-qPCR quantification, relative to the expression of *JaUBI10*, revealed that transcripts of JaTyrAs and other lignin pathway genes were much more abundant in the internodes compared to the leaves.
Values are mean ± SD ($n$ = 3). Asterisks indicate significant difference (Student's $t$-test; $p < 0.05$)

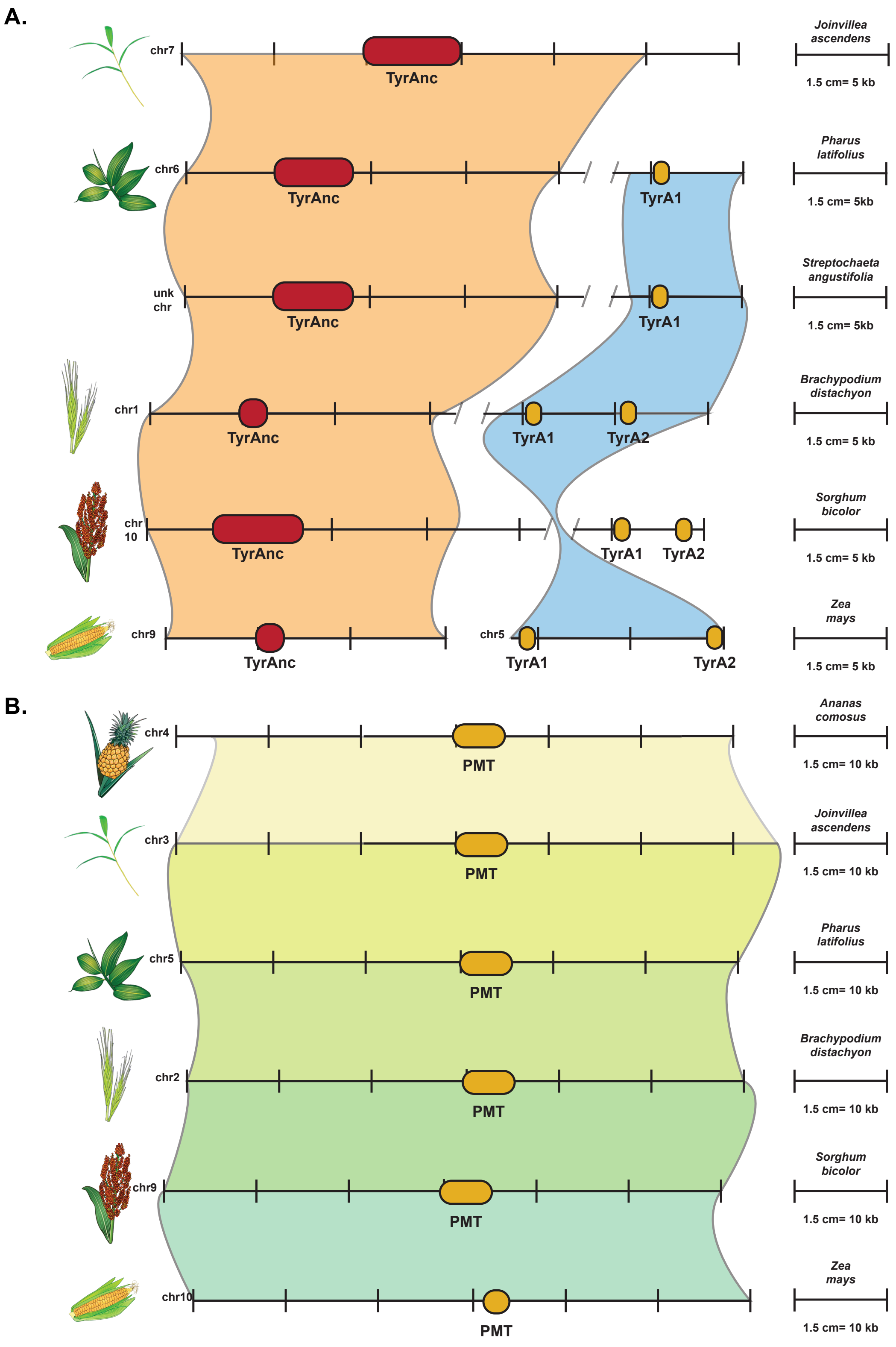


**Figure S5. PMT and TyrAnc are syntenic across Poales species, while TyrA1 becomes syntenic within grasses.**
**A.** Synteny of TyrA across Poales species showing differences between the isoforms TyrA1 and TyrAnc. TyrAnc is shown in red, while TyrA1/2 is shown in yellow. Relative distance on each chromosome is given by the scale for each species.
**B.** Synteny of PMT across Poales species. Relative distance on each chromosome is given by the scale for each species.

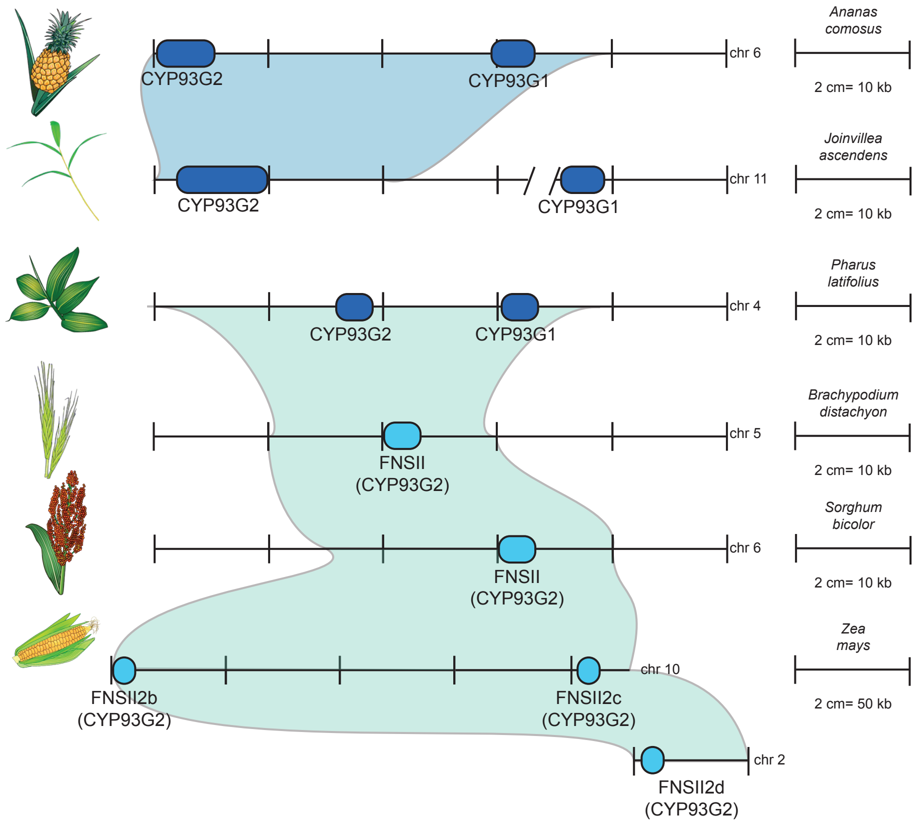


**Figure S6. FNSII is syntenic across grasses, as well as between pineapple and Joinvillea, but synteny breaks down between grasses and non-grass Poales.**

Synteny of CYP93G across Poales species are shown. Dark blue genes are CYP93Gs with either unknown activity or dual FNSII/F2H activity. Light blue genes are CYP93Gs with only FNSII activity. Relative distance on each chromosome is given by the scale for each species.

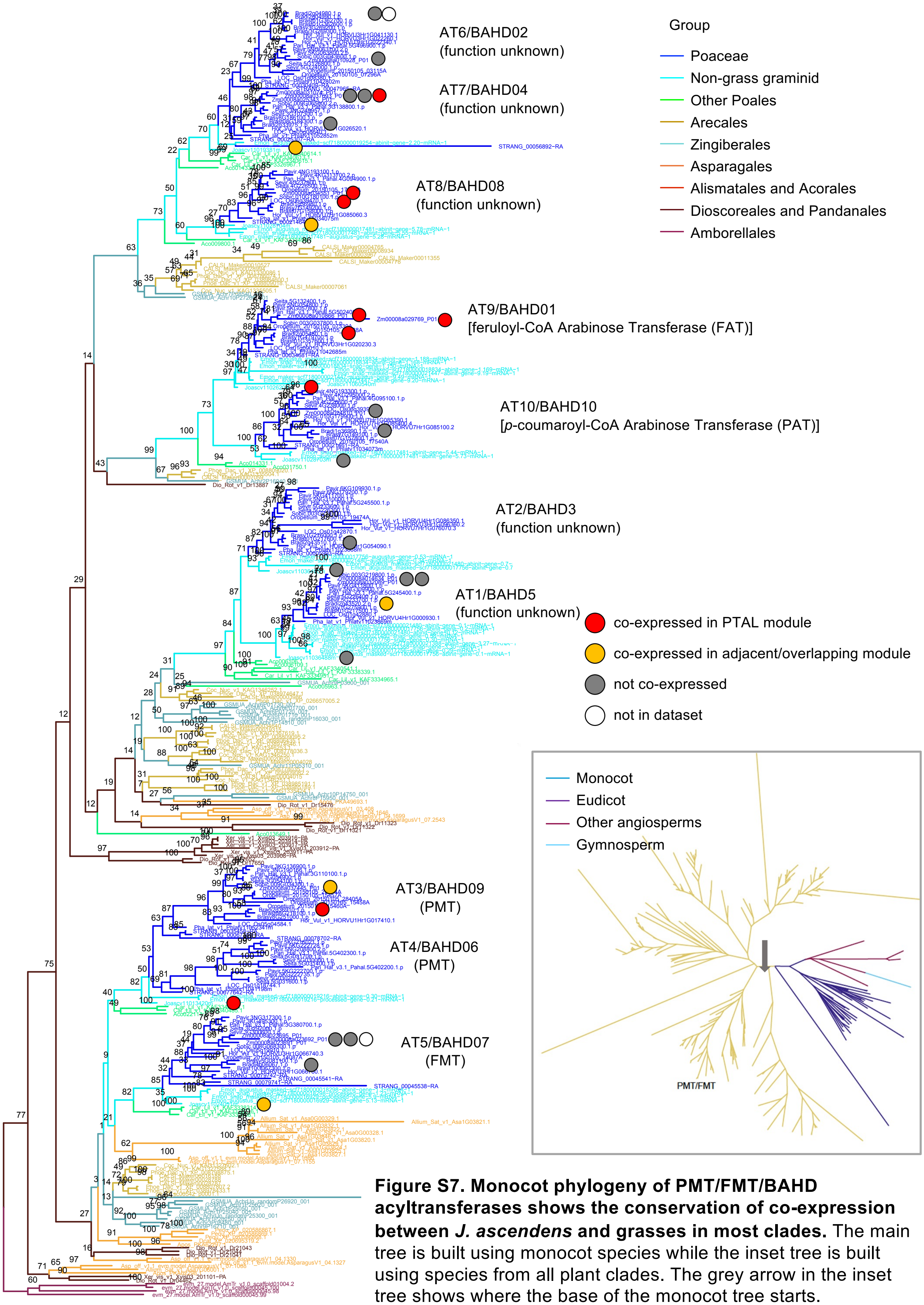


**Figure S7. Monocot phylogeny of PMT/FMT/BAHD acyltransferases shows the conservation of co-expression between *J. ascendens* and grasses in most clades.** The main tree is built using monocot species while the inset tree is built using species from all plant clades. The grey arrow in the inset tree shows where the base of the monocot tree starts.

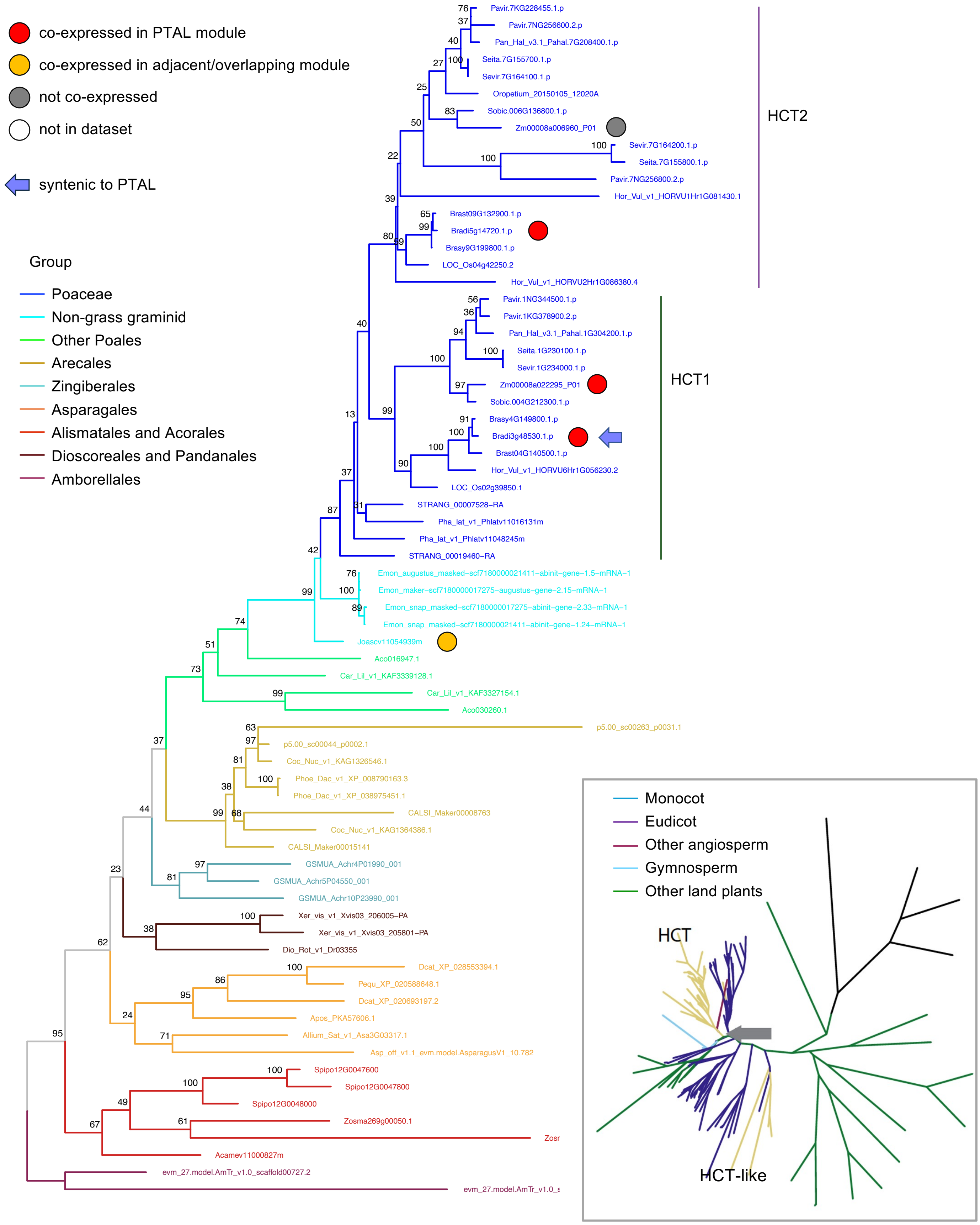


**Figure S8. HCT phylogeny shows the conservation of co-expression between *J. ascendens* and grasses.** The main tree is built using monocot species while the inset tree is built using species from all plant clades. The grey arrow in the inset tree shows where the base of the monocot tree starts.

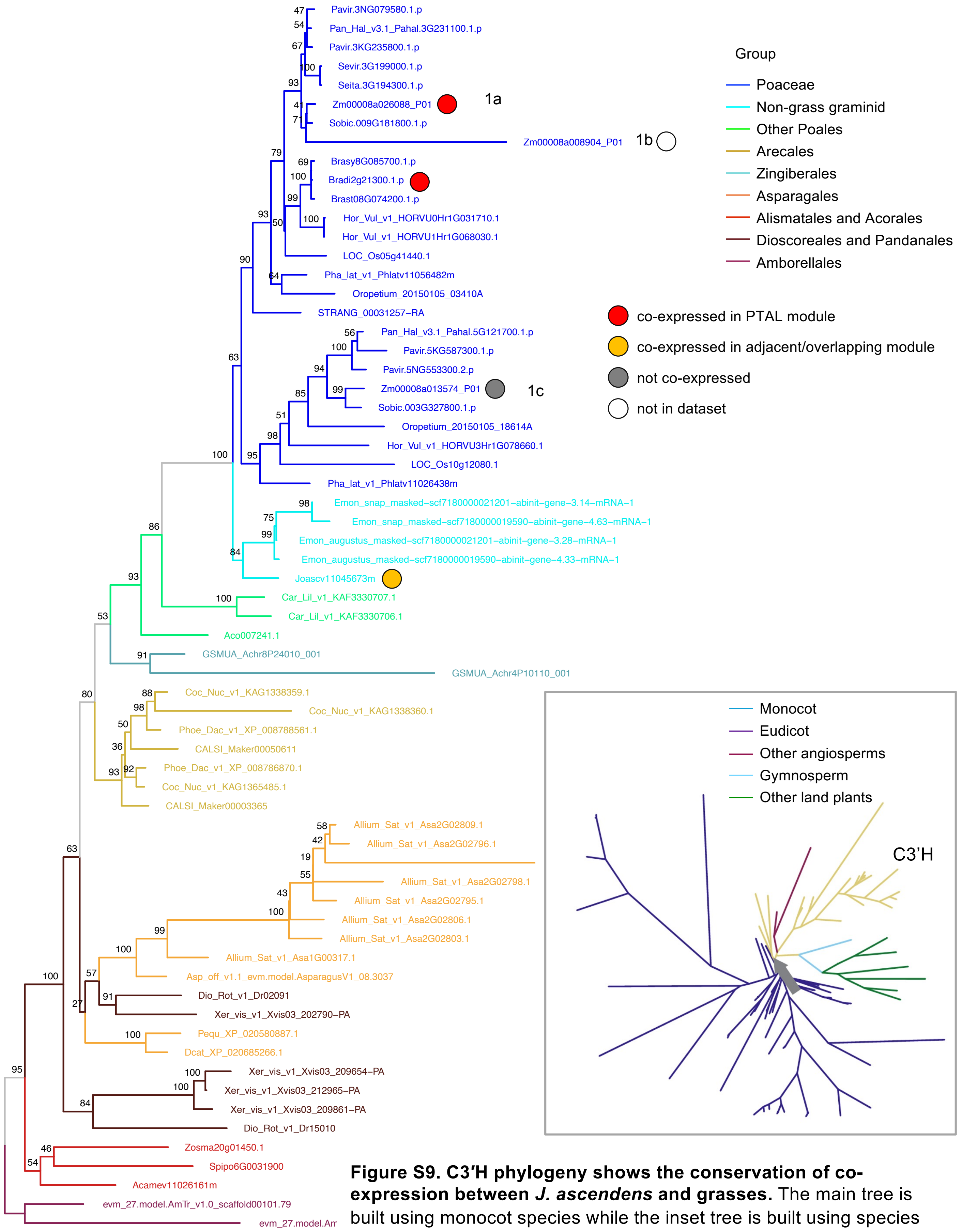


**Figure S9. C3′H phylogeny shows the conservation of co-expression between *J. ascendens* and grasses.** The main tree is built using monocot species while the inset tree is built using species from all plant clades. The grey arrow in the inset tree shows where the base of the monocot tree starts.

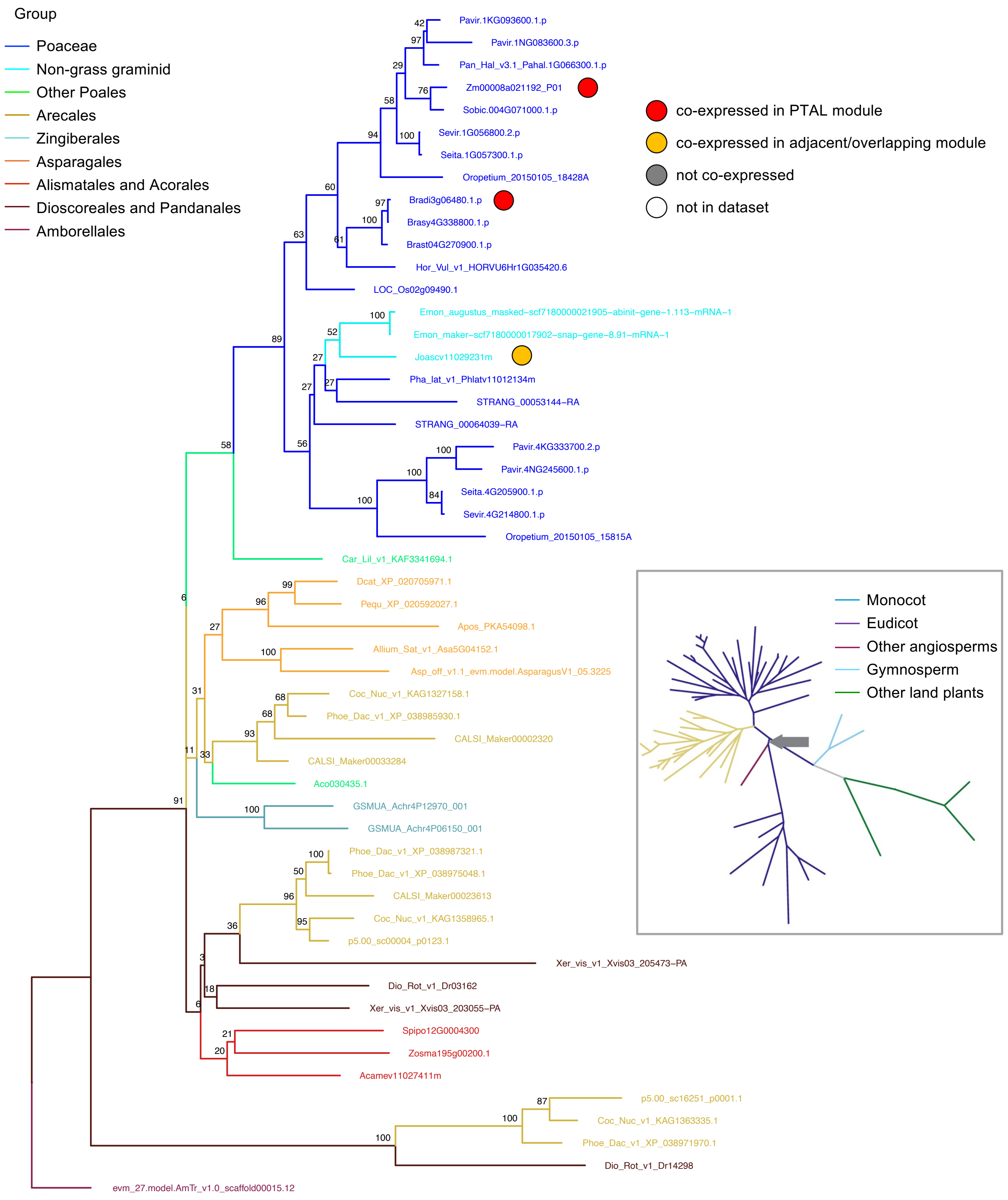


**Figure S10. CAD phylogeny shows the conservation of co-expression between *J. ascendens* and grasses.** The main tree is built using monocot species while the inset tree is built using species from all plant clades. The grey arrow in the inset tree shows where the base of the monocot tree starts.

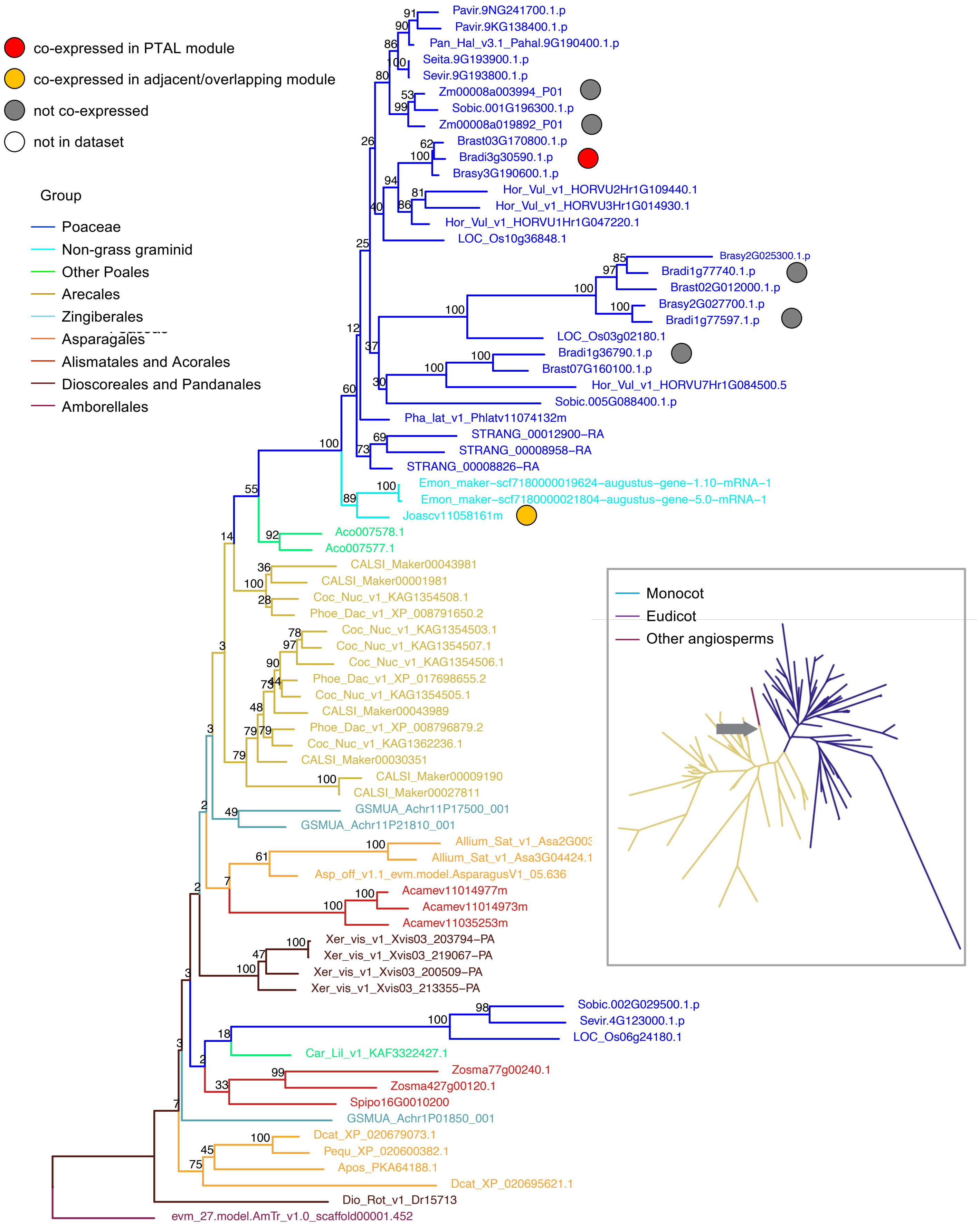


**Figure S11. F5H phylogeny shows conservation of co-expression between *B. distachyon* and *J. ascendens*, but loss in *Z. mays*.** The main tree is built using monocot species while the inset tree is built using species from all plant clades. The grey arrow in the inset tree shows where the base of the monocot tree starts.

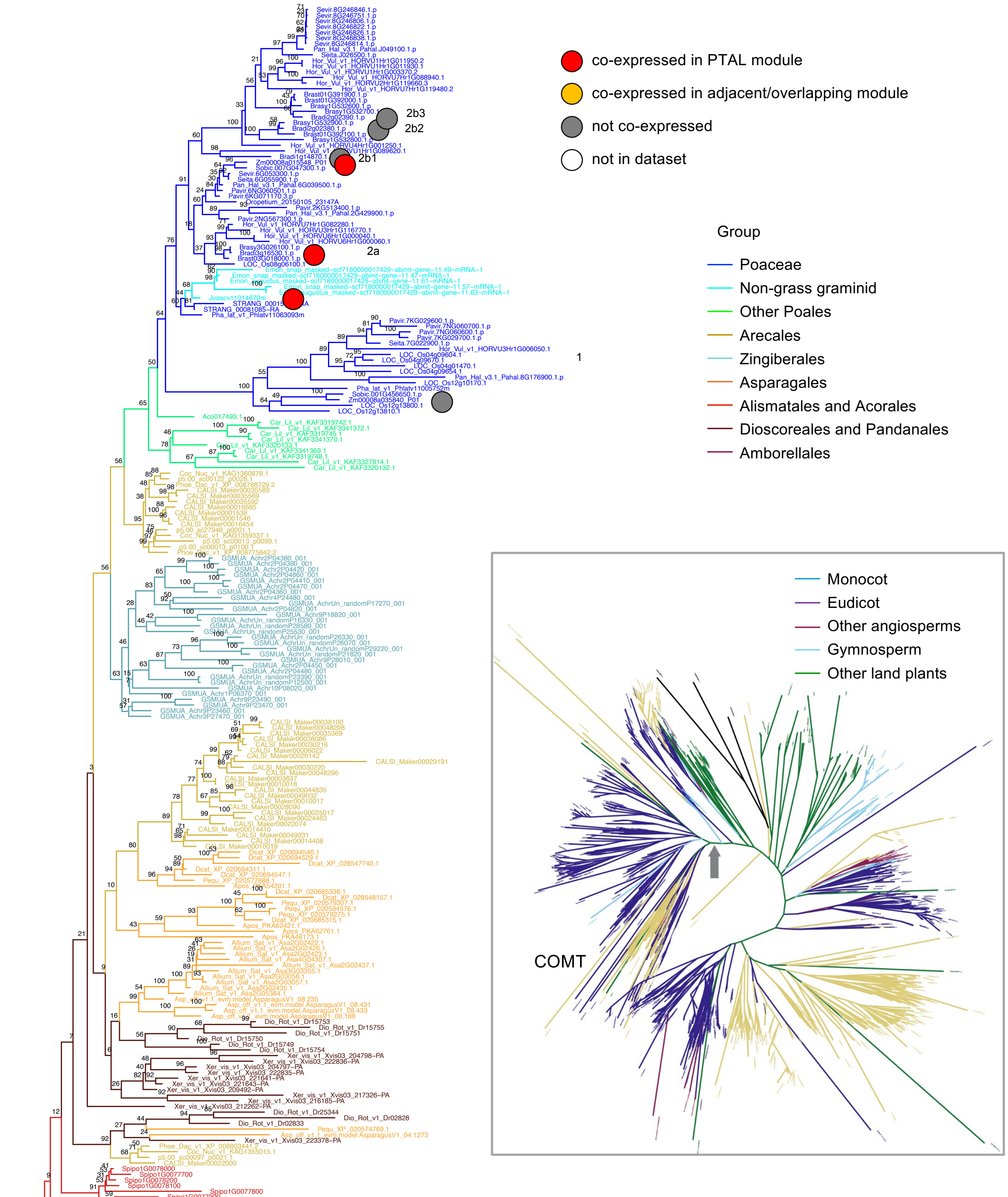

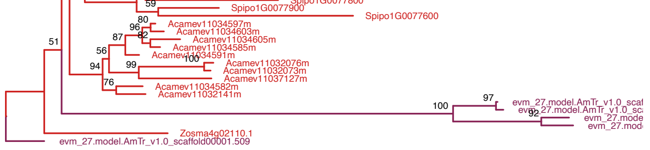

**Figure S12. COMT phylogeny shows the conservation of co-expression between *J. ascendens* and grasses.** The main tree is built using monocot species while the inset tree is built using species from all plant clades. The grey arrow in the inset tree shows where the base of the monocot tree starts.

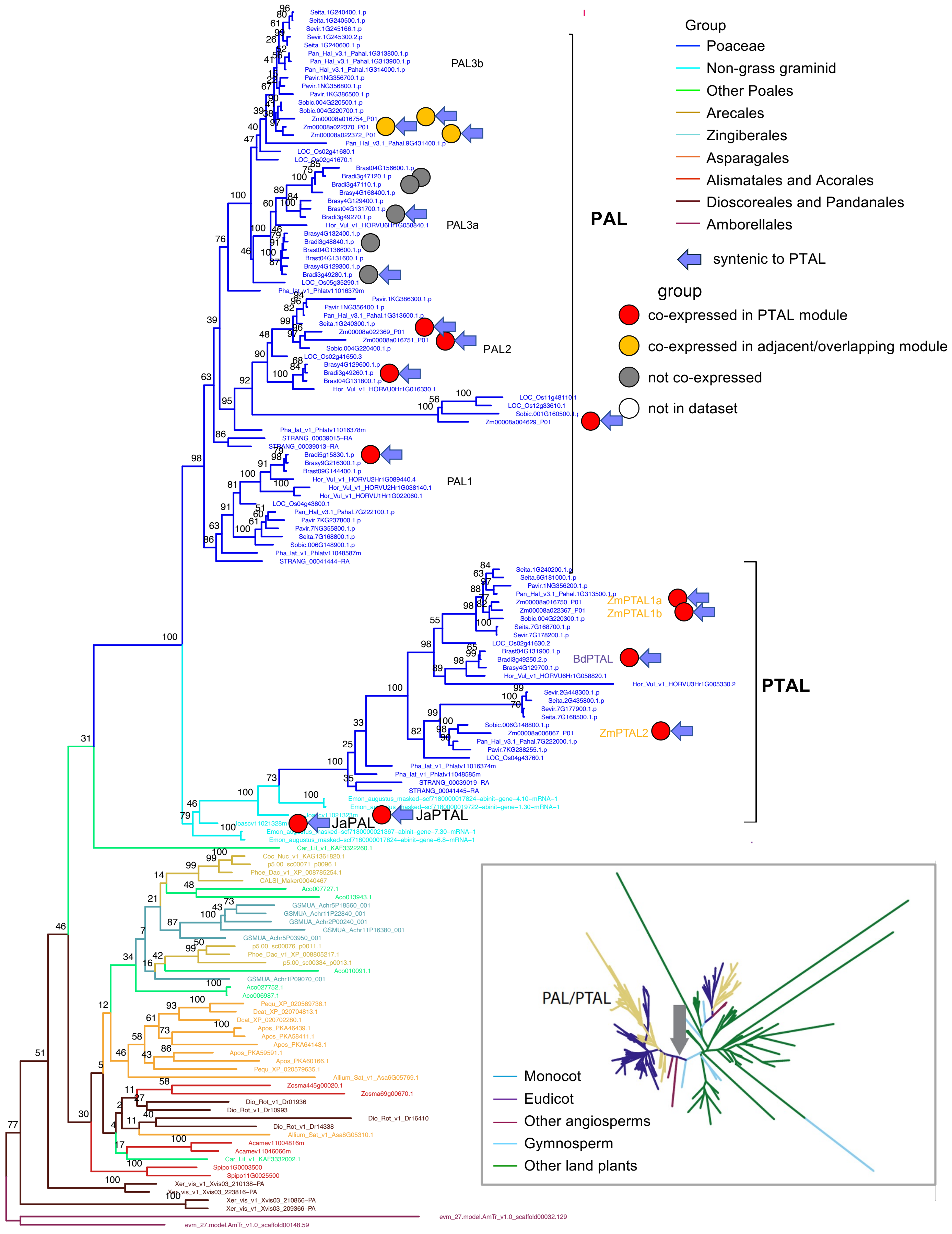


**Figure S13. PAL phylogeny shows maintenance of co-expression between *J. ascendens* and grasses, but expansion of isoforms in grasses.** The main tree is built using monocot species while the inset tree is built using species from all plant clades. The grey arrow in the inset tree shows where the base of the monocot tree starts.

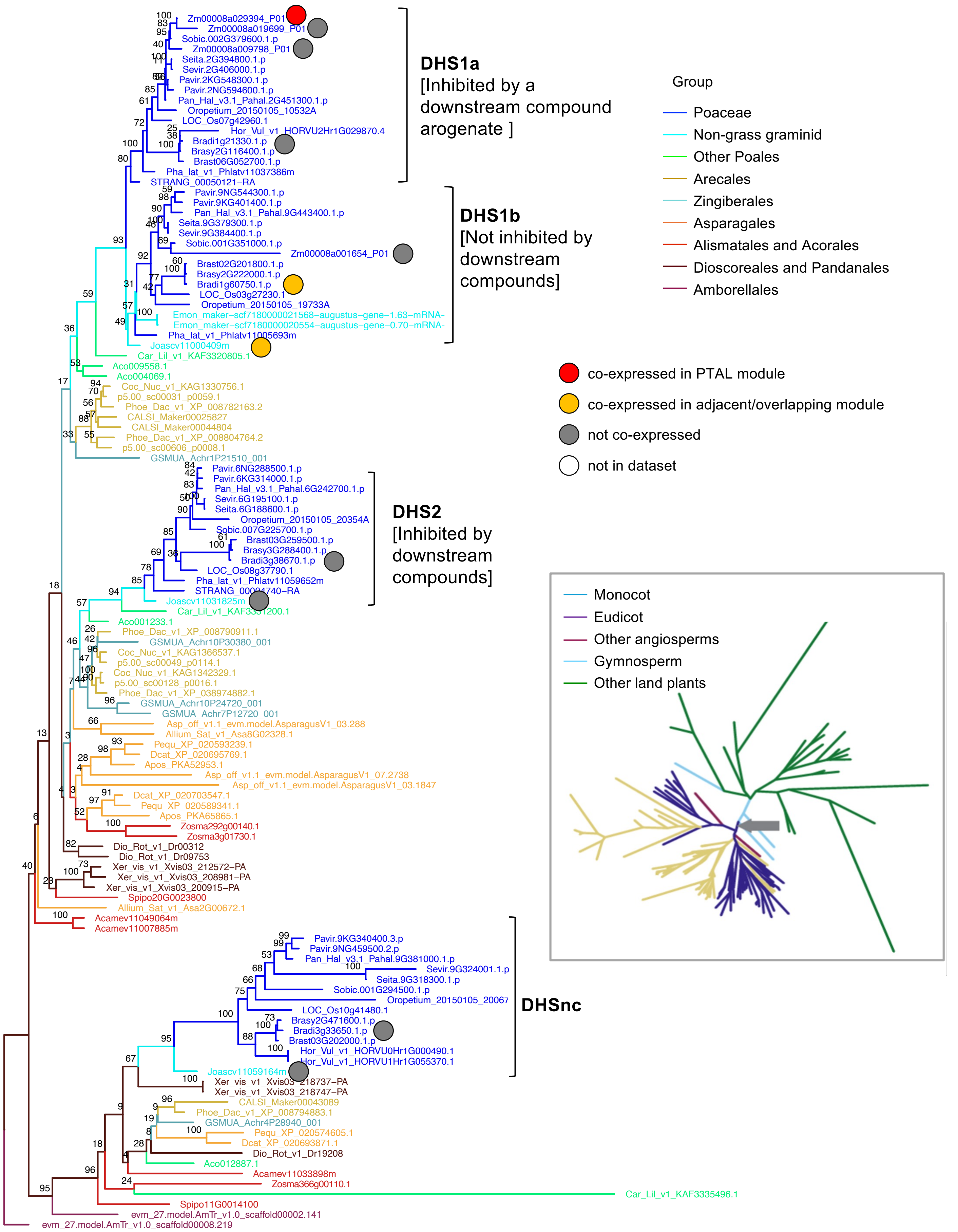


**Figure S14. DHS phylogeny shows the conservation of co-expression between *J. ascendens* and grasses.** The main tree is built using monocot species while the inset tree is built using species from all plant clades. The grey arrow in the inset tree shows where the base of the monocot tree starts.

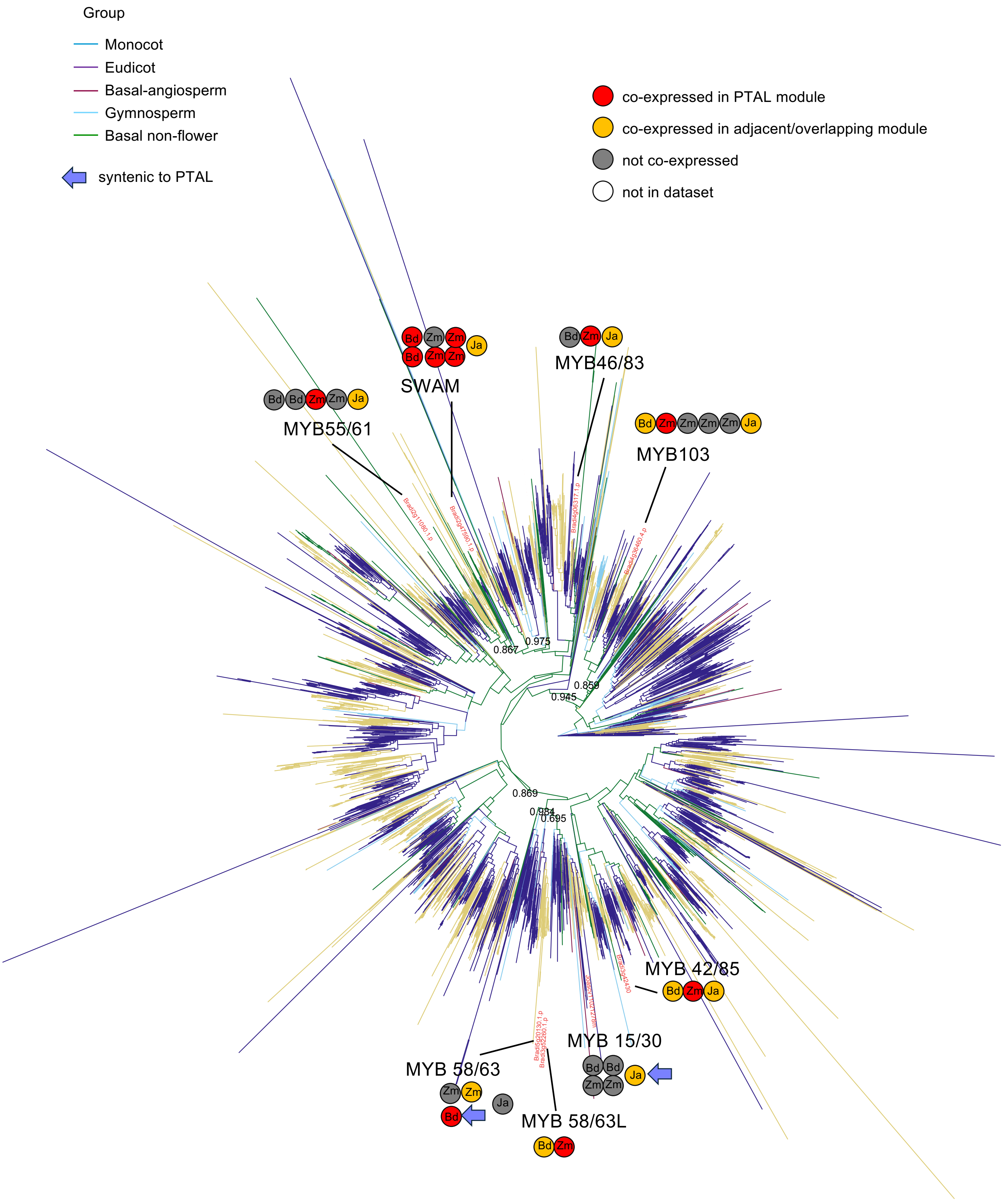


**Figure S15. MYB phylogeny shows changes and conservation of co-expression between *J. ascendens* and grass isoforms in different clades.**

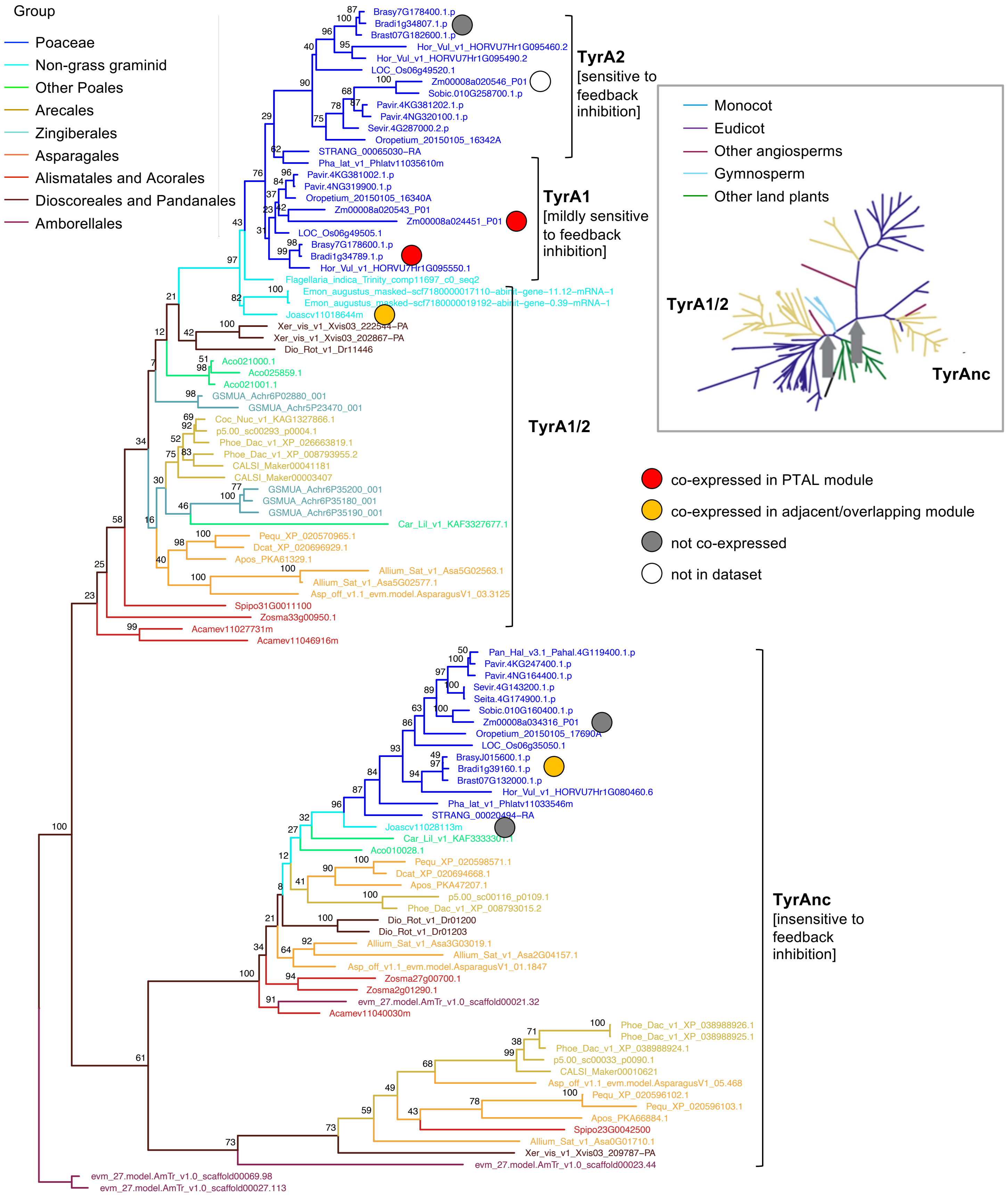


**Figure S16. TyrA phylogeny shows the maintenance of co-expression between *J. ascendens* and grass isoforms in one clade, and expansion of co-expression in grasses in another clade. The main tree is built using monocot species while the inset tree is built using species from all plant clades. The grey arrow in the inset tree shows where the base of the monocot tree starts.**

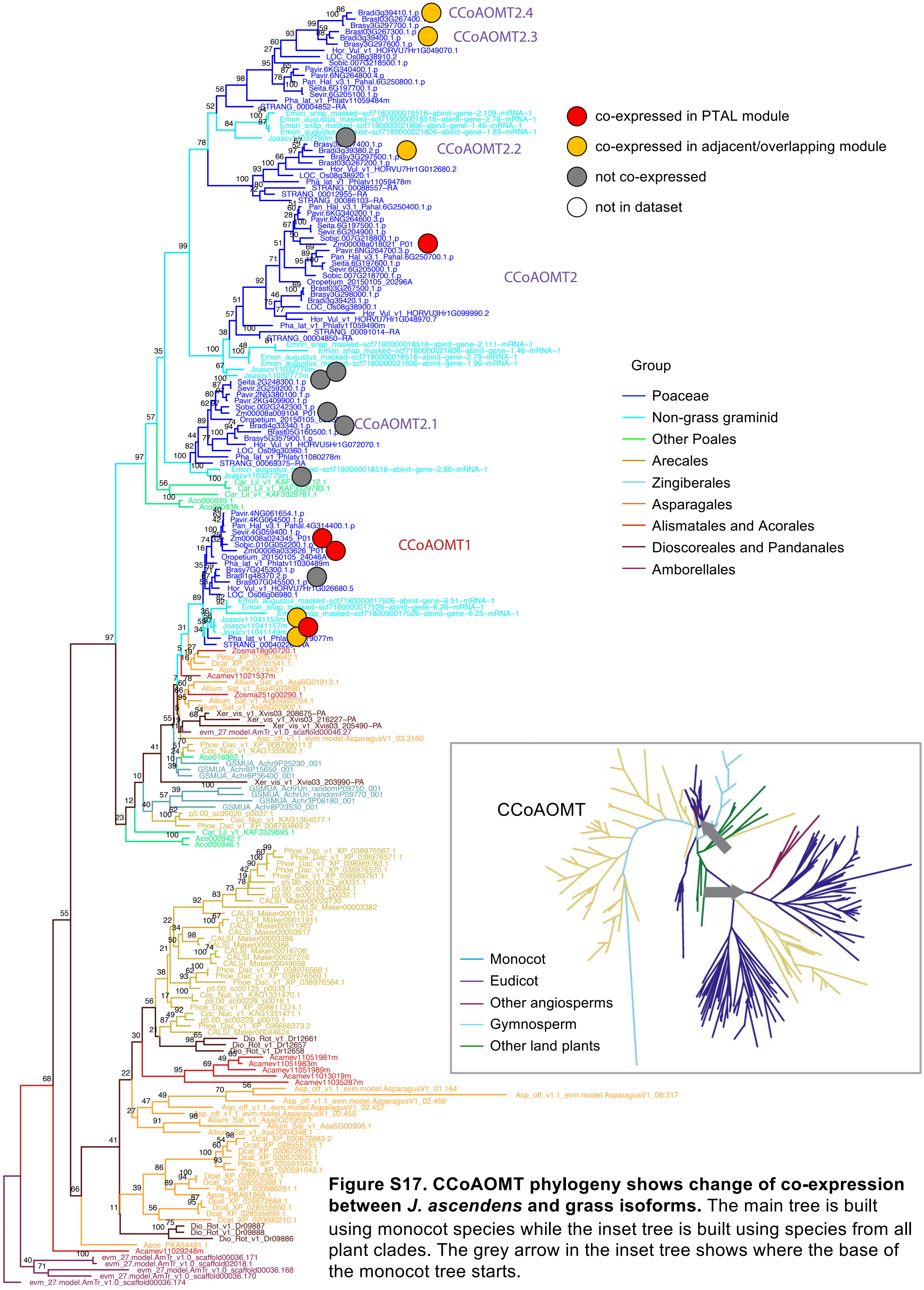


**Figure S17. CCoAOMT phylogeny shows change of co-expression between *J. ascendens* and grass isoforms.** The main tree is built using monocot species while the inset tree is built using species from all plant clades. The grey arrow in the inset tree shows where the base of the monocot tree starts.

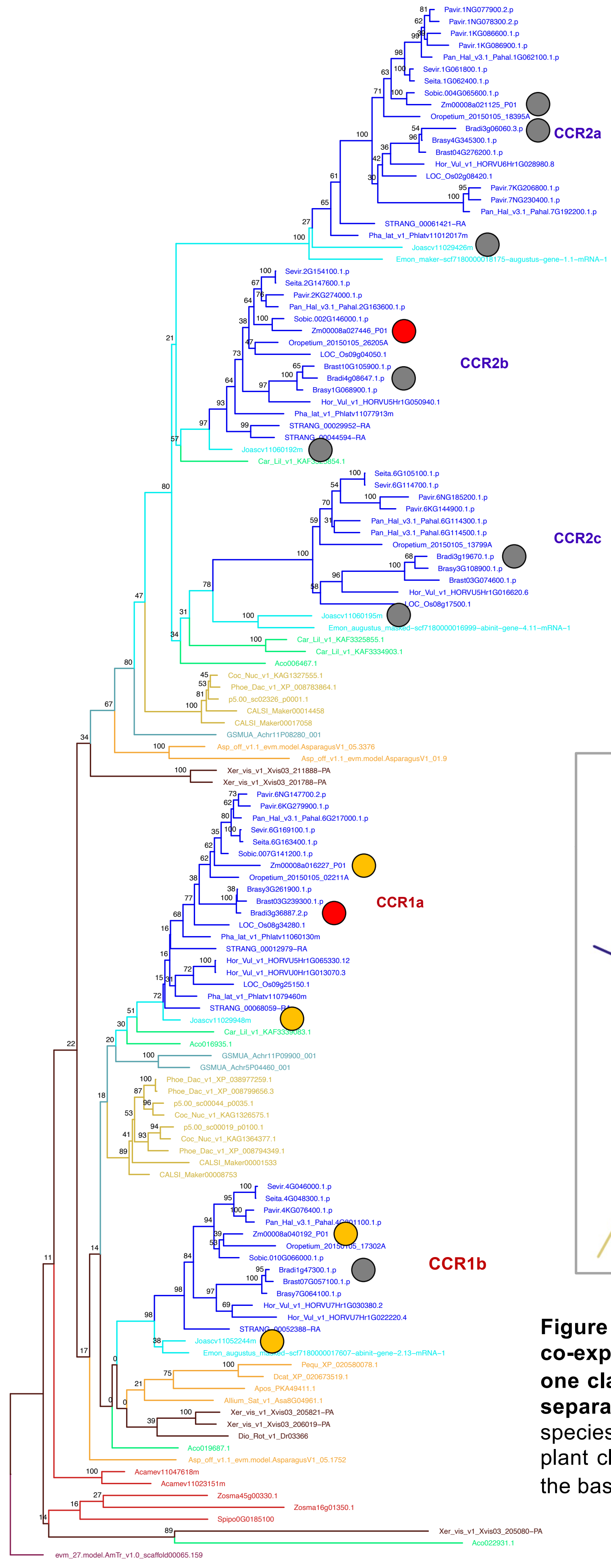


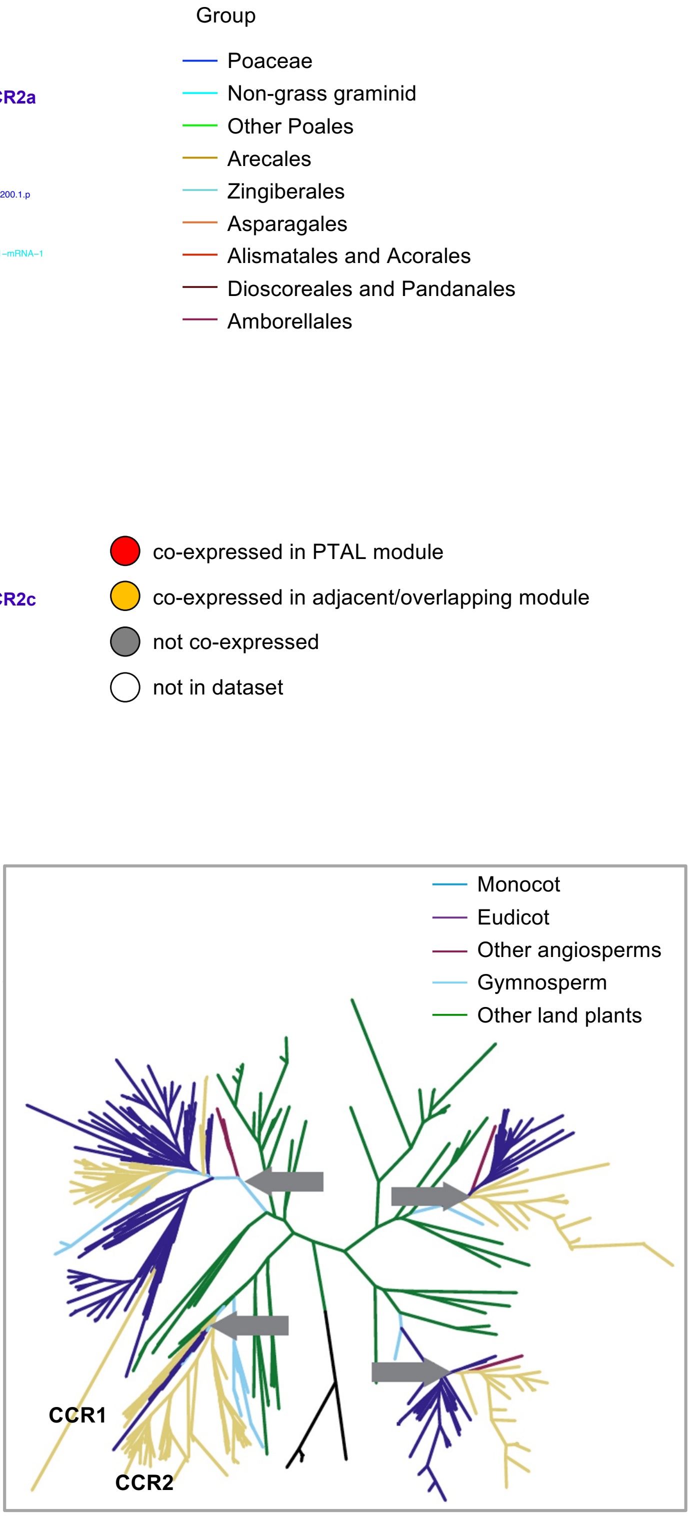


**Figure S18. CCR phylogeny shows the conservation of co-expression between *J. ascendens* and grasses in one clade and the gain of co-expression in grasses in a separate clade.** The main tree is built using monocot species while the inset tree is built using species from all plant clades. The grey arrow in the inset tree shows where the base of the monocot tree starts.

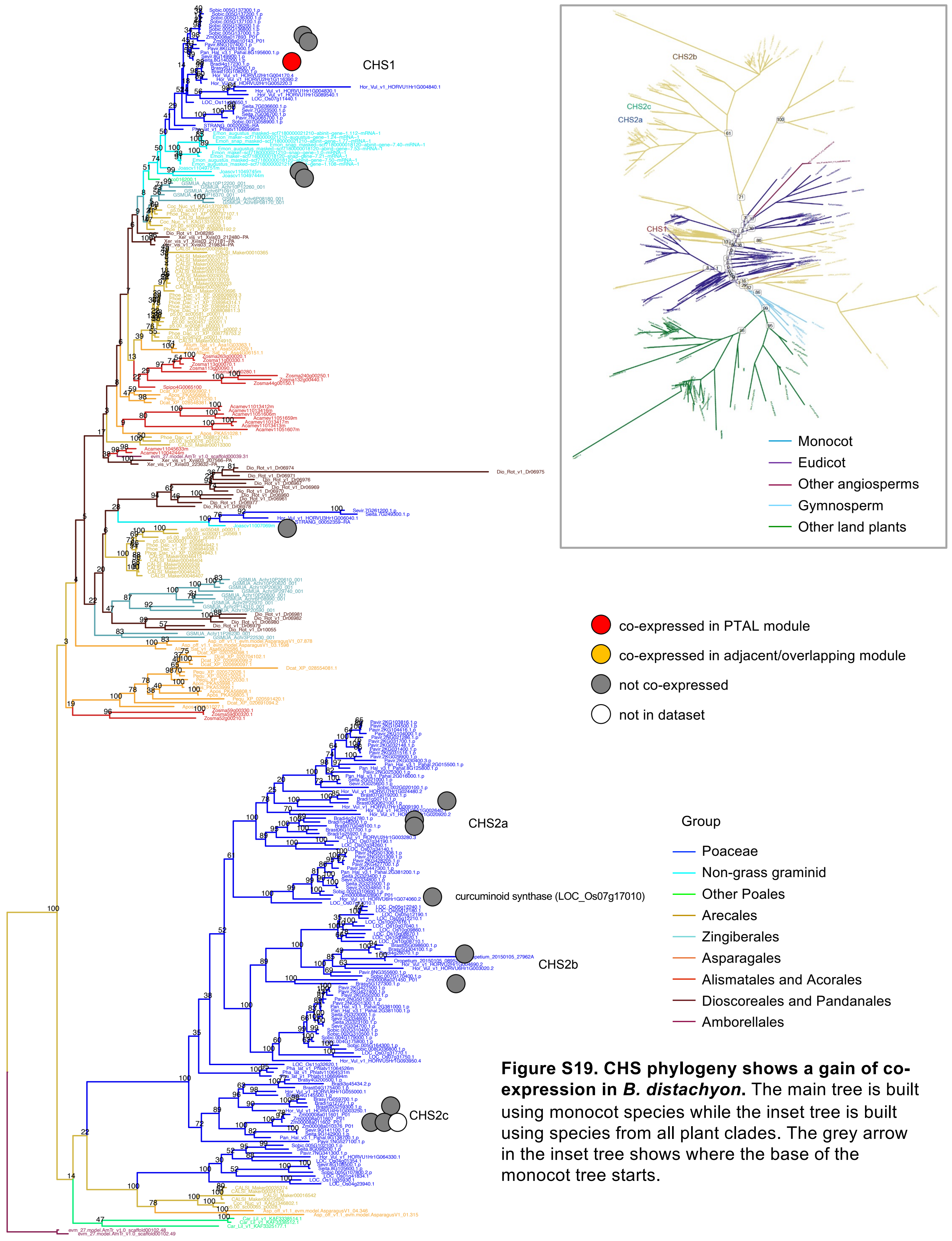


**Figure S19. CHS phylogeny shows a gain of co-expression in *B. distachyon*.** The main tree is built using monocot species while the inset tree is built using species from all plant clades. The grey arrow in the inset tree shows where the base of the monocot tree starts.

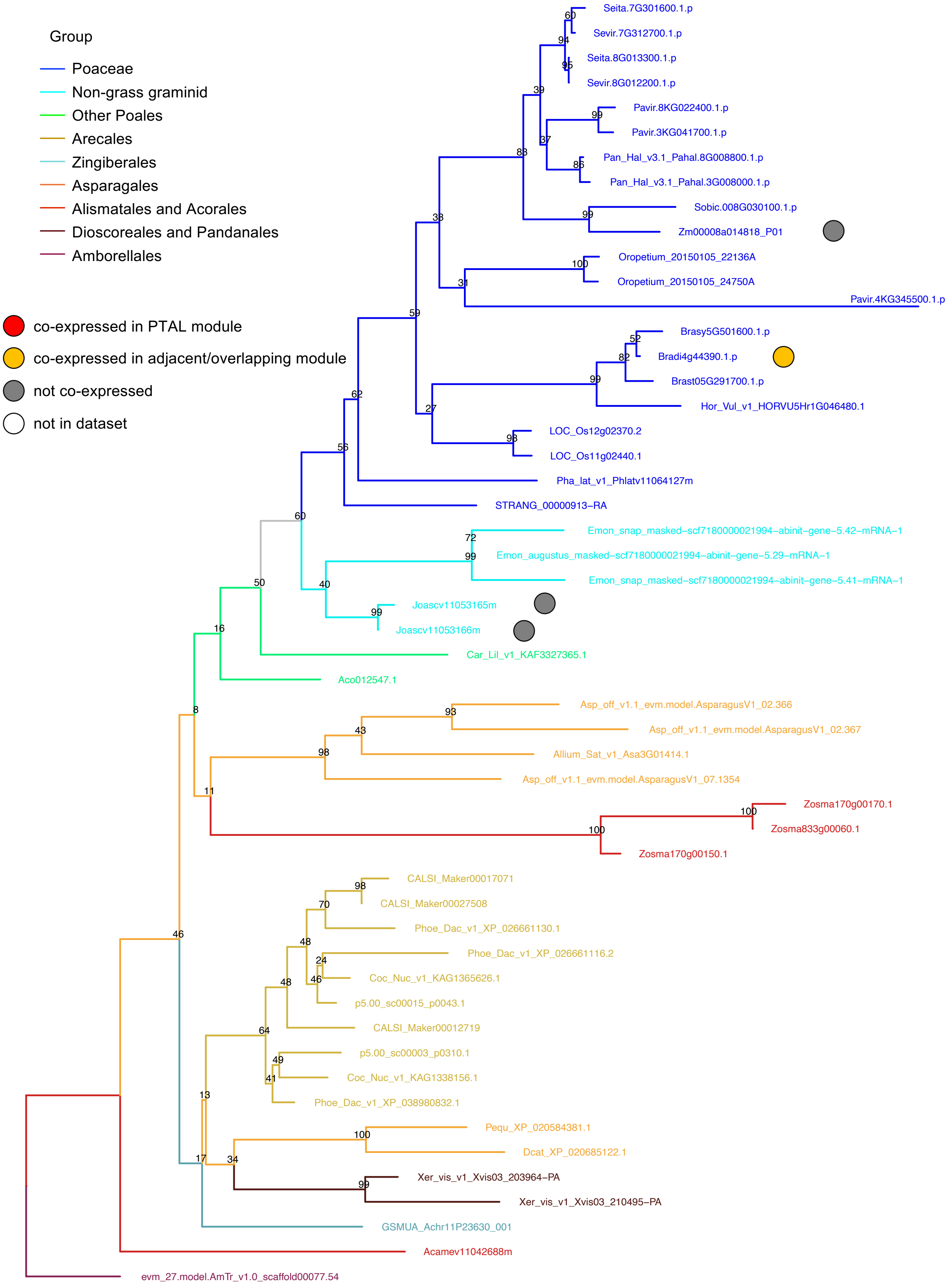


**Figure S20. CHIL phylogeny shows the expansion of co-expression in *B. distachyon* enzymes from *J. ascendens*. The tree is built using monocot species.**

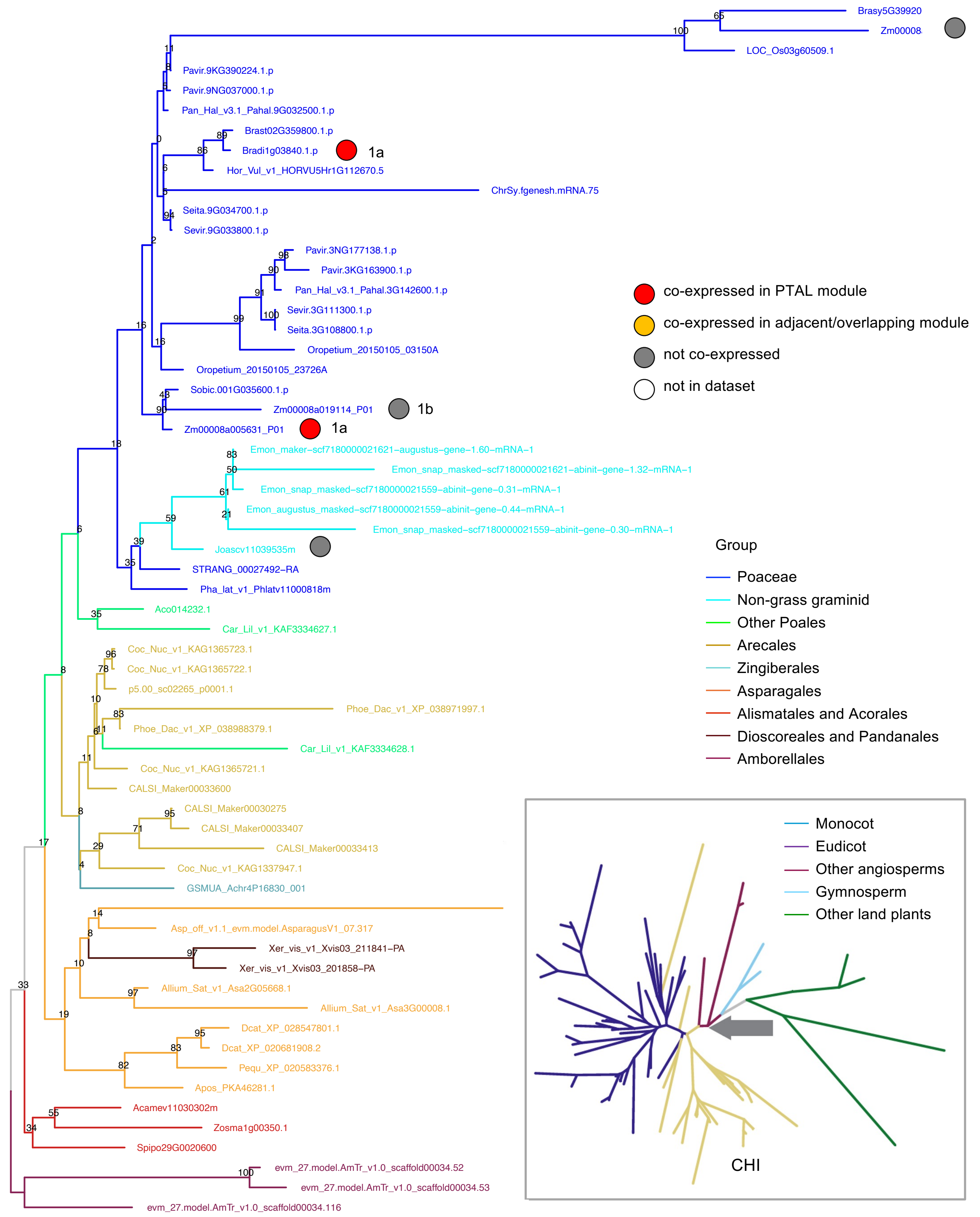


**Figure S21. CHI phylogeny shows a gain of co-expression in grasses.** The main tree is built using monocot species while the inset tree is built using species from all plant clades. The grey arrow in the inset tree shows where the base of the monocot tree starts.

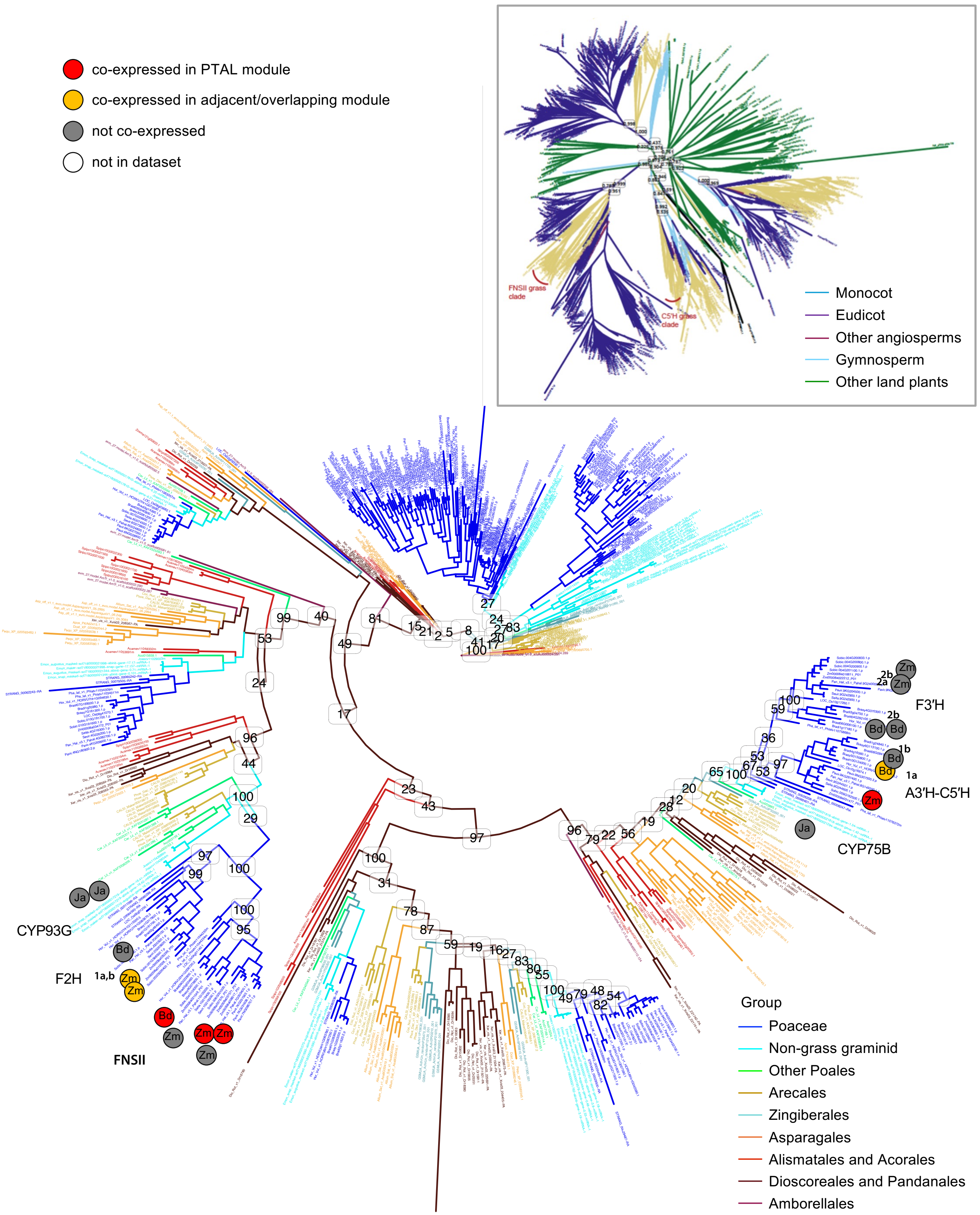


**Figure S22. CYP450 phylogeny shows the expansion of co-expression for two clades of tricin-producing enzymes from *J. ascendens* to grasses.** The main tree is built using monocot species while the inset tree is built using species from all plant clades.

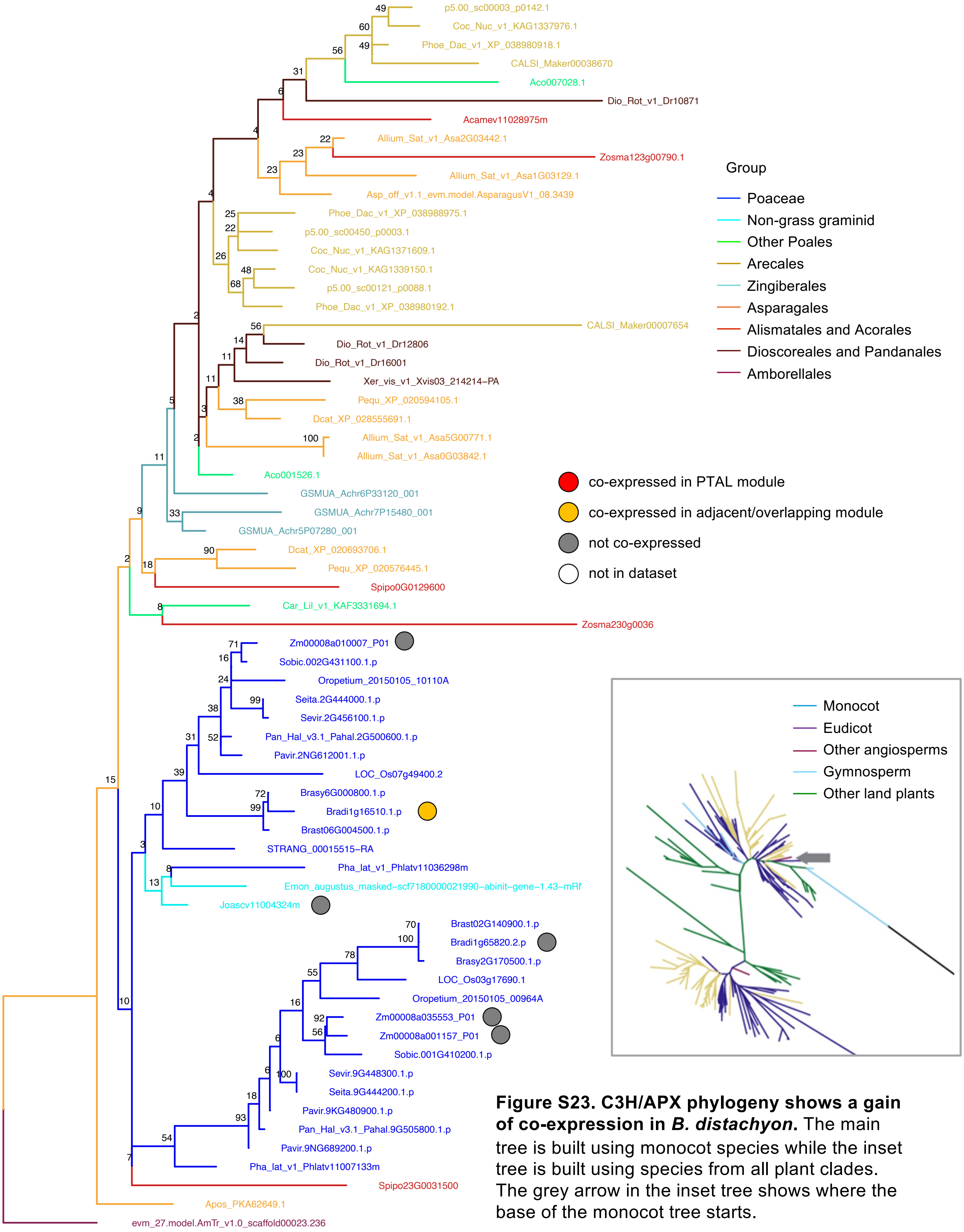


**Figure S23. C3H/APX phylogeny shows a gain of co-expression in *B. distachyon*.** The main tree is built using monocot species while the inset tree is built using species from all plant clades. The grey arrow in the inset tree shows where the base of the monocot tree starts.

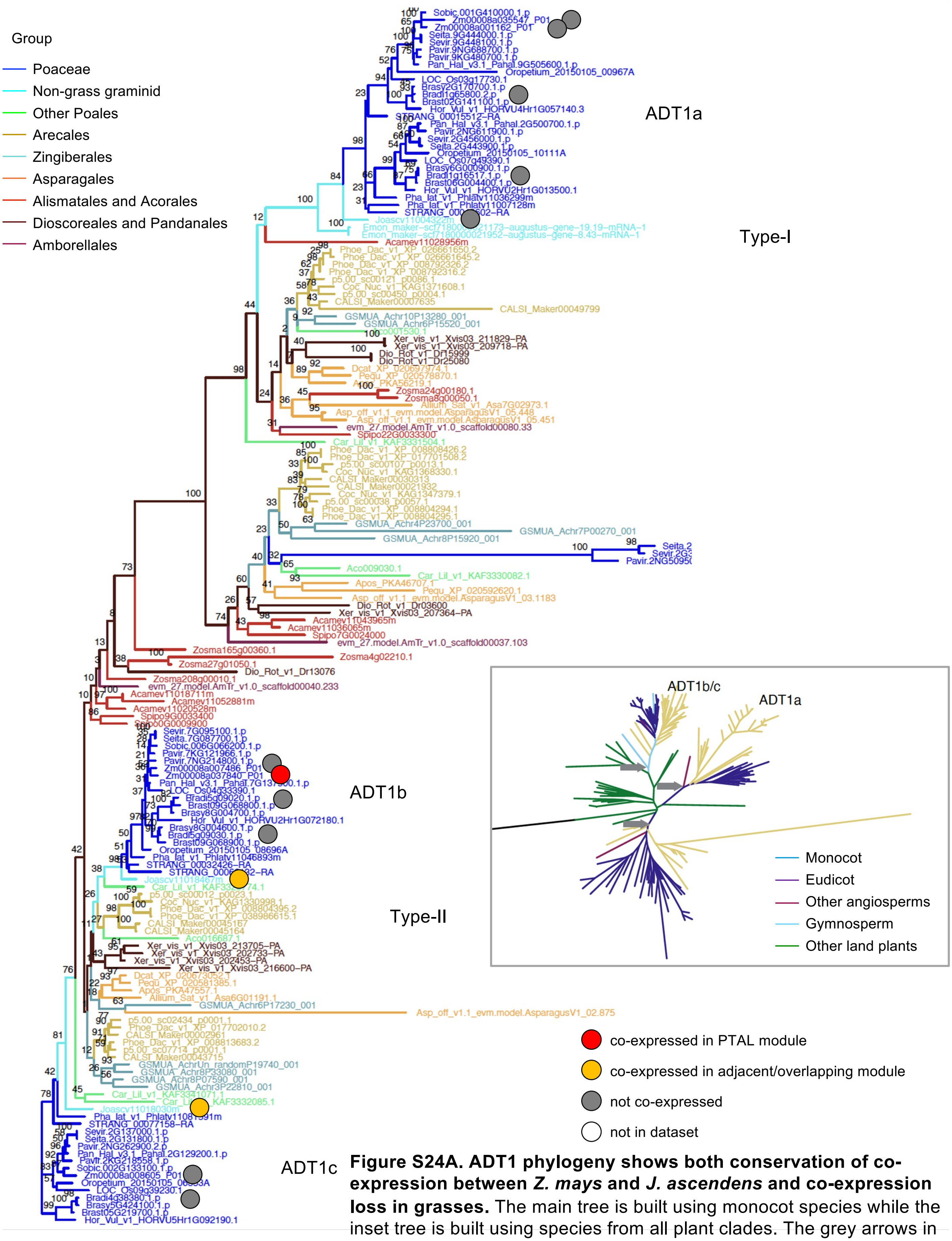


**Figure S24A. ADT1 phylogeny shows both conservation of co-expression between *Z. mays* and *J. ascendens* and co-expression loss in grasses.** The main tree is built using monocot species while the inset tree is built using species from all plant clades. The grey arrows in the inset tree shows where the base of the monocot tree starts.

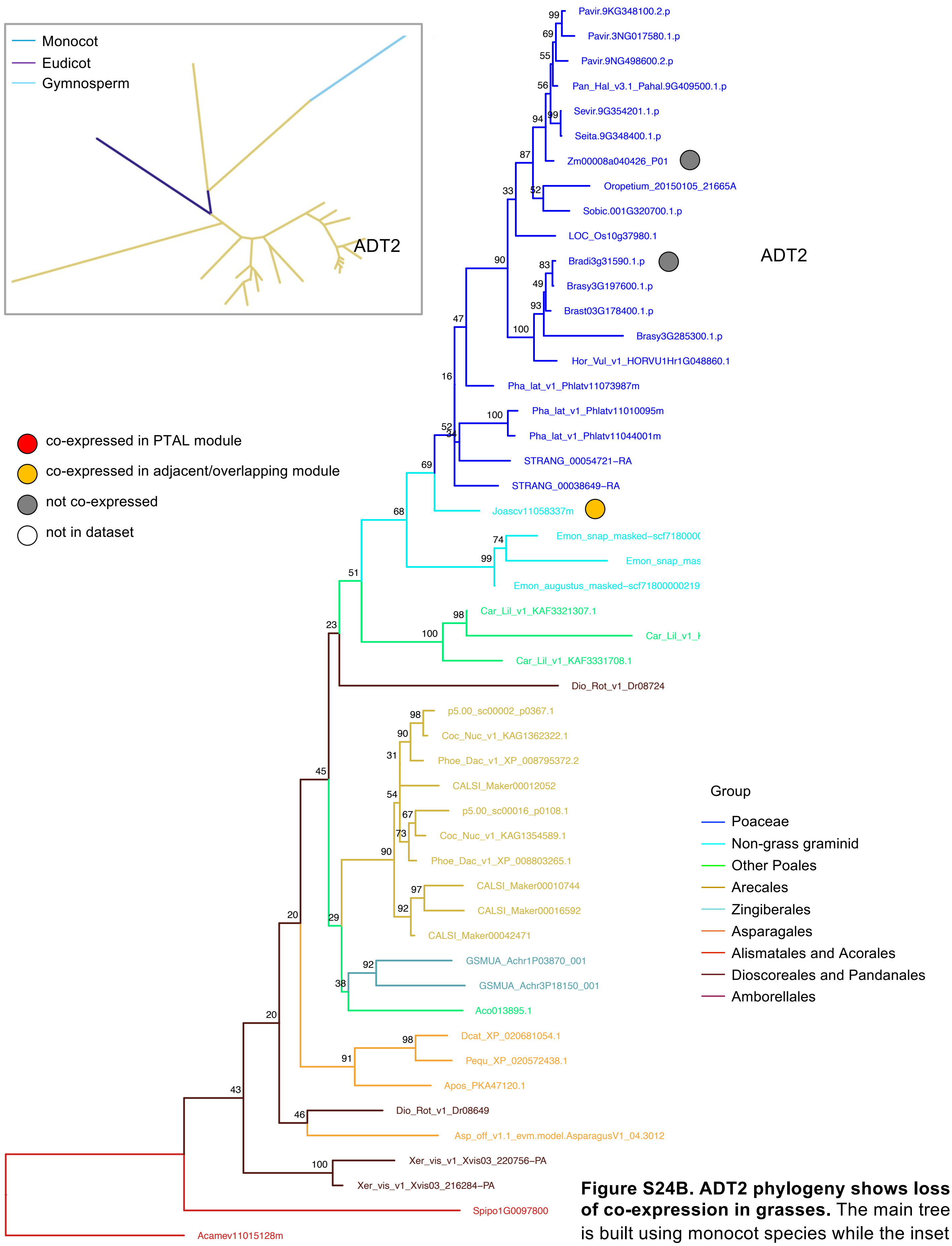


**Figure S24B. ADT2 phylogeny shows loss of co-expression in grasses.** The main tree is built using monocot species while the inset tree is built using species from all plant clades.

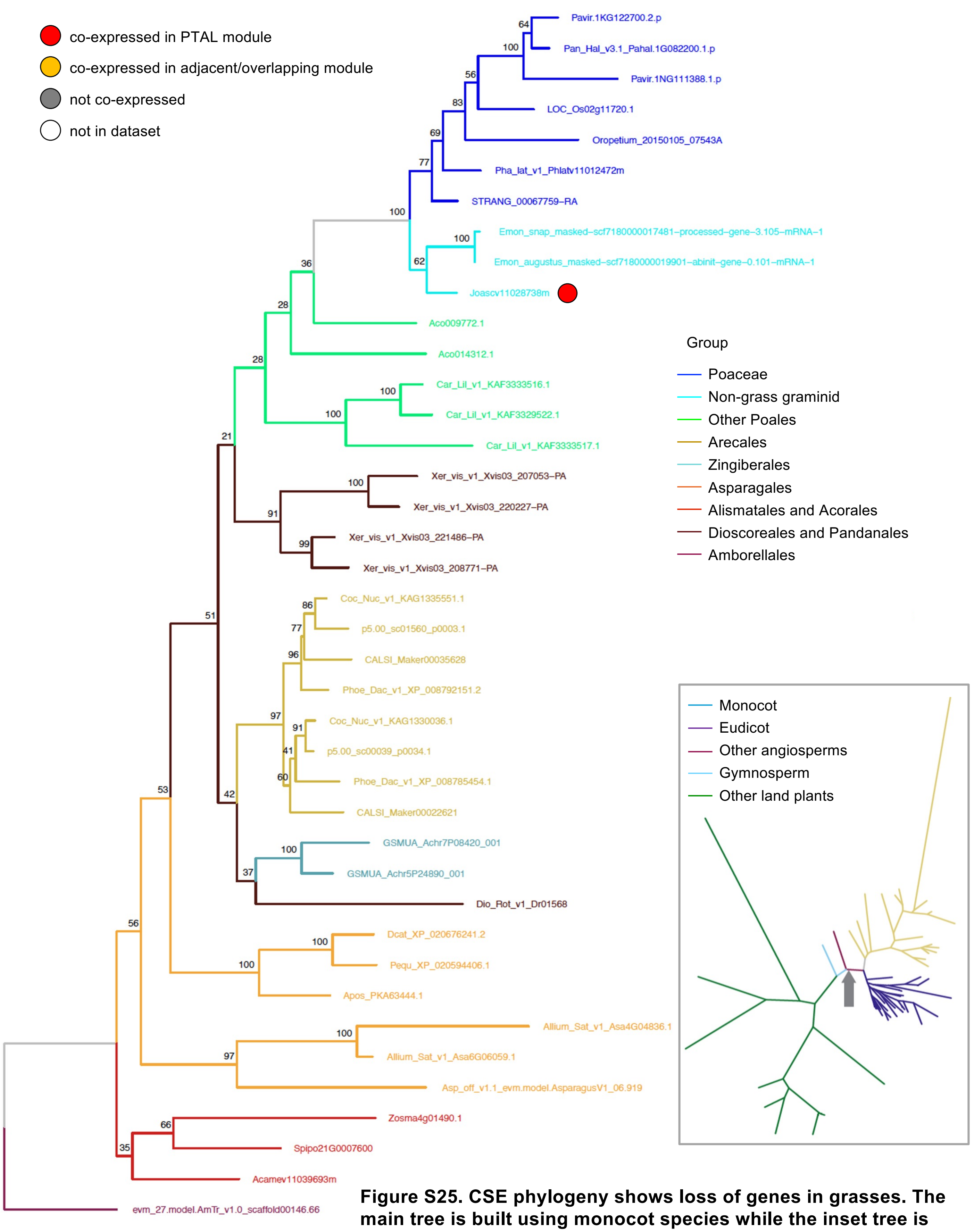


**Figure S25. CSE phylogeny shows loss of genes in grasses. The main tree is built using monocot species while the inset tree is built using species from all plant clades.** The grey arrow in the inset tree shows where the base of the monocot tree starts.

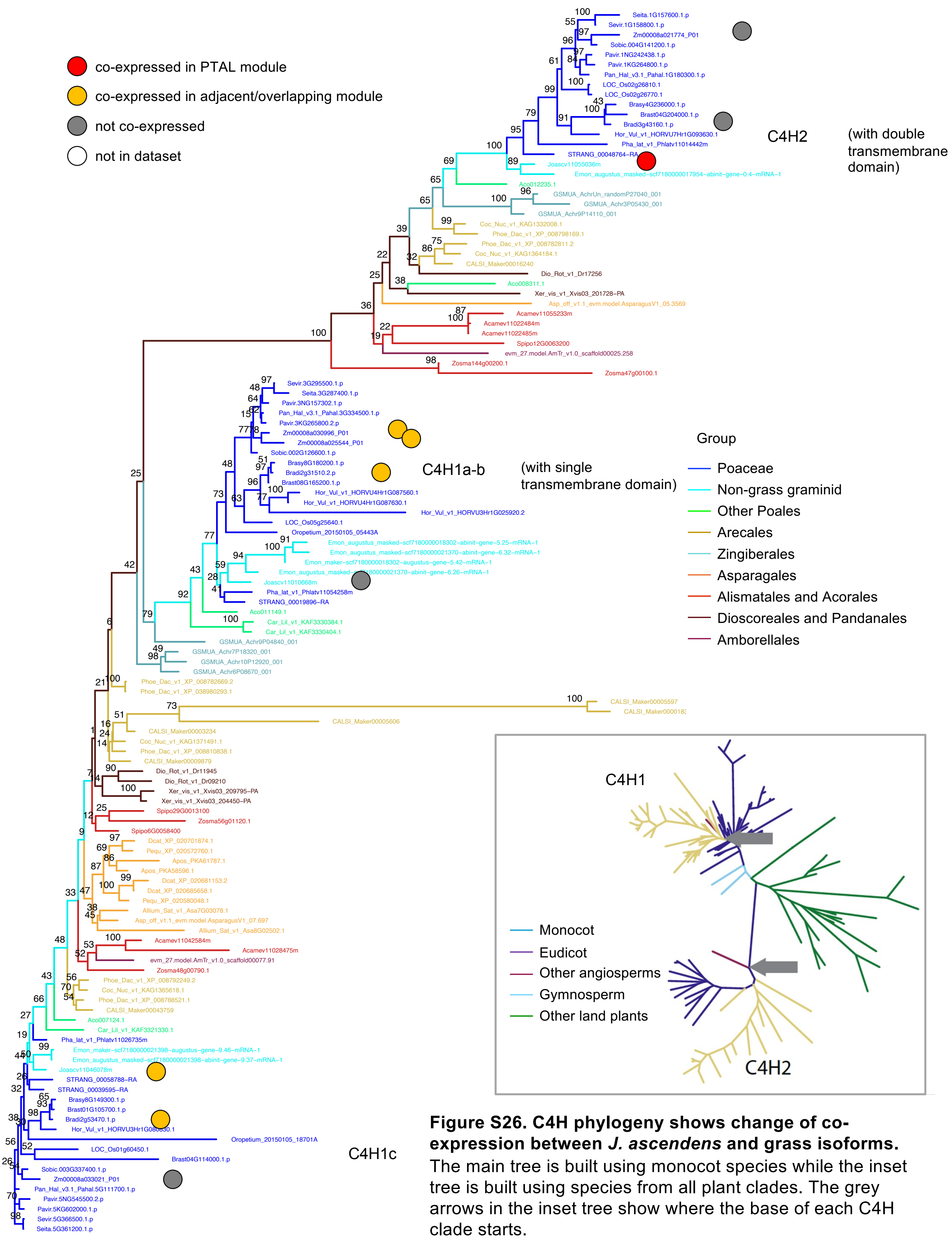


**Figure S26. C4H phylogeny shows change of co-expression between *J. ascendens* and grass isoforms.** The main tree is built using monocot species while the inset tree is built using species from all plant clades. The grey arrows in the inset tree show where the base of each C4H clade starts.

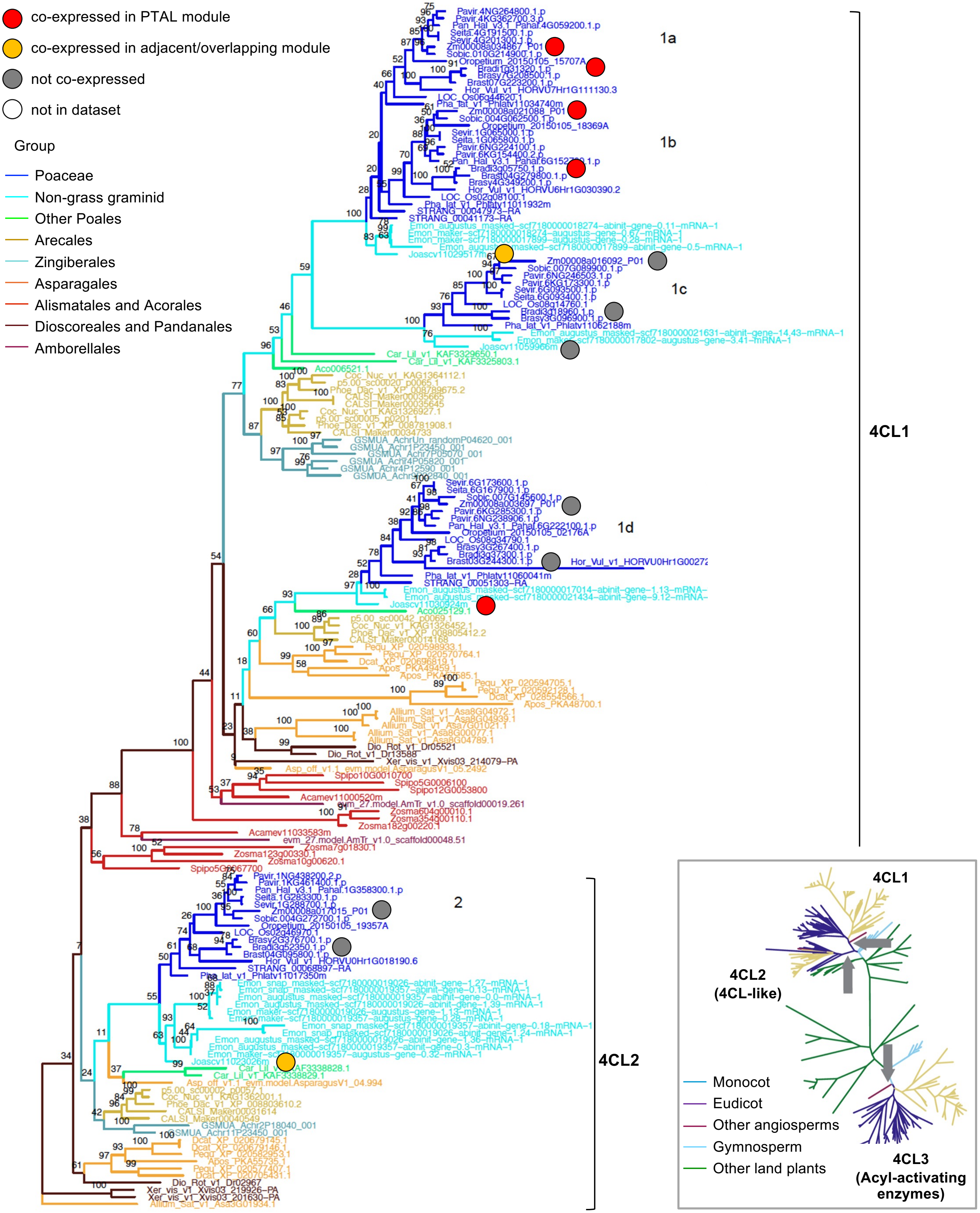


**Figure S27A. 4CL1-2 phylogeny shows conservation of co-expression between grasses and *J. ascendens* as well as a loss of co-expression in grasses.** Co-expression of isoform 4CL1a/b is maintained between grasses and *J. ascendens*, but isoforms 4CL1d and 4CL2 were co-expressed only in *J. ascendens*. The main tree is built using monocot species while the inset tree is built using species from all plant clades. The grey arrows in the inset tree show where the base of each 4CL clade starts.

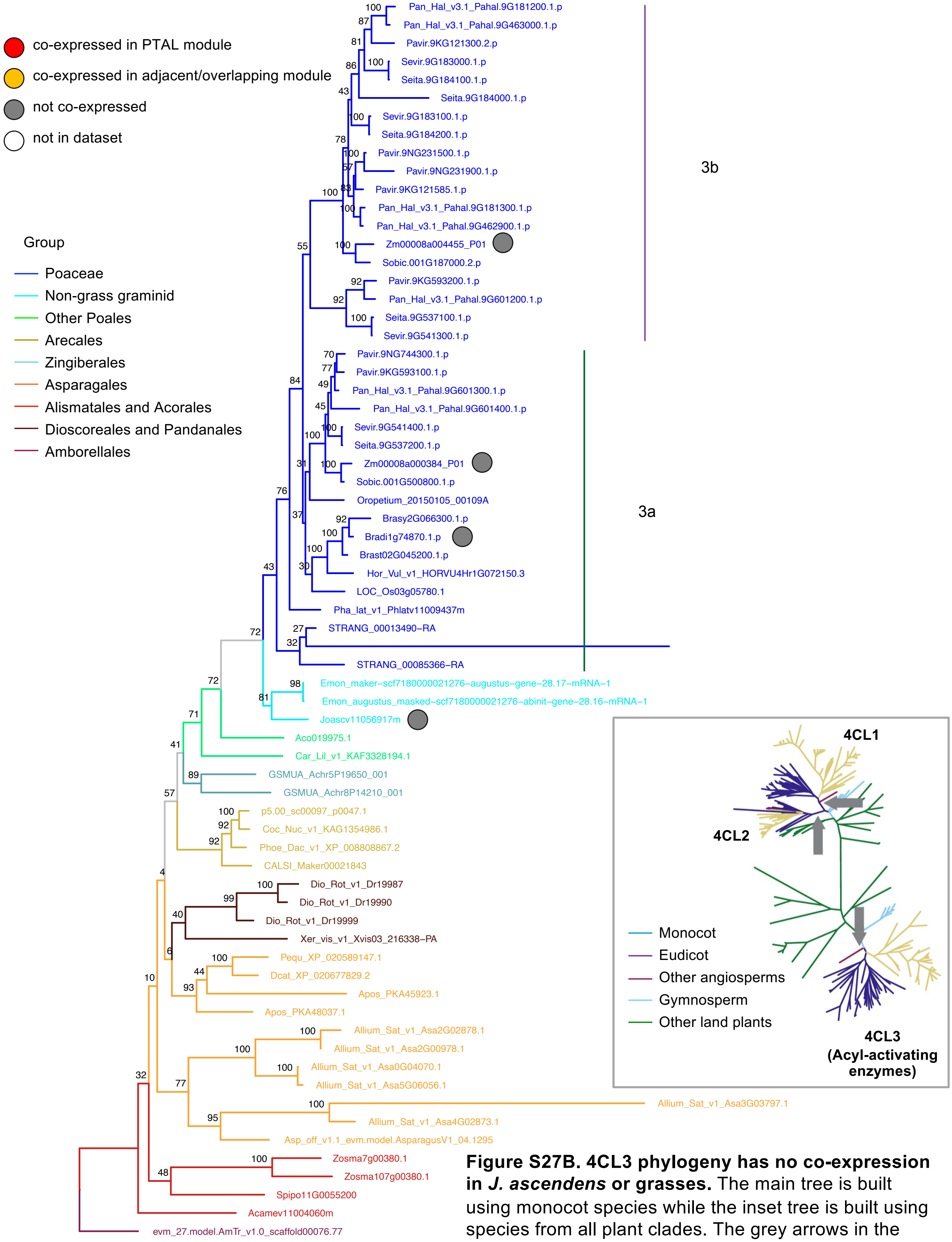


**Figure S27B. 4CL3 phylogeny has no co-expression in *J. ascendens* or grasses.** The main tree is built using monocot species while the inset tree is built using species from all plant clades. The grey arrows in the inset tree show where the base of each 4CL clade starts.

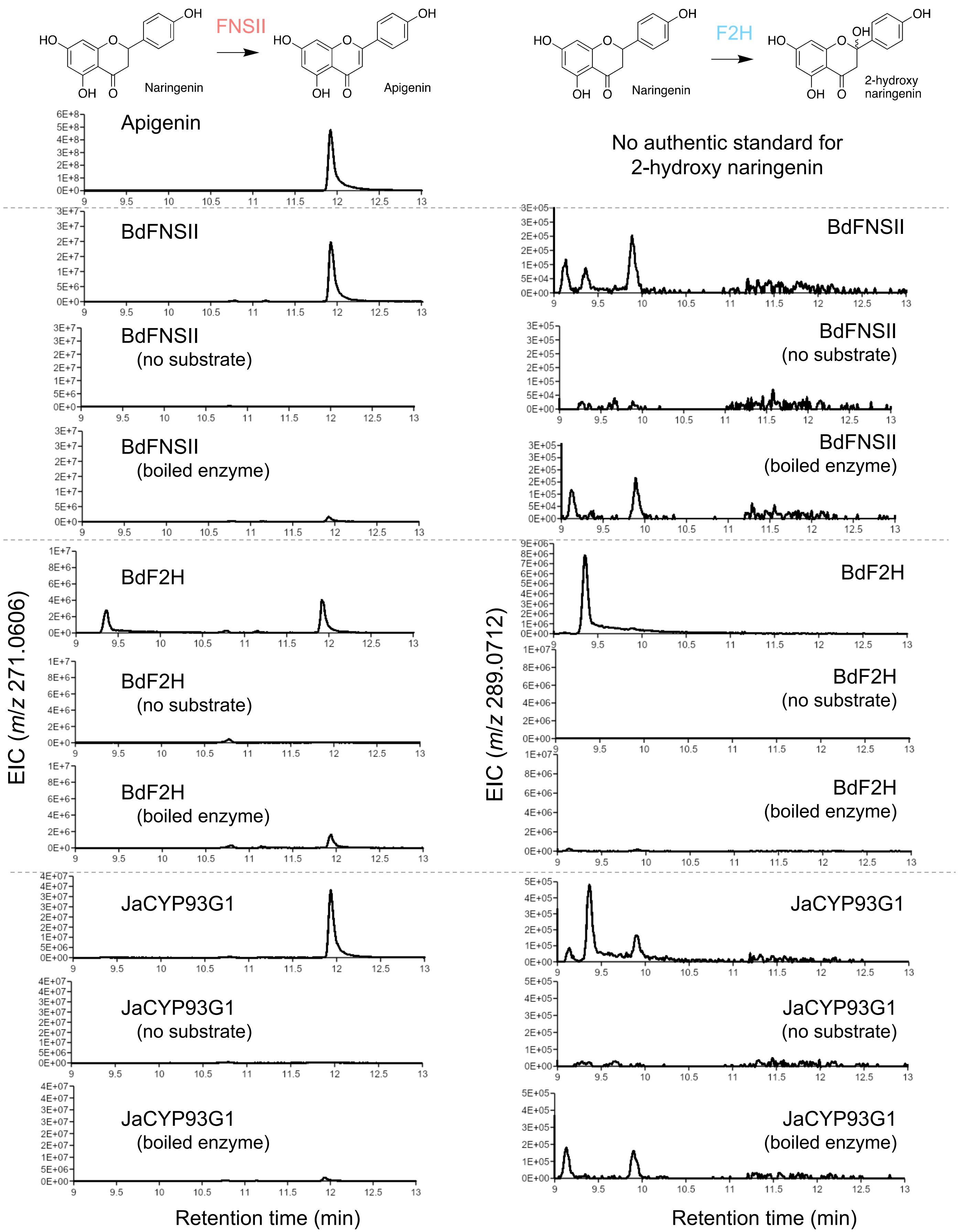


**Figure S28. FNSII and F2H activities of FNSII and F2H from *B. distachyon* and their closest homolog in *J. ascendens* (JaCYP93G1), showing the JaCYP93G1 has FNSII activity**

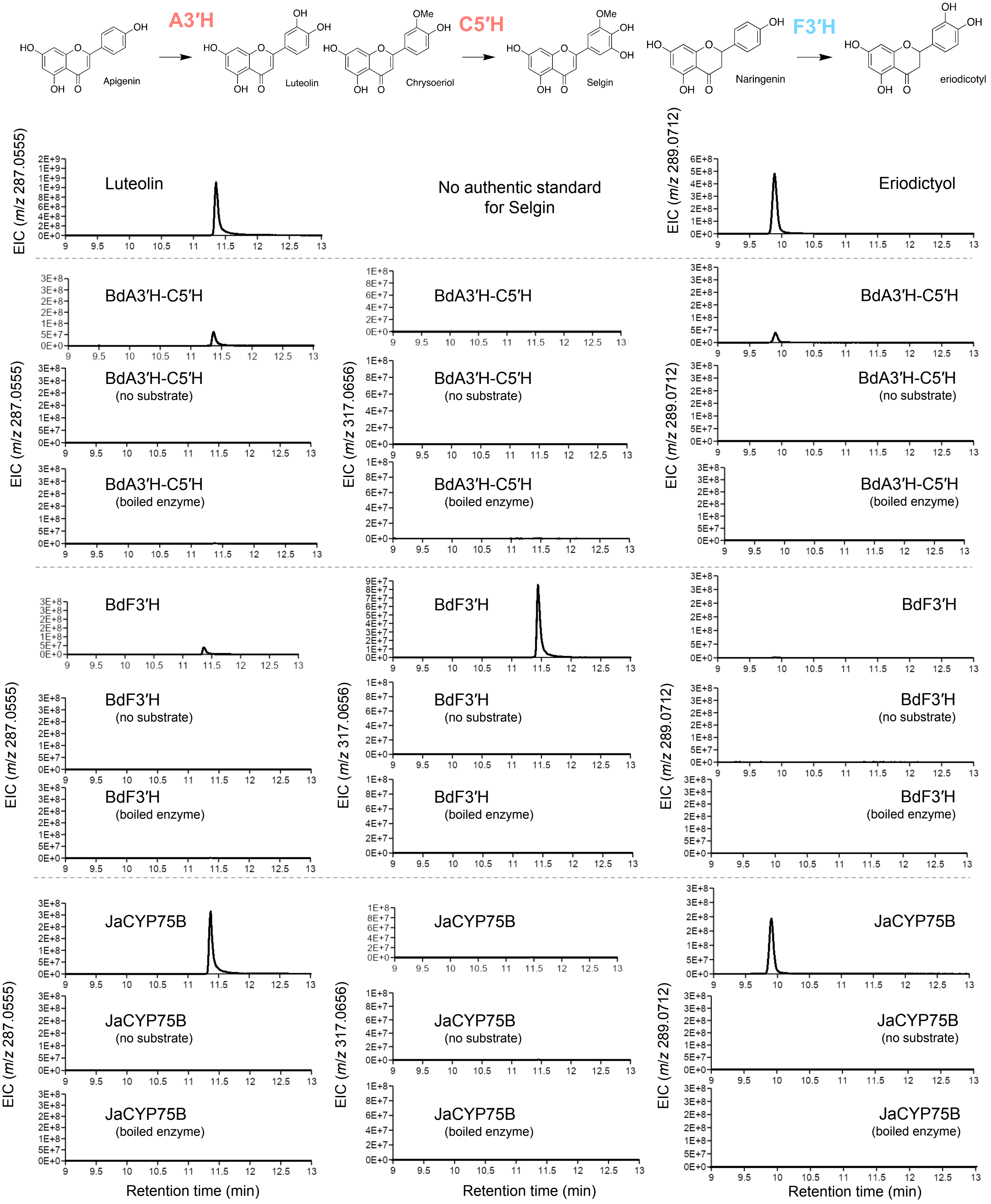


**Figure S29. A3′H, C5′H, and F3′H activities of A3′H-C5′H and F3′H from *B. distachyon* and their closest homolog in *J. ascendens* (JaCYP75B), showing the *J. ascendens* CYP75B enzyme lack C5′H activity**

**A.**

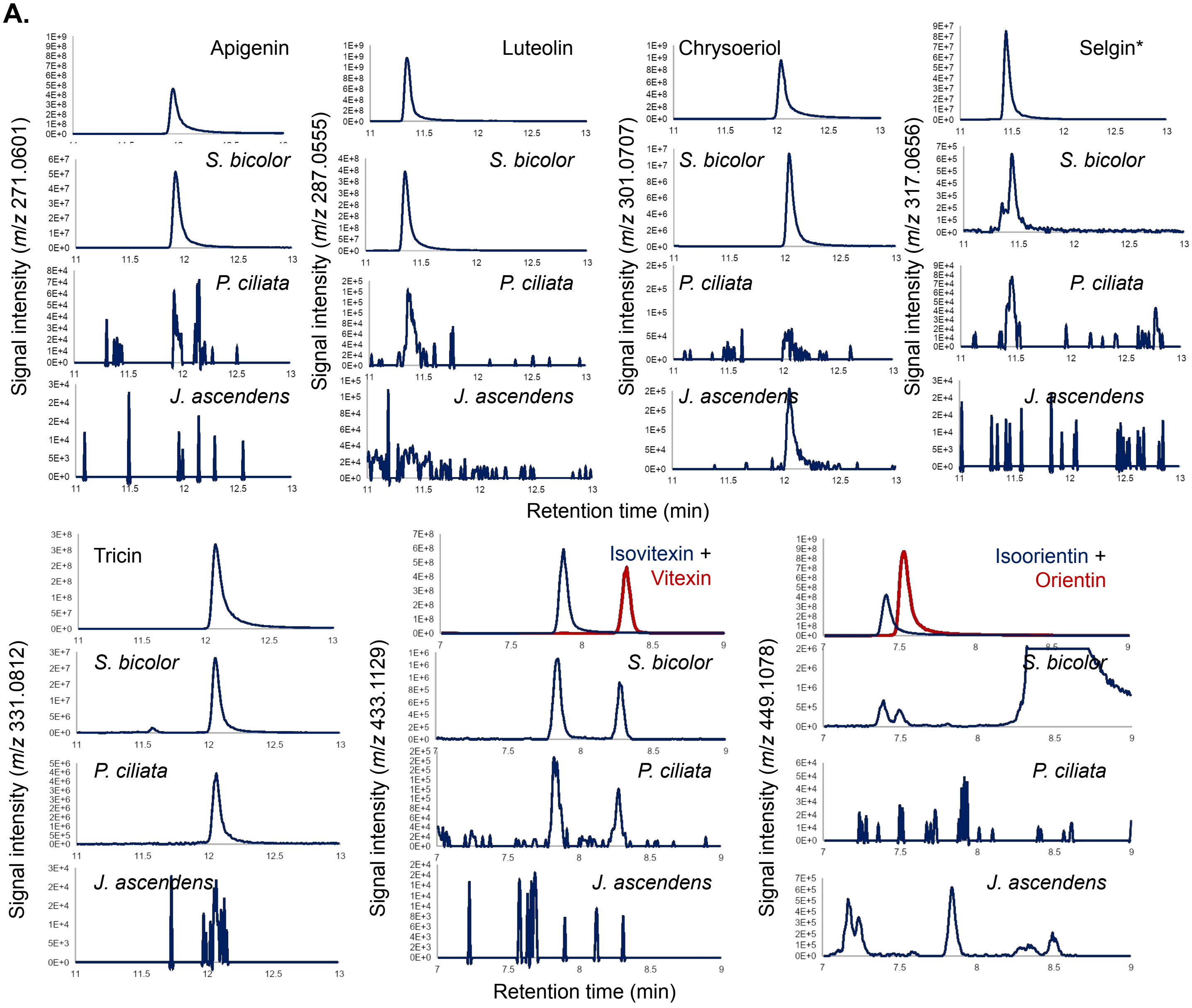


**B.**

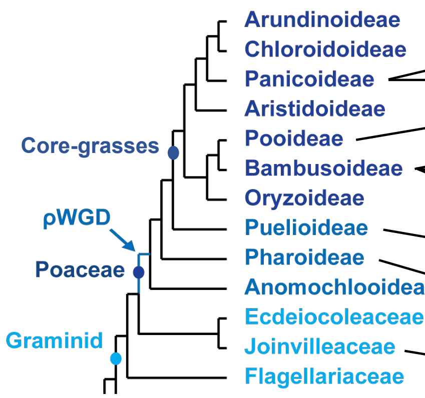


| | Tricin intermediates | | | | | Flavone *C*-glucosides | | |
|---|---|---|---|---|---|---|---|---|
| | Apigenin | Luteolin | Chrysoeriol | Selgin | Tricin | Vitexin | Isovitexin | Orientin + Isoorientin |
| *S. bicolor* | 988 | 6132 | 203 | 15 | 402 | 21 | 12 | 15 |
| *S. virtdis* | 9 | 80 | 15 | 1 | 51 | 263 | 7 | 74 |
| *B. distachyon* | 0 | 0 | 8 | 2 | 249 | 4 | 28 | 312 |
| *S. tsuboiana* | 2 | 6 | 10 | 3 | 1612 | 1 | 2 | 14 |
| *P. pygmaeus* | 1 | 3 | 7 | 1 | 559 | 1 | 0 | 7 |
| *P. aurea* | 9 | 3 | 22 | 2 | 688 | 16 | 7 | 25 |
| *P. ciliata* | 1 | 3 | 1 | 1 | 82 | 3 | 2 | 0 |
| *P. lappulaceus* | 0 | 0 | 4 | 0 | 1 | 0 | 0 | 0 |
| *S. angustofolia* | 0 | 0 | 5 | 0 | 1 | 4 | 0 | 1 |
| *J. ascendens* | 0 | 0 | 6 | 0 | 0 | 0 | 0 | 0 |

high / low

(Peak area /dry weight)

**Figure S30. Accumulation of tricin intermediates and flavone *C*-glucosides determined by LC-MS analysis of the MeOH extract. Chromatogram of each compound for the selected species (A) and normalized LC-MS peak intensity (B) is shown.**

*Selgin standard peak was from C5′H enzyme reaction product.

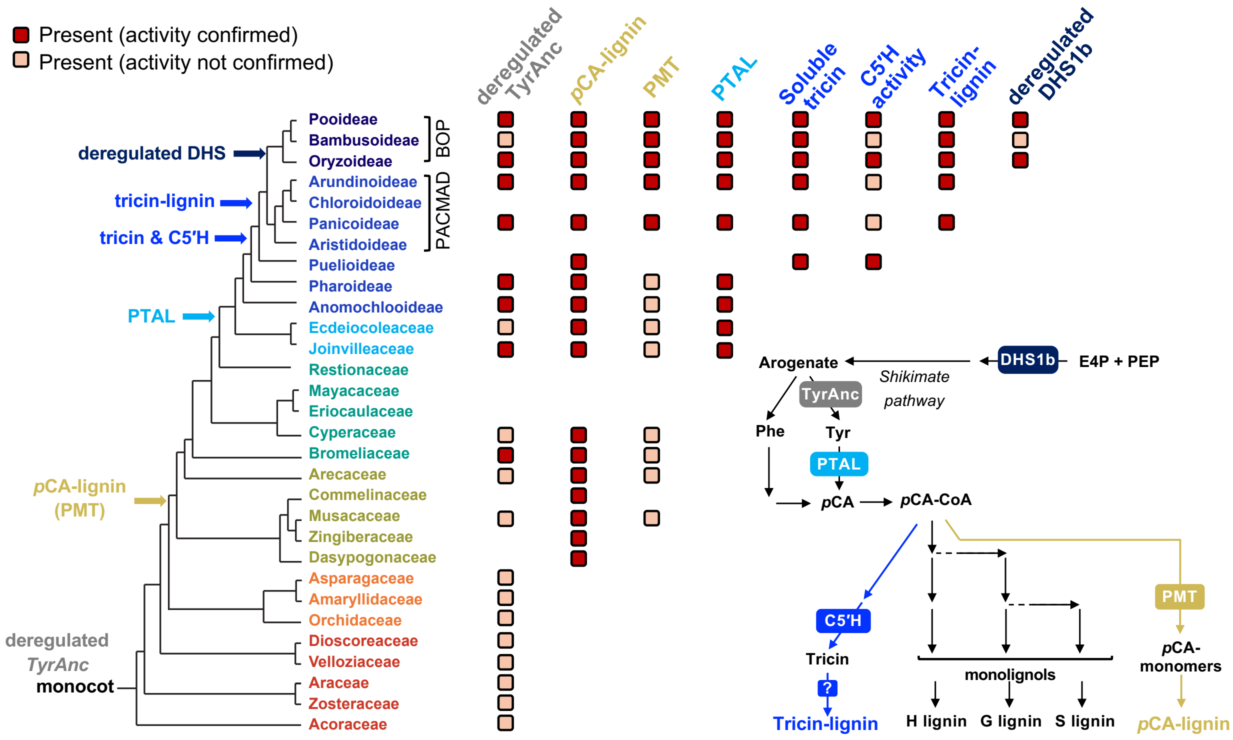


**Figure S31. Grass lignin network evolved in a stepwise manner, in which *p*CA-lignin subunits occurred first, followed by tyrosine-derived lignin biosynthesis and then addition of tricin into the cell wall lignin.** Monocot and Poales ancestors already had dregulated TyrAnc and PMT, which produce tyrosine and *p*CA-lignin, respectively. At the divergence of grasses and *J. ascendens*, PTAL emerged and generated two entry pathways for lignin biosynthesis. After the divergence of *Pharus* and *Puelia* C5'H activity emerged within the grass CYP75B family, leading to the acquisition of the tricin biosynthetic pathway, with soluble tricin later being incorporated into the cell wall lignin within core-grasses. The evolution of PTAL likely elevated the demand of tyrosine, which further facilitated the de-regulation of TyrAnc and the emergence of deregulated DHS1b isoform, which increased the upstream shikimate pathway activity in BOP grasses. Dark red boxes denote the presence of confirmed enzyme activity or chemical accumulation, while light orange boxes indicate the presence of orthologous genes whose activity have not ben confirmed.

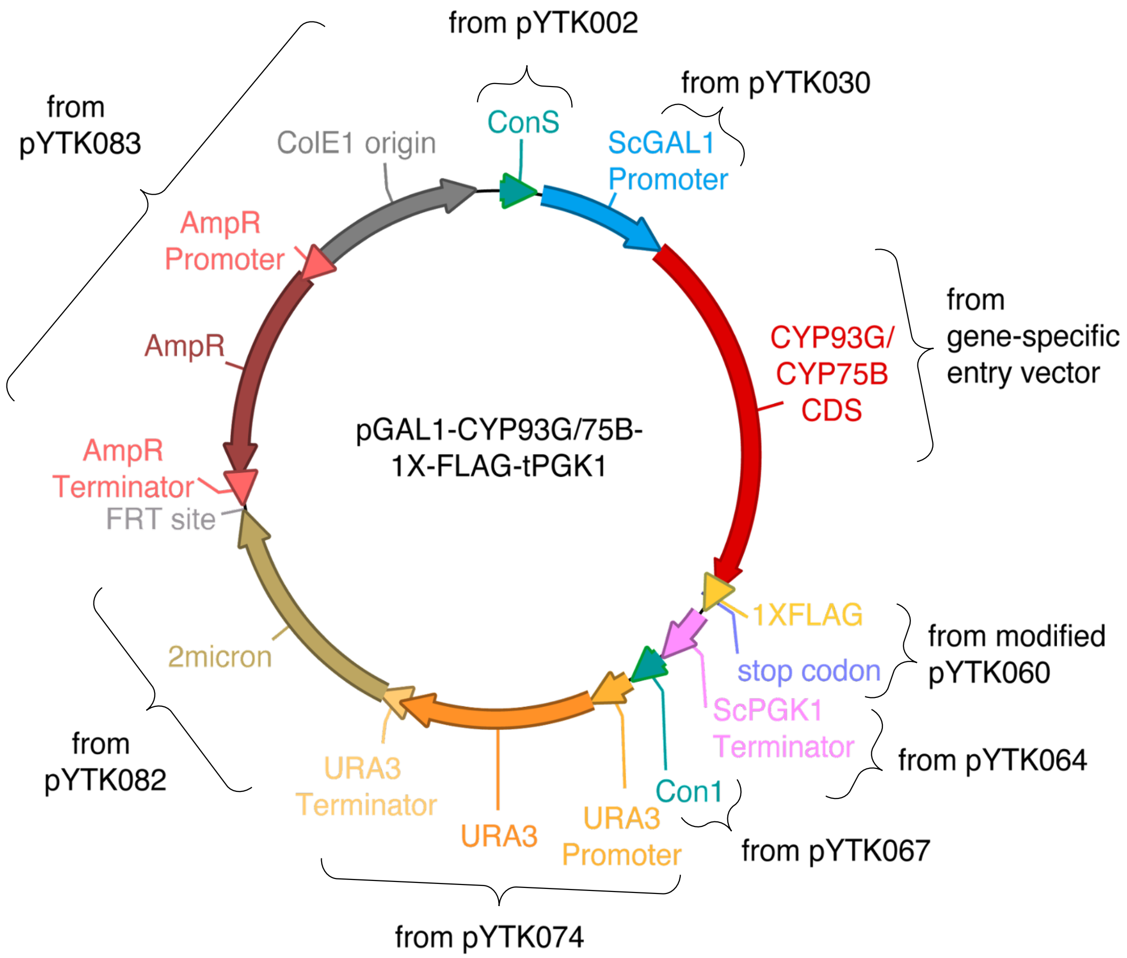


**Figure S32. Golden gate constructs for recombinant protein induction in *S. cerevisiae* WAT11**
